# Probing the energy landscape of α-Synuclein amyloid fibril formation by systematic K-to-Q mutagenesis

**DOI:** 10.1101/2025.11.01.685997

**Authors:** Antonin Kunka, Azad Farzadfard, Jacob Aunstrup Larsen, Federica Saraceno, Rasmus Krogh Norrild, Celia Fricke, Hossein Mohammad-Beigi, Ahmed Sadek, Jonas Folke, Susana Aznar, Alexander K. Buell

**Affiliations:** Department of Biotechnology and Biomedicine Technical University of Denmark Søltofts Plads, Building 227, 2800 Kgs. Lyngby, Denmark; Department of Biotechnology and Bioengineering Ecole polytechnique fédérale de Lausanne EPFL; Centre for Neuroscience and Stereology, Department of Neurology, Copenhagen University Hospital, Bispebjerg and Frederiksberg Hospital, Nielsine Nielsens Vej 6B, Entrance 11B, 2. Floor, 2400 Copenhagen, Denmark

**Keywords:** IDPs, polymorph, energy landscape, mutagenesis, Parkinson’s disease

## Abstract

The aggregation of natively disordered α-Synuclein (αSyn) into amyloid fibrils is a hallmark of Parkinson’s and other neurodegenerative diseases. Understanding αSyn’s pathological role remains a major challenge due to its complex, context-dependent energy landscape characterized by conformational plasticity and fibril polymorphism. Here, we present a systematic mutational analysis as a quantitative probe of the αSyn energy landscape, focusing on electrostatic contributions to key aggregation pathways. We engineered αSyn variants with one to eight lysine-to-glutamine substitutions and analyzed their aggregation under controlled conditions to delineate their effects on nucleation, elongation, seed amplification, fibril stability, and fibril polymorphism. We find that αSyn aggregation from a homogenous solution can be modelled well using global properties, including protein concentration, charge, and ionic strength. Microscopic pathways and the resulting fibril polymorphs are instead modulated by sequence-specific effects. We identify mutations of residues found in fibril cores as perturbations that significantly modify the αSyn free energy landscape, creating pathways and energy minima not accessible to the WT under the same experimental conditions. In contrast, mutations outside of the fibril core affect the magnitude of the relevant energy barriers whilst overall maintaining a WT-like free energy landscape. Our work outlines a scalable, quantitative framework that increases the informational output of the mutational studies of αSyn using conventional assays. The approach can be extended by incorporating additional mutational and functional data to deepen our understanding of αSyn’s energy landscape and its role in health and disease.

## Introduction

α-Synuclein (αSyn) is a small 14.5 kDa protein found in neuronal cells, as well as the peripheral nervous system, and red blood cells. (1–5) It plays diverse roles in synaptic vesicle trafficking and recycling, dopamine regulation, calcium signaling, mitochondrial function, and lipid metabolism. (6–12) Pathologically, αSyn is the principal component of intracellular inclusions that characterize synucleinopathies, including Parkinson’s disease (PD), Lewy body dementia (LBD), and multiple system atrophy (MSA). (13) Although it has been demonstrated that αSyn aggregates are toxic and propagate between cells in a prion-like manner, it remains unclear whether disease arises primarily from gain-of-function toxicity or loss of normal protein function. (14–18)

One major obstacle to resolving αSyn’s role in disease and developing effective therapies is the complexity of the protein itself. Even outside of its biological background, characterization of αSyn presents a multifaceted challenge due to its extreme conformational plasticity and context-dependent behaviour. Its ability to transition from its disordered monomeric state to various oligomeric or fibrillar forms, each potentially associated with distinct cellular outcomes, complicates both mechanistic understanding of the diseases and their therapeutic targeting. (16,18–22) This structural flexibility is also modulated by mutations, post-translational modifications (PTMs), differential expression, and cellular localization, further complicating the distinction between physiological and pathological states. (23–29)

The complex energy landscape of αSyn is often probed through a combination of advanced biophysical methods (e.g., NMR), which provide invaluable resolution but are often not scalable and/or are technically demanding. (30–34) A compelling approach is to instead exploit mutational analysis which is well established for studying energetics of folding, binding, and stability of globular proteins.(35–38) Studies involving familiar mutants (39,40), N- and C-terminal truncations (41–43), PTMs and their mimetics (44–48), and large-scale variant analyses (49,50) have collectively highlighted regions critical for αSyn aggregation, suggesting that mutational scanning can indirectly map features of the underlying energy landscape, analogous to what is possible for folded proteins. Despite the wealth of insightful results from these and other studies (51–58), our ability to predict how individual substitutions alter specific assembly pathways of αSyn (e.g., nucleation, elongation, seed amplification) within specific biological contexts remains limited.

Mutational studies of αSyn generally investigate three classes of protein variants: (i) physiologically or pathologically relevant variants (e.g., disease-associated familial mutants), (ii) targeted sequence perturbations designed to test specific hypotheses or probe mechanisms (e.g., Φ-value analysis) (59), and (iii) random variants used for unbiased searches that typically require large libraries to be informative. The first class provides direct insight into pathogenic mechanisms by revealing how disease mutations reshape the energy landscape and assembly behaviour. The second enables detailed physico-chemical and mechanistic interpretation through systematic comparison with the WT protein. However, such analyses are only valid if the introduced mutations do not substantially distort the energy landscape or introduce alternative folding or assembly pathways. Unlike folded proteins, intrinsically disordered proteins are characterized by shallow, frustrated energy landscapes, making them particularly sensitive to even subtle sequence perturbations that can markedly alter their conformational or assembly behaviour. Consequently, it is essential to characterize the accessible structural states of each variant and interpret mechanistic differences with caution when mutagenesis substantially remodels the underlying energy landscape.

Here, we investigate to what degree systematic mutational analysis coupled with scalable bulk assays yield quantitative insights into how electrostatic interactions shape the αSyn energy landscape. By constraining the aggregation conditions to favour a limited set of microscopic pathways, we aimed to assess how these sequence changes reshape the energy landscape of αSyn under *in vitro* conditions reflecting specific biological contexts. We observed that mutational effects fall broadly into two categories: those that affect the energy barriers on a WT-like landscape, and those that re-shape the energy landscape and create minima not accessible by WT under the same solution conditions. Our work outlines a scalable, quantitative framework that increases the informational output of the mutational studies of αSyn using conventional assays. The presented approach can be extended by incorporating additional mutations and assays, with the aim of providing deeper insights into the energy landscape of αSyn and its link to physiological and pathological functions.

## Results

### Design, overview and naming convention of αSyn mutants used in this study

αSyn contains 15 lysines distributed across the sequence, grouped in clusters containing 1 to 3 lysine residues within imperfect repeats (Figure 2a). We substituted lysines with glutamines as an amino-acid with similar size to lysine with physiochemical properties mimicking lysine acetylation as a physiologically relevant posttranslational modification. (48,60–62) We generated 62 variants including all 15 single-point K-to-Q mutants, six mutant cluster variants (two 3-point mutants: KQ1 and KQ7; four 2-point mutants: KQ2, KQ3, KQ4, and KQ5), and their combinations (Figure 2a, Supplementary Table 1). We primarily focused on the KQ cluster variants (i.e., numbered according to their position in the sequence from N to C terminus, with KQ6 designating the single point mutation K80Q for completeness) as reporters of the sequence dependency of the lysine mutation. The rest of the variants were used to support and extend the observations made with the KQ cluster variants, e.g., discern contributions of individual mutations within each KQ cluster, or study epistatic effects between the clusters in aggregation assays (Figure 1b, Supplementary Figure 1).

**Figure 1.**
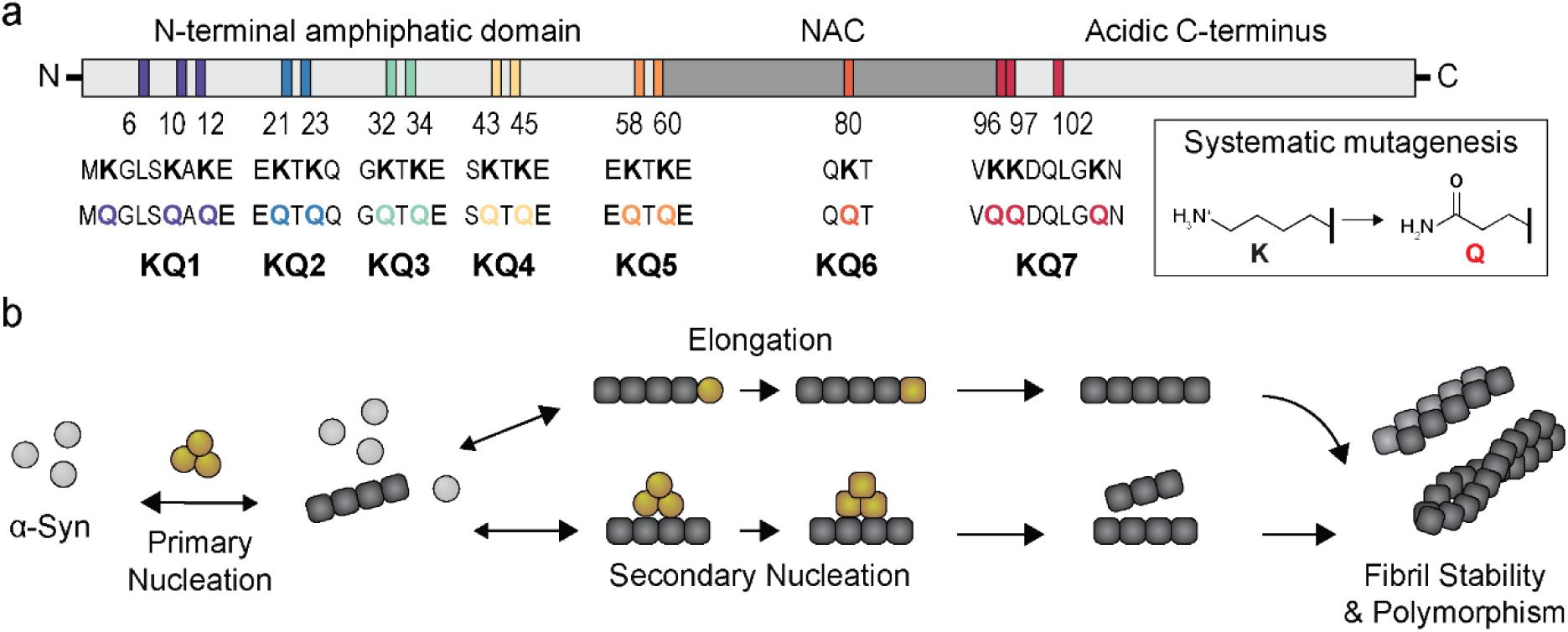
Overview of the αSyn mutants and aggregation pathways involved in this study. (a) Schematic overview of the αSyn sequence with highlighted positions of lysine residues mutated to glutamines in this study. The αSyn variants containing 2-3 mutations of adjacent lysines are referred to as KQ1-7 cluster mutants based on their position in the sequence from N to C terminus (e.g., KQ1 = K6Q + K10Q + K12Q, KQ2 = K21Q + K23Q, etc., Supplementary Table 1). **b) Schematic view of the distinct aggregation pathways and aggregate properties of αSyn** probed in this study.

**Figure 2:**
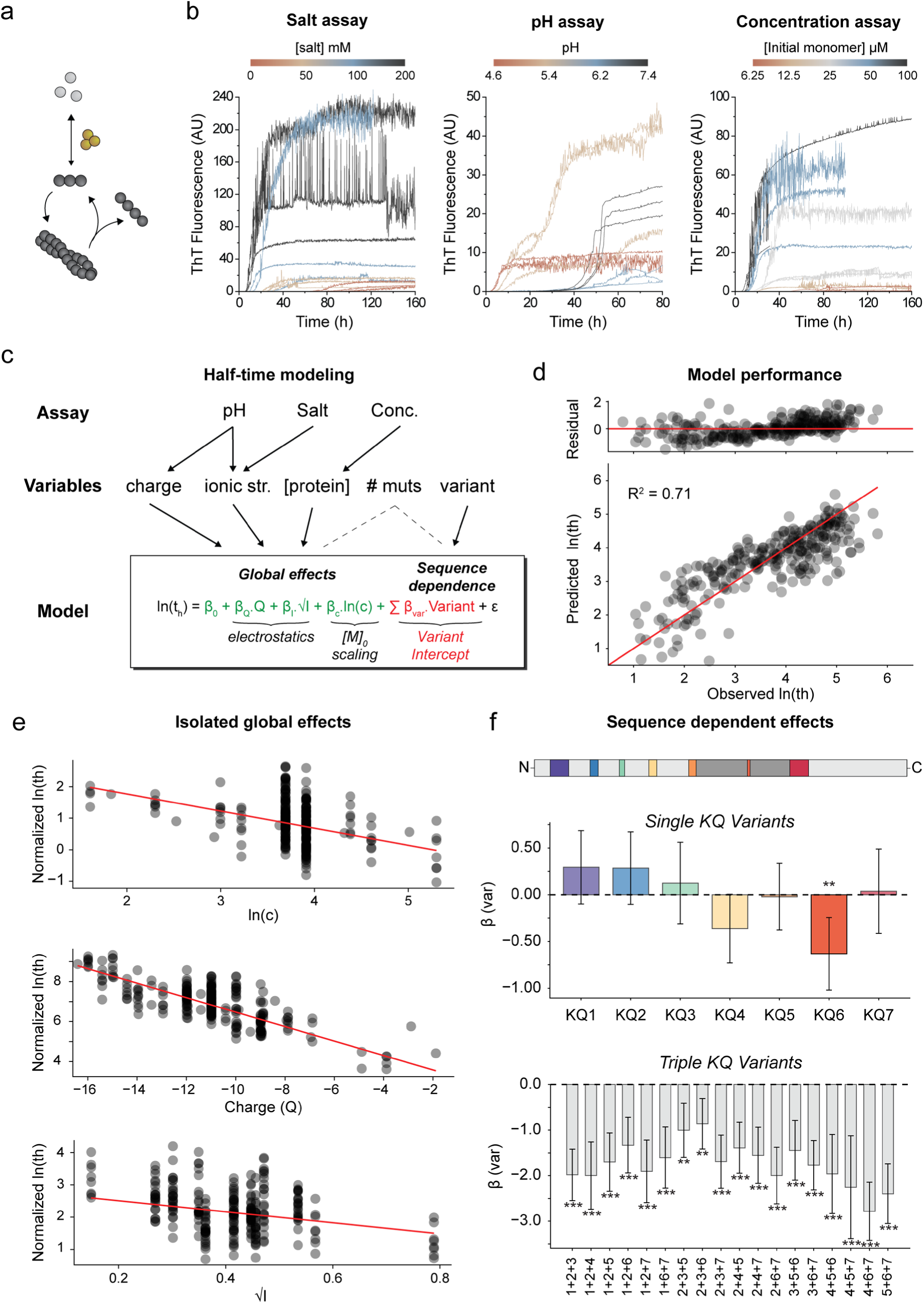
Effect of KQ mutations on *de novo* aggregation of α-Synuclein. (a) Dominant microscopic aggregation pathways probed in the unseeded ThT assays. **(b) Example of ThT aggregation kinetics of αSynWT** as a function of (left) increasing salt concentration, (middle) pH, and (right) initial monomer concentration. Raw data for all mutant variants are shown in **Supplementary** Figures 2-4. **(c) Weighted-least squares linear regression model of aggregation half-times.** The selected parameters that were varied across the assays were used to model log-transformed aggregation half-times ln(th) as a combination of global and variant specific effects. The global terms involved charge (Q), ionic strength (√I), and initial monomer concentration (ln(c)) multiplied by global slopes shared across the variants (β), and global intercept (β0) representing the global baseline. The sequence dependent term consisted of variant-specific coefficients (βvar) and binary variant encodings. ε denotes residual variance between fit and experimental data. **(d) Model evaluation.** Comparison of observed log-transformed half-times and their prediction by the model. The proportion of explained variance (goodness-of-fit, adjusted R^2^) is shown. The red line corresponds to 1:1 line, with residuals showing difference between observed and predicted values. **(e) Isolated global effects from the model.** The model prediction (red line) is overlayed with the observed data (black circles) normalized for all but one model parameter (e.g., for top graph depicting initial monomer scaling: norm_ln(th) = ln(th) − (βQ*Q + βI*√I + βvar*Variant)). **(f) Variant-specific intercepts** for single (top) and triple (bottom) cluster variants. Positive values indicate that the variants are on average slower compared to the global model predictions, whereas variants with negative intercepts are faster. Error bars indicate ± 95% confidence intervals. Asterisks denote levels of statistical significance: * p < 0.05; ** p < 0.01; *** p < 0.001.

### Dissecting global and sequence-specific effects of KQ mutations in α-Synuclein aggregation

To gain a global understanding of how positive charge removal influences αSyn aggregation (Figure 2a), we analyzed the aggregation kinetics across a range of solution conditions, including varying monomer concentration, pH, and salt using a Thioflavin-T (ThT) fluorescence assay (Supplementary Figures 2-4, Example data for WT Figure 2b). Each measurement was fitted with a logistic function (Equation M1, materials and methods) to obtain aggregation half-times, yielding a comprehensive dataset of 268 measurements (Supplementary file 1).(63) We parameterized all datapoints by variables describing both extrinsic factors (buffer type, salt type, ionic strength, pH) and intrinsic sequence features (number of mutations, variant, charge, Equation M2) and used them to model the aggregation half-time by linear regression (Figure 2c).

We tested models based on three key assumptions. (i) Under a simple nucleation– polymerization mechanism, the log-transformed aggregation half-time (ln(t_h_)) scale linearly with the logarithm of the initial monomer concentration (ln(c)) and with the square of the net charge, consistent with interactions between two charged monomers. (ii) The majority of the observed variance in aggregation kinetics can be accounted for by global physicochemical descriptors. (iii) Any residual variance can be explained by a combination of variant-specific effects on aggregation, other sources not accounted for by global descriptors, and experimental noise.

To this end, we modeled ln(t_h_) as a function of initial monomer concentration, nominal charge, ionic strength, and variant-specific intercepts (Equation 1, Figure 2c). The global terms accounting for electrostatics and protein concentration alone (Equation 2) explained ∼30% of the total variance, which increased to 56% when the number of mutations was included (Equation 3, Supplementary Figure 5a). Using variant-specific intercepts instead of number of mutations improved the fit (adjusted R² = 0.71, Equation 1, Figure 2d), consistent with assumptions (ii) and (iii): global scaling parameters capture most of the variance, but sequence-dependent effects still contribute significantly beyond noise. Statistical tests supported linear over quadratic charge scaling, but since the difference was only marginal, we refrain from drawing further mechanistic conclusions (Supplementary Tables 2 and 3). In the final model (Equation 1), we therefore implemented a linear scaling of charge (Q) and square root of ionic strength (√I) as parameters that modulate electrostatic interactions (Figure 2c).

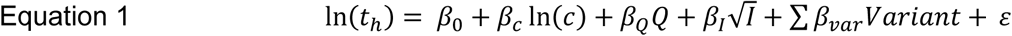

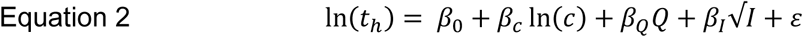

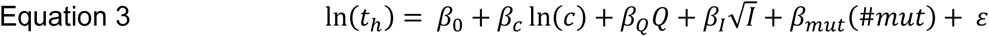

Where β_0_ is the model intercept (baseline), ln(c) is logarithm of initial monomer concentration, Q is theoretical net charge, I is ionic strength, #mut is the number of mutations, Variant is a binary encoding for each variant, β_c_, β_Q_, and β_I_ are coefficients for the global parameters, β_var_ are the variant-specific coefficients (Figure 2f), and ε is the residual error between fitted and experimental data representing unexplained variance (Supplementary Figure 5e).

The model comprised the global physical factors known to influence αSyn aggregation and described it well. Specifically, the model captured the increasing aggregation rate with decreasing net charge (β_Q_ = -0.4) and a strong aggregation-enhancing effect of ionic strength via charge screening (β_√I_ = –1.7) (Figure 2e, Supplementary Table 2). The global monomer scaling coefficient (β_c_ = - 0.5) is consistent with weakly monomer-dependent secondary processes (e.g., fragmentation, or saturated secondary nucleation) as dominant aggregation mechanisms under these conditions, as previously described for WT αSyn (43,64–68).

We modelled the sequence-dependent effects through variant-specific coefficients that shift the global baseline by fixed intercept (Figure 2f). This approach assumes that all variants share the same scaling with respect to global parameters as WT—a simplification that avoids overfitting but may not strictly hold. We observed that variants containing mutations in multiple (three) lysine clusters systematically accelerated aggregation beyond what could be explained by global effects or by the additive contributions of single-cluster mutations. Arguably, this effect can be caused by (i) a discrepancy between the theoretically calculated and the actual net charge (69), (ii) distinct aggregation mechanisms of triple KQ cluster variants, such as altered nucleation pathways or enhanced secondary processes; or (iii) non-linear contributions and epistatic effects of the individual cluster mutations. We cannot distinguish between these scenarios, since these variants have been tested in only one type of assay. We confirmed the robustness of the model coefficients by re-fitting it to a dataset excluding the triple-cluster variants to avoid potential bias (Supplementary Table 3).

The effects of the single-cluster mutations were generally mild, with KQ6 significantly accelerating aggregation (Figure 2f). Although not statistically significant, the coefficients of the remaining KQ clusters were highly consistent across different tested models, whether fitted to the full dataset or with the triple-cluster variants excluded (Supplementary Tables 2 and 3). On average, variants with mutations at the N-terminus (KQ1–KQ3) had inhibitory effects (slower aggregation), those with mutations of lysines found near or within the fibril cores (KQ4 and KQ6) had enhancing effects (faster aggregation), while the remaining variants (KQ5 and KQ7) exhibited behavior close to the wild-type (Figure 2f, Supplementary Table 2). Variants KQ4 and KQ6 displayed the largest residual errors (Supplementary Figures 5e), mostly in datasets where concentration and type of salts were varied (Supplementary Figures 2d, 4e). This indicates their altered sensitivity to solution conditions and potential deviation from the WT-like global aggregation behavior captured by the model.

### Monomer compaction is a poor predictor of half-time scaling

Although simple, the model captured both the global and sequence-dependent effects reasonably well given the noise level of the raw data (noise ceiling for triplicate means based on Fisher r ≈ 0.85; Supplementary Figure 2-4, Figure 2). Consistently, we find that the effects of the charge modulations persist even at 200 mM salt, where long-range electrostatic interactions are screened (Debye length < 1 nm), suggesting that modified local interactions are responsible for the altered aggregation propensity. To investigate the effect of mutations on the conformational space of αSyn monomers, we complemented our experimental analysis with coarse-grained molecular dynamics simulations of WT and KQ cluster variants using the CALVADOS2 force field (Supplementary Figure 6).(70) Across individual variants, we observed negative correlations between aggregation half-times and single-chain compactness, measured by the radius of gyration (R_g_) (Supplementary Figure 6d). However, neither R_g_ nor other ensemble-averaged descriptors captured the global variation in experimental aggregation kinetics of all variants (R^2^=0.27, Supplementary Figure 6d). In particular, the half-time scaling at physiological salt concentration remained unexplained, as all variants sampled predominantly extended conformations yet exhibited widely differing aggregation kinetics (half-times of ∼30–130 h, Supplementary Figure 6d). Together, these findings suggest that while single-chain compaction is correlated with aggregation kinetics, the sequence dependence of the KQ mutants arises from altered localized interactions rather than monomer conformational properties. This finding reflects the fact that the free energy landscape of an isolated monomer is distinct from that of a monomer in contact with another monomer or a fibril end, because of the additional electrostatic and other interactions provided by the intermolecular contacts.

### Variants with KQ mutations near the fibril core form stable fibrils that are not efficiently elongated by wild type αSyn

In order to gain deeper mechanistic understanding in the role of changes in global and local electrostatic interactions, we studied the effects of KQ mutations on αSyn fibril polymorphism, thermodynamic stability and seeding properties. We prepared fibrils from all KQ cluster variants under the conditions at which WT structures of polymorphs 2a and 2b have been solved previously using cryo-EM microscopy (Table 4). (71)

First, we used urea depolymerization and microfluidic transient incomplete separation to measure the thermodynamic stability of the fibrils (Figure 3a and b, Supplementary Figure 7) (72). We used Bayesian inference with Markov Chain Monte Carlo (MCMC) sampling to fit the data and to verify that no parameter correlation between Gibbs free energies (ΔG) and denaturant m-values obscured our conclusions (Supplementary Figure 7). (73,74) Fibrils formed by variants carrying mutations outside or at the periphery of the resolved fibril cores (KQ1, KQ2, KQ3 and KQ7) (75) exhibited stability comparable to or lower than that of the WT. In contrast, variants with mutations of residues resolved in αSyn fibril core structures (KQ4, KQ5, and KQ6) formed fibrils with higher stability than the WT under the tested solution conditions (Figure 3b, Supplementary Table 4).

**Figure 3.**
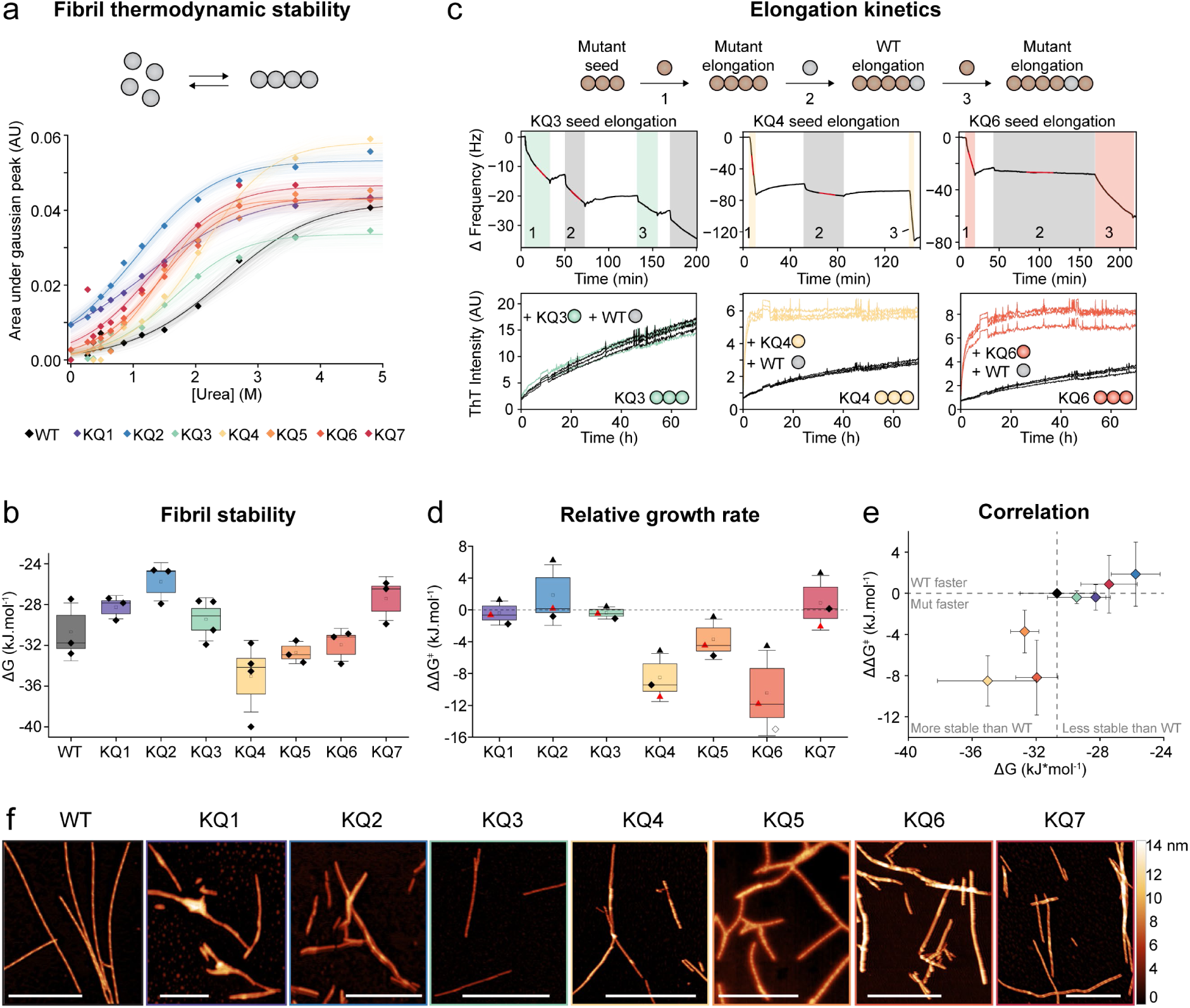
Analysis of fibrils formed by KQ cluster variants. a) Urea depolymerization of fibrils formed by KQ cluster variants. The monomer concentration in equilibrium with fibrils at increasing concentrations of urea was determined from the area under the Gaussian peak (y-axis) using transient incomplete separation in laminar flow. (72) The data were fitted to the isodesmic depolymerization model using Markov-chain sampling of solutions as described in our previous study. (74) Hundred randomly selected solutions from a total of 2000 sampled solutions are shown as curves with the best solution highlighted in bold. **b) Stability of WT and KQ fibrils** obtained from fitting the urea depolymerization experiments. The mean and median values from three independent measurements are depicted by open squares and line, respectively. The SE and SD (n = 3) are visualized as box and whiskers, respectively (Supplementary Table 4). The individual depolymerization curves and correlation plots between ΔG and the m-values are shown in Supplementary Figure 7. **c) Elongation kinetics** of KQ variant fibrils monitored by ThT fluorescence and quartz crystal microbalance (QCM)**. (Top and middle)** QCM experiments of KQ fibril growth. Immobilized variant fibrils were first allowed to elongate by variant monomers (colored boxes), followed by a washing step and elongation by WT monomers (grey boxes). The elongation rates of the variants and WT were quantified from the slopes (highlighted in red) of the changes of third harmonic overtone frequency (black lines) during the first and second injections, respectively. **(Bottom)** Comparison of aggregation kinetics of 80 μM WT (black) or KQ3 (green), KQ4 (yellow), or KQ6 (orange) monomers in the presence of 2.5 μM respective variant seeds monitored by ThT fluorescence assay. Aggregation of all variants together with analysis of monomer conversions to fibrils are shown in Supplementary Figure 8. **d) Relative growth rate (ΔΔG^ǂ^)** of mutant variants and WT monomers on fibrils of KQ cluster variants. Negative ΔΔG^ǂ^ values indicate that KQ seeds are elongated faster by their respective monomers compared to the WT monomer. The black and red symbols represent values derived from ThT and QCM experiments, respectively. The mean, median, SE, and SD of three independent measurements are depicted by open squares, line, box and whiskers, respectively. The open symbol in KQ6 corresponds to dataset where elongation of WT was not observed. **e) Correlation between stability (ΔG) and relative growth rate (ΔΔG^ǂ^)** of KQ fibrils. Datapoints and error bars correspond to the mean ± SE from b and d. **f) Representative AFM images of elongated fibrils.** The scale bar corresponds to 1 μm. Results from fibril height analysis together with all values related to this figure are provided in Supplementary Table 4.

We next quantified the changes to the free energy barriers of elongation (ΔΔG^ǂ^) from relative growth rates fibrils mutant and WT monomers on the fibrils formed by KQ cluster variants (Equation M3, Figure 3c and d) .(59) This strategy avoids complications that could stem from variations in the number of growing fibril ends between experiments (see method section for details). We observed little difference between elongation of KQ1, 2, 3, and 7 fibrils by WT and the respective mutant monomers. In contrast, variants KQ4, KQ5 and KQ6 which carry mutations near the known fibril cores showed distinct behaviour. WT monomer was inefficient in elongating their seeds, especially those formed by KQ4 and KQ6, even though elongation by their respective KQ monomers was fast.

To understand the structural bases for such differences, we characterized the morphology of the elongated fibrils in terms of their height and apparent helical pitch length using atomic force microscopy (AFM) (Figure 3f, Supplementary Figure 9). The WT fibrils were formed by two distinct populations characterized by shorter (ca. 160 nm, pink box in Figure 4f) and slightly longer (ca. 220 nm, green in Figure 4f) apparent pitch lengths with heights ranging from 6 to 8 nm. The KQ7 variant formed WT-like fibrils, whereas KQ5 fibrils had regular surface patterns with short helical pitch (∼100 nm). The rest of the variant fibrils appeared mostly flat or exhibited irregular patterns or short frequencies along their main axis (apparent pitch < 100 nm). The lack of twist in KQ4 and KQ6 fibrils was further confirmed by TEM, which revealed a dominant population of flat fibrils often forming stacks or clusters on the grid (Supplementary Figure 10). Fibrils formed by KQ1, KQ2, KQ3, and KQ5 variants exhibited lower heights (3–6 nm) compared to WT fibrils, suggesting either their tighter packing or that they are formed by a single protofilament (Supplementary Table 4). Interestingly, the denaturant m-values scaling was consistent with the expected dependence of exposed surface area in a rolled “Clarkson’s scroll” geometry (see Materials and Methods for details, Supplementary Figure 11). This suggests that the depolymerization m-values correlate with solvent exposure analogously to protein folding. (76)

**Figure 4:**
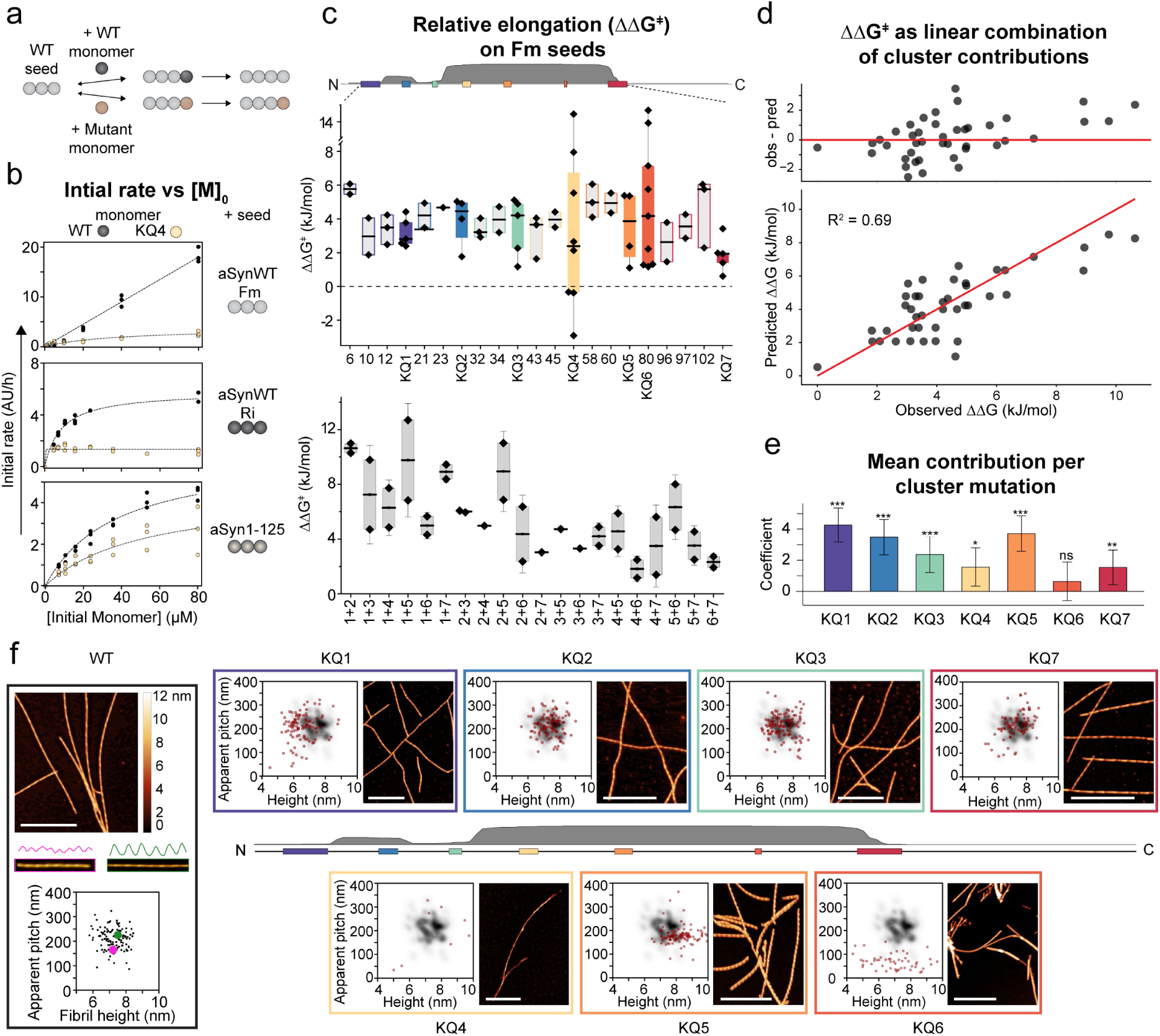
Effects of KQ mutations on αSyn WT fibril elongation. (a) Schematic overview of elongation as the dominant aggregation process. Either WT (grey) or mutant (gold) monomers elongate pre-formed WT fibrils with different efficiencies. **(b) Example initial rate analysis** from aggregation kinetics of WT (black) and KQ4 (yellow) monomers seeded by different WT (Fm - top, Ri - middle), or C-terminally truncated (aS125 - bottom) fibrils. The initial rates were obtained from linear fitting of the raw ThT data in the 0 – 2.5 h range. Fit to the equation describing saturating elongation (or linear function in case of the WT monomer on Fm) are shown as dotted lines. The raw data for all KQ variants elongation of Ri and aS125 are shown in Supplementary Figure 16. Fitting parameters from the initial rate analysis are provided in Supplementary Table 5. **(c) Mutational effects on the energy barrier of αSyn elongation. (top)** Schematic profile of αSyn sequence showing frequency of solved fibril structures where given residue was resolved (grey). Position of lysine clusters is highlighted by coloured boxes. **(middle)** ΔΔG^ǂ^ of single-point mutant and KQ cluster variants, **(bottom)** double cluster variants. Negative values (below dashed line) correspond to mutant monomer elongating WT fibrils faster than WT. The mean and median values are depicted by open squares and lines, respectively. The boxes represent 25 to 75 percentiles of the mean; outlier values are marked by whiskers. Examples of raw datasets and the ΔΔG^ǂ^ values are provided in Supplementary figures 12-16 and Supplementary Tables 5 and 6, respectively. **(d) Weighted least square regression of ΔΔG^ǂ^ values** using equation 5. Individual points correspond to the mean values from panel (c) with 1:1 line depicted in red. Residuals (observed-predicted) are shown on top. **(e) Variant-specific coefficients** βj from equation 5 representing the average contribution when single mutation, or whole cluster is mutated. Values correspond to the mean +/-standard errors. Asterisks denote levels of statistical significance: * p < 0.05; ** p < 0.01; *** p < 0.001. **(f) AFM analysis of products from elongation of WT-Fm seeds.** The apparent pitch lengths (i.e., 360 ° turn) and heights were extracted from the manually selected fibril profiles using an automated python script. (*59,72*) Kernel density estimation was applied to the morphological fingerprint (i.e., height vs pitch plot) of WT seeds elongated by WT monomer for better visualization. Each red dot corresponds to the profile of a single fibril. The height profile is color-coded according to the bar next to the WT image. White scale bars correspond to 1 μm. The sequence profile in the middle corresponds to the one in (c).

Altogether, we observed that the effects of the K-to-Q mutations on fibril stability are governed more strongly by their position within the sequence than their total number. AFM data of the fibrils formed by the different KQ cluster variants support the hypothesis that the fibril twists are modulated by interactions between N- and C-termini within the fibrils. (77) Our findings that near-core lysines (K43, K45, K58, K60, K80 and K96) modulate the fibril structure and stability is consistent with high-resolution structural characterization of αSyn WT fibrils, where these residues are found to stabilize protofilament interfaces by forming salt bridges or binding polyanionic molecules. (71,77–85). Hence, their perturbation by K-to-Q mutations could lead to formation of different fibril polymorphs. The correlation between variant fibril stability and their ability to be efficiently elongated by WT monomer (Figure 3e) suggests that mutations near the fibril core alter the conformational landscape of αSyn more dramatically, creating stabilizing interactions and structural features that are not easily amenable to WT elongation under identical solution conditions. Although the absence of a well-defined helical pitch precludes high-resolution structural analysis of these potentially distinct polymorphs, their incompatibility with WT monomer elongation likely arises from the energetic penalty associated with incorporating lysine residues into fibril cores where glutamines are present.

### Seeded growth of WT αSyn fibrils is governed strongly by sequence-specific mutational effects

We established that the effect of KQ mutations on the αSyn free energy landscape is sequence specific, yet it remained unclear whether the differential seeding competence is governed more by the intrinsic structural properties of the fibrils, or the monomers. We performed seeded aggregation assays to assess how the mutations influence fibril elongation across different structural polymorphs (Figure 4a and b). We used three types of seeds including full-length WT fibrils assembled under two distinct solution conditions that yield different sets of polymorphs (WT–Fm and WT–Ri) (71,74,86–88), and fibrils formed by a C-terminally truncated variant lacking six negative charges per monomer unit from the fuzzy coat (αSyn1-125, Table 4). We analyzed the elongation of WT–Fm seeds with 41 variants carrying one to six mutations to discern the contribution of the monomer (Figure 4c, Supplementary Figures 12-15) and further examined elongation of single KQ cluster variants (n=7) on the other two seed types to assess the influence of fibril structure (Figure 4b, Supplementary Figure 16). Relative elongation rates were then used to derive the changes of the energy barriers of elongation (ΔΔG^ǂ^), as described above (Figure 4c).

Elongation of the WT - Fm seeds by mutant monomers was slower compared to the WT (Figure 4c), except for a few repeats with KQ4. The average increase in energy barrier of elongation was 4.0 ± 1.2 kJ.mol^-1^, 3.8 ± 3.9 kJ.mol^-1^, and 5.4 ± 2.8 kJ.mol^-1^ for single-point mutations, KQ cluster variants (1-3 mutations), and double KQ cluster variants (3-6 mutations), respectively (Figure 4c, Supplementary Tables 5 and 6). Linear regression analysis showed that the monomer net charge (i.e., number of mutations) accounted for only ∼35% of the variance in ΔΔG^‡^ values, indicating a strong sequence-specific component to elongation (Equation 4, Supplementary figure 17).

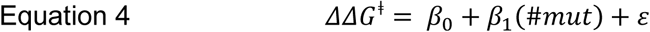

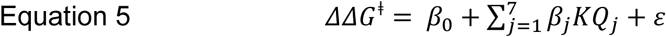

Where β_0_ is the global intercept (expected to be zero since ΔΔG^ǂ^ values are defined relative to WT), β_1_ is the global slope coefficient for the number of mutations (#mut), β_j_ are the variant-specific coefficients for mutations in cluster j (j=1-7), KQ_j_ is the binary encoding for mutations in cluster j, and ε is the residual error between fitted and experimental data representing unexplained variance (Supplementary figure 17).

Instead, we used cluster-specific coefficients representing the average contribution from each KQ cluster (i.e., either single mutation or whole cluster mutated) (Equation 5, adjusted R² = 0.69, Figure 4d). Among these, clusters KQ1, KQ2, KQ3, and KQ5 showed the most pronounced inhibitory effects on elongation of WT seeds, whereas the other clusters had weaker or statistically insignificant contributions (p > 0.05, Figure 4e). The model (Equation 5) captured well the main trends observed for the ΔΔG^‡^ values. First, the effects of single-point mutations within individual clusters were non-additive, as shown by comparison of their ΔΔG^ǂ^ values with those obtained when the entire KQ cluster was mutated (Figure 4c). Second, the combined effect of mutating two distinct clusters was well approximated by the mean of their individual effects, indicating near-additive behaviour between different clusters.

The strong effects of K to Q mutations outside of the fibril core (e.g., K6Q, K102Q, Figure 4c) underscore the important role of the electrostatic interactions of the fuzzy coat in the elongation kinetics of αSyn fibrils. (56,89,90) In comparison, the variants with mutations near core (KQ4 and KQ6), showed unusually large variation in ΔΔG^ǂ^ values. (Figure 4c). Further analysis using WT-Fm seeds of different maturation ages revealed that their elongation rates were highly dependent on seed age (Supplementary Figure 18). Given that polymorph composition evolves during fibril maturation (74), the pronounced variance in ΔΔG^‡^ values likely arises from fibril polymorph heterogeneity combined with variant-specific polymorph selectivity, consistent with the distinct aggregation behaviours of KQ4 and KQ6 observed in other assays. This indicates that K43+K45 and K80 are critical for recognizing specific αSyn fibril folds in line with other reports. (48,54)

These conclusions are consistent with the AFM analysis of the fibrils elongated by different monomers (Figure 4f). We observed WT-like fingerprints for monomer variants with mutations in the fuzzy coat (KQ1, KQ2, KQ3, and KQ7), whereas elongation by variants with mutations in the core resulted in only subpopulations of fibrils with short apparent pitch (KQ5) or mostly flat fibrils (KQ4, KQ6) (Figure 4f, Supplementary Figure 9).

Comparable results of the KQ cluster variants were obtained when elongation was studied using WT–Ri and αSyn1–125 fibrils as seeds. The ΔΔG^‡^ values did not show significantly different trend across all three polymorphs, but the different seed types varied in the saturation behaviour of the elongation rates as a function of monomer concentration, perhaps reflecting their distinct surface properties (Supplementary Figure 16, Supplementary Table 5). (91) N-terminal variants (KQ1–3) show slower kinetics and moderate-to-high saturation elongation constants (*K*e) across all three polymorphs, consistent with weakened initial monomer-fibril interactions. Variants with mutations near the core (KQ4–6) display lower *K*e values and polymorph-dependent kinetics, possibly suggesting rapid monomer attachment with rate-limiting rearrangement step or increased propensity of monomers to attach in an out-of-register conformation. (92) The KQ7 variant exhibits WT-like kinetics and saturation.

Together, our results show that templated aggregation is influenced by both the intrinsic properties of the fibril and the sequence features of the monomer, with the latter playing a dominant role in defining elongation kinetics. The sequence-dependent effects of K-to-Q mutations highlight critical roles of lysines in shaping the complex polymorphic landscape of αSyn. However, it remains uncertain whether the observed effects genuinely reflect the removal of interactions facilitated by the lysines in the wild-type protein, or if they instead result from newly enabled interactions specific to glutamine residues. (56)

### Secondary Nucleation and Polymorph Specificity of KQ Variants in Seed Amplification Assay at low pH

Finally, we investigated whether the same principles govern αSyn aggregation under conditions where secondary nucleation plays a dominant role (Figure 5a). We used the solution conditions established in our recently developed seed amplification assay (SAA) in which aggregation is dominated by secondary nucleation (i.e., low seed concentration, quiescent conditions, pH 3, 250 mM Na_2_SO_4_, Table 5) (93) and followed aggregation of WT or mutant monomers in the presence 1 nM WT-Fm or WT-Ri seeds (Figure 5b, Supplementary Figures 19 and 20). To dissect the effects of fibril structure from monomer properties, we modelled the log-transformed aggregation half-times of 26 αSyn single and double cluster KQ variants and WT (Figure 5c, Supplementary Table 7) by linear regression using Equation 6. Analogously to relative growth rates at neutral pH, the mutational effects on ln(t_h_) were sequence-dependent and could be modelled reasonably well by a linear combination of cluster-specific coefficients (Equation 6, adj R^2^=0.66, Figure 5d and e). Interestingly, the trend of variant specific contributions was similar compared to the relative growth rates at neutral pH; mutations in N-terminal (KQ1–KQ3) and KQ5 clusters prolonged the aggregation half-time, while KQ4 and KQ6 had effects similar to WT, and KQ7 exhibited moderately accelerated aggregation. This result suggests that secondary nucleation under low-pH and elongation at neutral-pH are driven by interactions between similar sequence regions.

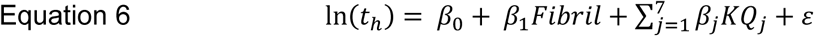

**Figure 5.**
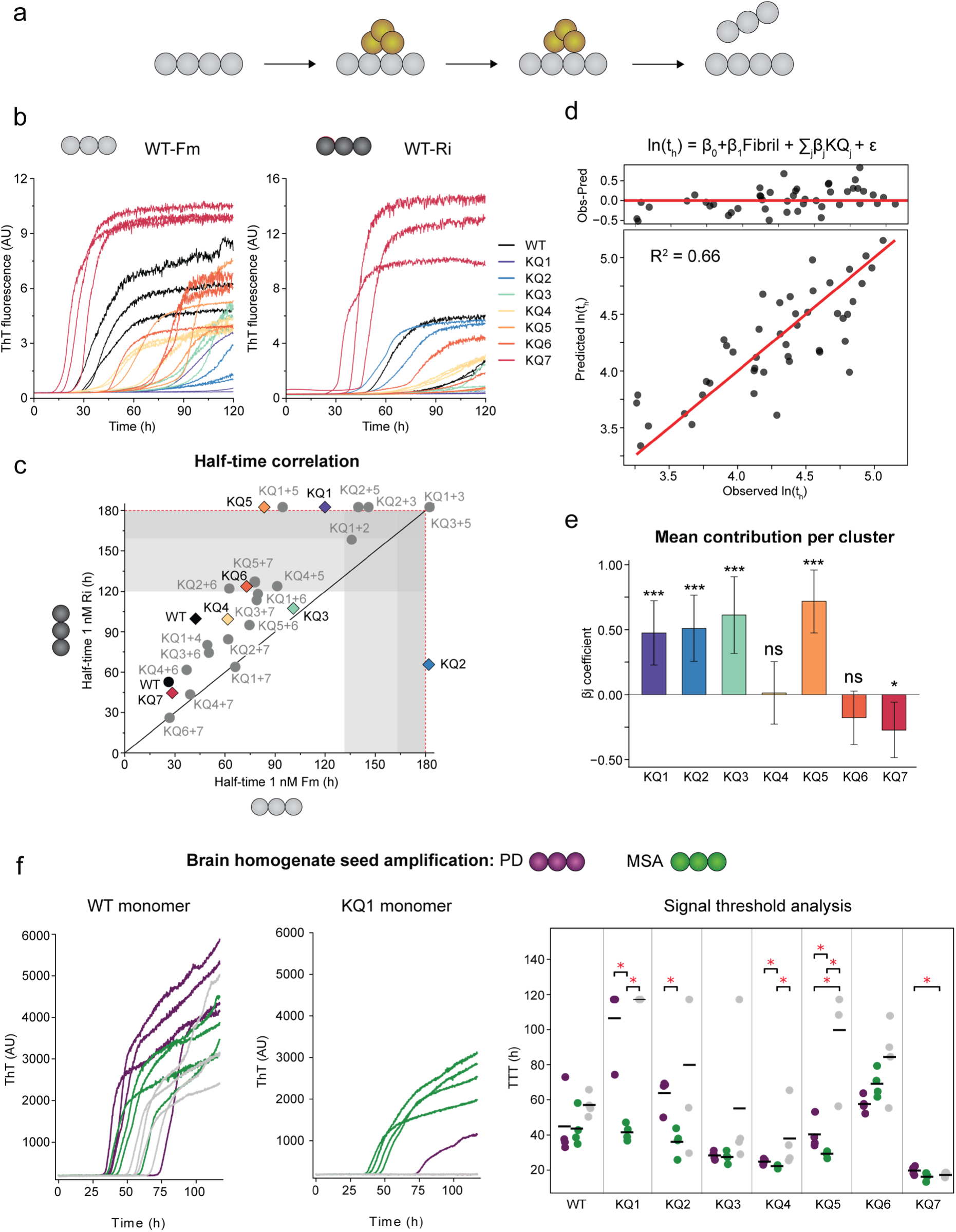
Effect of KQ mutations on seed amplification of two WT polymorphs at low pH. (a) Schematic overview of dominant mechanism probed in the assay. At low pH (pH = 3), amplification of pre-formed seeds is dominated by the secondary nucleation (yellow) of monomers on the fibril surface. **(b) ThT kinetics of WT and KQ cluster variants in the presence of 1 nM WT - Fm (left) or WT -Ri (middle) seeds.** The seeded aggregation kinetics of all variants carried out in the presence of 0, 1 nM, and 1 μM of sonicated WT seeds in conditions described in our previous study are show in Supplementary Figures 19 and 20. (*93*) (c) Correlation between aggregation half-times in the presence of 1 nM Fm and Ri seeds. The values correspond to the mean half-time values obtained from fitting the raw data from experiments carried out in triplicates in two independent measurements (circles, diamonds). The grey zones correspond to the endpoints of the measurements and points within these zones were derived with lower confidence. The points outside of the red lines correspond to the variants whose ThT signal did not plateau during the experiments and are shown for illustration. The black line has slope of 1 to easily visualize the specificity of the mutant monomers to different WT polymorphs. **d) Weighted least square regression of log transformed half-times** using equation 6. Individual points correspond to the mean values from panel (c) with 1:1 line depicted in red. Residuals (observed-predicted) are shown on top. **(e) Variant-specific coefficients** βj from equation 6 representing the average contribution when a whole cluster is mutated. Values correspond to the mean +/-standard errors. Asterisks denote levels of statistical significance: * p < 0.05; ** p < 0.01; *** p < 0.001. **(f) Seed amplification of brain homogenates from patients with Parkinson’s (PD, violet) and multiple system atrophy (MSA, green).** The amplification was carried out using 10 μM WT (left) or variant (e.g., KQ1, middle) monomer in the presence of PD (violet), MSA (green), or healthy (control, grey) brain homogenate diluted 10^5^ times into the assay buffer. The reactions carried out in quadruplicates were monitored by ThT fluorescence. The time to reach a threshold (TTT, right) was calculated as a 10 x SD of the mean signal between first and fifth hour of each dataset (Supplementary Table 8). For datasets where TTT could not be detected during the experiment, the end time of the experiment was used instead. Statistical significance between groups was assessed using two-sided Mann–Whitney U tests, with significance levels indicated by stars (* p < 0.05, ** p < 0.01, *** p < 0.001).

Where Fibril is the binary encoding for fibril type (i.e., WT-Fm or WT-Ri).

In contrast to the analysis of the relative growth rates at neutral pH, including the fibril type into the model significantly improved the quality of the fit (Equation 6). The positive coefficient for Ri fibrils (β_Ri_ = 0.4) compared to the Fm baseline reflects the overall slower kinetics observed for this polymorph. Although this difference may arise from distinct surface properties of the two fibril types, we cannot exclude the possibility of a minor systematic deviation due to concentration differences introduced during fibril preparation and handling (estimated at 4 – 10%).

Notably, amplification of the seeds by a handful of variants was specific to the polymorph type. Specifically, variants KQ1 and KQ5 amplified Fm, but not Ri fibrils, whereas the opposite trend was observed for the variant KQ2 (Figure 5c). Interestingly, KQ2 is a double-point mutant that includes K23Q, a mutation commonly used as a substrate in the state-of-the-art SAA protocols. (94,95). To investigate whether the polymorph specificity is more general phenomenon, we used KQ cluster variants to amplify seeds from brain homogenates of patients with PD and MSA (Figure 5f). The addition of minute amounts of brain-derived samples (10^5^-fold buffer dilution) decreased the lag time of aggregation compared to buffer-only control in most cases (Supplementary Figure 21). Importantly, variants KQ1, KQ2, and to lesser degree KQ4 and KQ5 showed significantly higher sensitivity towards PD samples compared to those from MSA patients (p<0.05, Figure 5f). This finding corroborates our results with *in vitro* generated fibrils and highlights the potential of using selected variants in seed amplification assays to distinguish between disease-specific polymorphs from patient samples.

## Discussion and conclusions

In this study, we carried out a comprehensive mutational analysis of α-synuclein (αSyn) to (i) quantify how electrostatic interactions contribute to the kinetics and thermodynamics of its assembly, and (ii) assess the broader applicability of this sequence-perturbative approach for probing the degenerate energy landscapes characteristic of self-associating intrinsically disordered proteins (IDPs).

Several systematic mutational studies aiming to elucidate sequence determinants of αSyn aggregation have been carried out (see, for example, (96) for their review). However, their global mechanistic interpretation is often difficult, due to the reported aggregation half-times or rates stemming from experiments that have been conducted under conditions influenced by many variables—such as shaking speed, the presence of beads, buffer composition, reaction vessel size, and, importantly, the existence of multiple aggregation pathways. These factors can significantly impact the observed aggregation behaviour, making it difficult to isolate sequence-specific effects and carry out quantitative comparison between results of different studies.

We circumvented the considerable variability inherent to αSyn aggregation data by studying the effects of mutations in (i) large number of experiments, (ii) under controlled conditions, where one or a few well-defined microscopic steps dominate the aggregation process, and (iii) using WT seeds where possible to constrain the structural changes to a minimum. We recently demonstrated that amyloid fibril elongation, which can be viewed as a templated-folding reaction, can be studied using conservative mutations and using WT seeds in all cases. (59) Under well-controlled conditions, this approach provides structural insights into the transition state ensemble similar to studies of protein folding.

Here, we investigated whether this modelling approach can be extended to different types of mutations and aggregation steps beyond elongation, within a more complex and polymorphic energy landscape. We find that, with few exceptions, K-to-Q mutations impair or slow down the aggregation of αSyn under all conditions studied. Aggregation from homogeneous monomer solutions can be modelled reasonably well (R^2^=0.53) by global scaling of intrinsic (net charge, protein concentration) and extrinsic (ionic strength) properties. In contrast, sequence-specific contributions become significant under conditions where aggregation is governed by monomer–fibril interactions, and the energy landscape is dictated by the available fibril structure. We observe similar magnitudes of energy changes (relative to the WT reference) for the fibril growth and fibril stability of the mutant variants (Figure 3), most of which displayed distinct morphological features compared to WT fibrils. Unlike Φ-value analysis, the structure—and thus the stability—of the KQ fibrils was not imposed by seeded growth using WT fibrils. Instead, we interpret the observed correlation such that the loss of electrostatic interactions important for WT seed growth promotes the formation of new interaction networks, stabilizing distinct fibril polymorphs in some of the KQ variants.

In another set of experiments, where we imposed the structure of WT fibrils by using them as seeds, we were able to robustly quantify the increase of the energy barrier of elongation to be around 4-6 kJ/mol which is in a range comparable to disruption of a surface-exposed salt bridge of folded proteins. (97) We find that changes on the order of 8-10 kJ/mol (corresponding to approx. two-thirds of the overall height of the energy barrier (98)) are sufficient to render variants essentially incompatible for elongation of WT seeds and vice versa (Figure 3Figure 4). The changes to the energy barrier are higher compared to stability perturbations predicted for single point mutations by FoldX on 47 representative WT polymorph structures (mean ΔG_0_= 1.6 ± 0.9 kJ/mol). (Supplementary Figure 22, Supplementary Table 9) Both experimental and *in silico* data exhibit substantial variability, consistent with the pronounced polymorphism of αSyn fibrils, which hinders determining whether contact changes arise during formation of the transition state or exclusively within the fibrillar state.

To gain further insights into how individual KQ cluster mutations shape the global αSyn free energy landscape, we grouped them based on their relative effects (compared to WT) observed in all our assays (Figure 6a). The results reveal that mutations found in the fibril cores (Figure 6b), K43Q+K45Q (KQ4) and K80Q (KQ6), have the largest and most distinct effects across all assays, specifically in seeding, cross-seeding, and fibril stability. These mutations are examples of perturbations that significantly modify the free energy landscape of the WT sequence, creating pathways and energy minima not accessible to the WT under the same experimental conditions (Figure 6c). These observations are consistent with other studies showing that acetylation of lysines K43 and K80, or modifications of adjacent regions, significantly alter aggregation and seeding, identifying them as key modulators of αSyn aggregation. (48,54,58,99). In contrast, mutations of lysines found outside of the fibril cores (KQ1-3, KQ7, Figure 6b) negatively affected energy barriers whilst overall maintaining a WT-like free energy landscape minima (Figure 6c).

**Figure 6.**
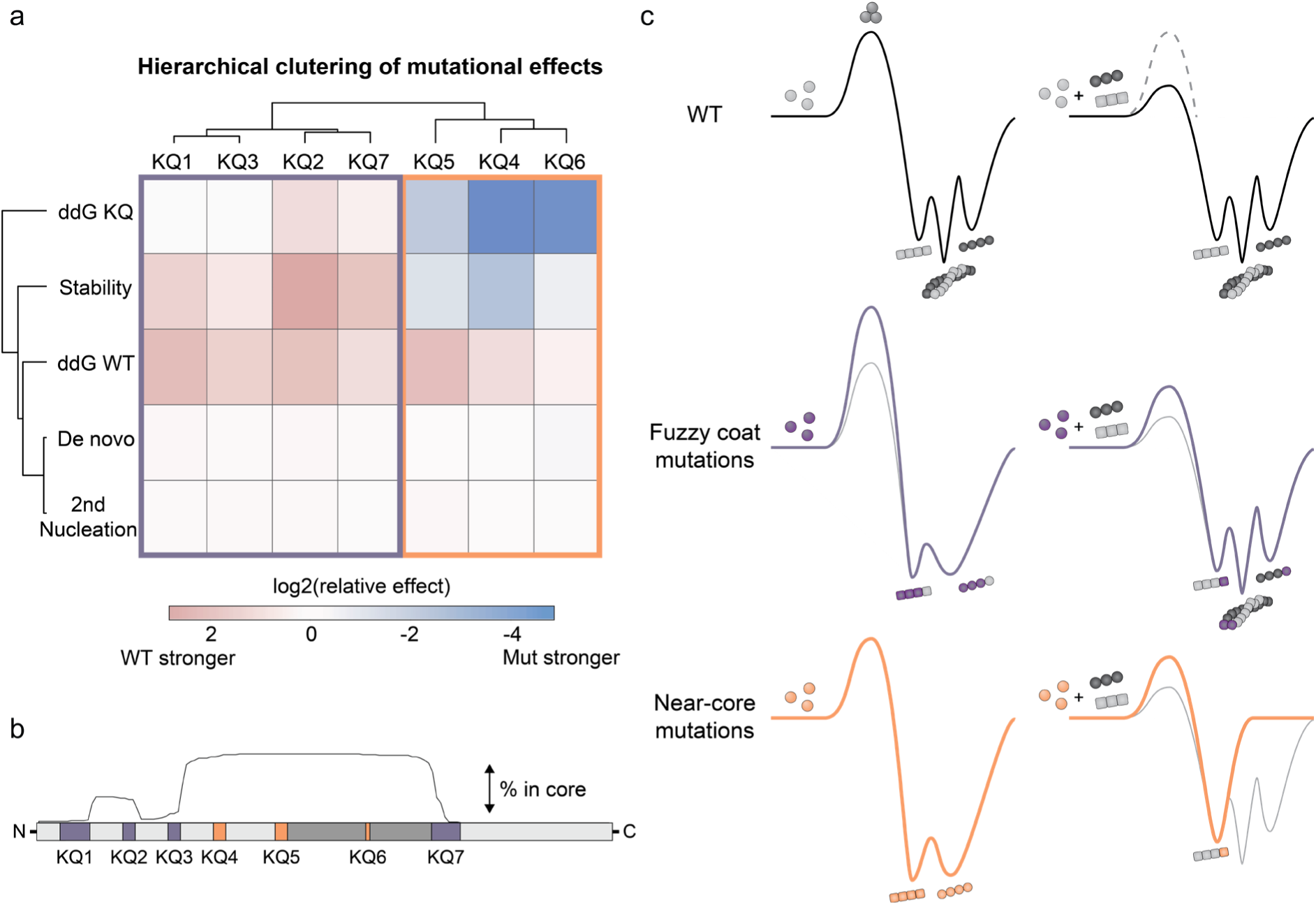
(a) Hierarchical clustering of mutations based on the magnitudes of mutational effects. Relative effects of mutations relative to WT across different assays were log2 transformed to have comparable amplitudes (see materials and methods for details). The assay data include differences of energy barriers between WT and mutant on respective KQ seeds (ddG KQ, Figure 3d), fibril stability (Figure 3b), cluster specific coefficients from wls regression of elongation on WT-Fm (ddG WT, Figure 4e), cluster specific coefficients from wls regression of half-times from unseeded experiments (De novo, Figure 2f), and cluster specific coefficients from wls regression of half-times from assays at low pH and low seed concentrations (2^nd^ Nucleation, Figure 5e). **(b) Sequence and structural context of the mutational effects.** The mutated lysine clusters are depicted in simplified color-coding scheme according to their clustering in (a). The graph above depicts frequency of the lysine residues resolved in available αSyn fibril structures. **(c) Schematic illustration of mutational effects on αSyn energy landscape. (Top)** WT energy landscape showing high barrier of de novo aggregation leading to multiple co-existing fibril polymorphs. Addition of WT seeds (right) leads to accelerated aggregation. **(Middle)** WT-like energy landscape of variants with KQ mutations of residues found in fuzzy coats of αSyn fibrils. These mutations slow-down aggregation (higher barrier) leading to polymorphs that are efficiently elongated by WT monomers. Conversely, mutant monomers can elongate WT fibrils seeds with generally slower kinetics. **(Bottom)** Altered energy landscape with KQ mutations of residues found near cores of αSyn fibrils. These mutations aggregate with kinetics similar to WT leading to polymorphs that are elongated by WT monomers with low efficiency. Conversely, mutant monomers are inefficient in elongating WT seeds, indicating selectivity to specific polymorphs.

The work presented here is an attempt to unify commonly used aggregation assays to assess the impact of sequence perturbations in a formalized and systematic manner. It highlights mutational studies as a tool to probe the αSyn energy landscape. Our results demonstrate that it can yield quantitative information, provided it is carried out under well controlled conditions and complemented by structural, or morphological analysis to ensure meaningful and interpretable results. The clustering in Figure 6 provides a compelling framework for understanding how specific mutations, in this case those that alter electrostatic interactions, modulate the aggregation landscape. However, expanding the mutational space will be essential for further validation and generalizing the observations made here. In this study, we selected mutations with the same chemistry to demonstrate the feasibility and scalability of such an approach. The scaled-down purification protocol developed here allows to obtain ca 30 αSyn variants within 10 days in sufficient amount (1-3 milligrams) and purity (>95%) to perform all assays presented here. Extending the dataset by including (i) different mutations of the same residues (e.g., alanine, glutamate), (ii) mutations of negatively charged residues within the imperfect repeats (e.g. E-to-Q mutations), (iii) and mutations targeting different physio-chemical properties (e.g., aromaticity, hydrophobicity) within specific sequence regions will provide more complex and comprehensive insights. Moreover, the framework could be extended by other assays e.g. formation of oligomers, and, importantly, cellular assays that would help bridge the observation of mutational effects *in vitro* to more physiologically relevant environments. Together, these efforts will help clarify the pathways and sequence regions responsible for the transition from physiological to pathological αSyn conformations. They will also help identify *in vitro* assays with readouts that are directly translatable to biologically relevant outcomes, that can be used for efficient and high-throughput drug screening. Finally, we demonstrate that certain mutant variants exhibit selectivity in seed amplification assays, enabling discrimination between disease-associated protein conformations, highlighting their potential for diagnostic applications.

## Materials and methods

### Mutagenesis

Lysine-to-glutamine (KQ) variants were prepared starting from pET29a_αSyn WT plasmid using Golden gate mutagenesis protocol. The primers were ordered from TAG Copenhagen A/S (Denmark), all chemicals, restriction enzymes and buffers from New England Biolabs (USA) unless stated otherwise. Protocol utilizes set of primers provided in Table 1 and consists of (i) amplification (Table 2 andTable ***3***), (ii) purification, (iii) restriction/ligation, and (iv) transformation. Linear products of amplification were purified from agarose gel after electrophoresis in 1 % TAE agarose (120 V, 30 min) using GFX PCR DNA and Gel Band Purification Kit (Cytiva, USA). The Golden gate assembly mixture was prepared according to Table 2 and incubated at 37 °C for 16 h. The reaction was stopped by 10 min incubation at 85 °C. Residual template DNA was digested DpnI (1U, 1.5 h, 37 °C incubation) that was then deactivated by heating (85 °C, 10 min). The individual reactions were pooled together, cleaned using the GFX PCR DNA and Gel Band Purification Kit (Cytiva, USA), transformed into the BL21(DE3) competent *E. coli* which were plated on LB-agar containing kanamycin as selection marker (50 μg/ml), and incubated for 12 h at 37 °C. Single colonies were transferred into the 96-well plate containing 50 μl of TE buffer using sterile toothpick and send for sequencing (Eurofins Genomics, Germany). The same colonies were simultaneously transferred to a 96-DW plate containing 1 mL of LB (kanamycin) which was then used to create respective glycerol stocks. KQ cluster variants created in the first round of mutagenesis (together with single-point mutants) were used as templates for preparation of the double-cluster variants, which were subsequently used as templates for the triple-cluster variants.

**Table 1.**
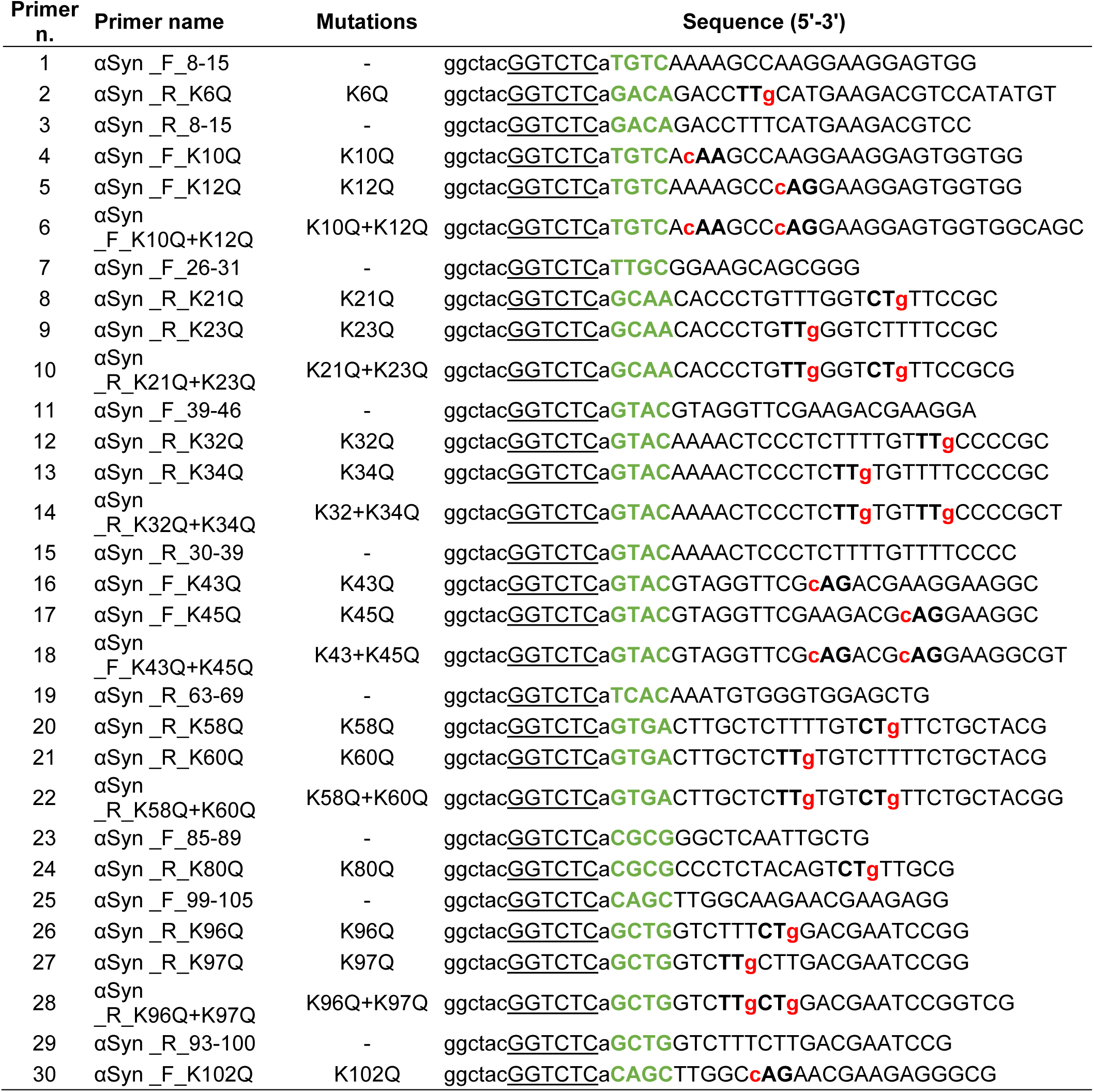
List of primers used for mutagenesis. Regions of complementarity between the pair of primers are shown in green and are followed by the sequences complementary to the plasmid. The mutated codons and nucleotides are highlighted in bold and red, respectively. Sequence upstream of the green contains random flanking sequence (lower case), BsaI recognition site (GGTCTC), and an extra base next to which BsaI cleaves. Reverse (R) and forward (F) direction of primer is encoded in the primer name.

**Table 2.** Composition of the reaction mixtures for mutagenesis (PCR 1) and Golden Gate Assembly (PCR 2). Phusion MM – master mix containing HF-Phusion DNA Polymerase, dNTPs, and buffer components.

| PCR 1 – amplification mixture |  |  | PCR 2 –Golden Gate Assembly |  |  |
| --- | --- | --- | --- | --- | --- |
| Component | Stock | Final | Component | Stock | Final |
| 2x Phusion MM | 2x | 1x | T4 buffer | 10x | 1x |
| Fwd_primer | 10 μM | 0.5 μM | T4 ligase | 400 kU/ml | 400 U/mL |
| Rev_primer | 10 μM | 0.5 μM | Bsal-HF | 20 kU/mL | 1.2 kU/mL |
| Plasmid DNA | 50-200 ng/μL | 100 ng | PCR mix | 50-200 ng/μL | 100 ng |
| mqH<sub>2</sub>O | - | - | mqH<sub>2</sub>O | - | - |

**Table 3.** Thermocycler protocol used for the PCR 1 mutagenesis. Times and temperatures were adjusted based on the size of the plasmid (approx. 5.8 kb) and fidelity of the HF-Phusion polymerase.

| Step | Temperature (°C) | Time (s) | n. cycles |
| --- | --- | --- | --- |
| Initial denaturation | 98 | 30 | 1 |
| Denaturation | 98 | 5 |  |
| Annealing | 55 | 15 | 28 |
| Extension | 72 | 180 |  |
| Final Extension | 72 | 600 | 1 |

**Table 4.** Conditions used for assembly of different fibril polymorphs used in this work.

| Fibrils | Buffer | T (°C) | Shaking (rpm) | Time (days) |
| --- | --- | --- | --- | --- |
| WT - Fm | 50 mM Tris-HCl 150 mM KCl pH 7.4 | 37 | 600 | 7-28 |
| WT - Ri | 5 mM Tris-HCl pH 7.4 | 37 | 600 | 14 |
| αSynS1-125 | 50 mM Tris-HCl 150 mM KCl pH 7.4 | 37 | 600 | 14 |
| KQ | 50 mM Tris-HCl 150 mM KCl pH 7.4 | 37 | 600 | 14-28 |

**Table 5.**
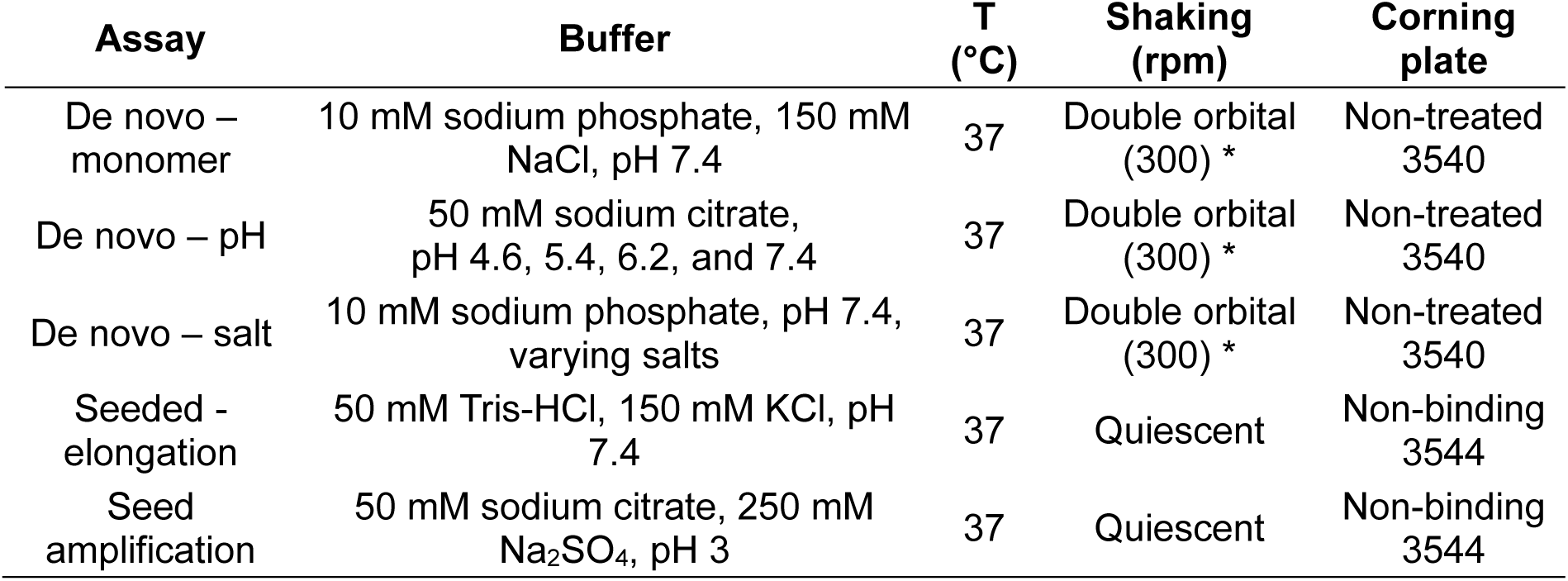
Overview of the conditions for different ThT assays used in this study. * Each well contained a single glass bead (d = 1 mm).

| Assay | Buffer | T (°C) | Shaking (rpm) | Corning plate |
| --- | --- | --- | --- | --- |
| De novo – monomer | 10 mM sodium phosphate, 150 mM NaCl, pH 7.4 | 37 | Double orbital (300) * | Non-treated 3540 |
| De novo – pH | 50 mM sodium citrate, pH 4.6, 5.4, 6.2, and 7.4 | 37 | Double orbital (300) * | Non-treated 3540 |
| De novo – salt | 10 mM sodium phosphate, pH 7.4, varying salts | 37 | Double orbital (300) * | Non-treated 3540 |
| Seeded - elongation | 50 mM Tris-HCl, 150 mM KCl, pH 7.4 | 37 | Quiescent | Non-binding 3544 |
| Seed amplification | 50 mM sodium citrate, 250 mM Na<sub>2</sub>SO<sub>4</sub>, pH 3 | 37 | Quiescent | Non-binding 3544 |

**Table 6.** Overview of the patients derived samples used for seed amplification assay.

| Group | Number | Age | Sex | Disease duration | Subtype |
| --- | --- | --- | --- | --- | --- |
| MSA | 1 | 69 | Female | 8 | MSA-P |
| PD | 1 | 64 | Male | 13 | - |
| Control | 1 | 77 | Male | - | - |

### Protein expression

#### Large scale protein expression and purification

Wild type, KQ cluster variants, and truncated variant (αSynS1-125) of αSyn were expressed in E. coli BL21 (DE3) cells transformed by the pET29a plasmid carrying the respective gene. The transformed cells were used to inoculate 1 L of LB medium containing kanamycin (30 μg.mL^-1^) as selection marker. Following the 3-hour incubation at 37 °C (OD_600_∼ 0.6–0.8), protein expression was induced by addition of IPTG (1 mM final concentration) and carried out for 4 hours at 37 °C. The cells were harvested by centrifugation (5,000×g, 20 minutes) and the resulting pellet resuspended in 20 mL of Tris buffer (10 mM Tris–HCl, 1 mM EDTA, pH 8.0) with 1 mM PMSF (phenylmethylsulfonyl fluoride). Cells were sonicated with a probe ultrasonicator for 8 min (10 s on time, 30 s off time, 12 rounds with 40% amplitude). 1 μL of commercial DNAse (Benzonase®) was added to the cell lysate and the insoluble fraction was removed by centrifugation (20 000×g, 30 min at 4 °C). Cell-free extract was boiled for 20 min and the heat-precipitated proteins removed by centrifugation (20 000×g for 20 min at 4 °C). αSyn was precipitated by addition of saturated (NH_4_)_2_SO_4_ (4 mL per 1 mL of supernatant). The solution was incubated on a rocking platform at 4 °C for 15 min and then centrifuged (20 000×g, 20 min, 4 °C) to obtain a protein pellet. The pellet was dissolved in 7 mL of 25 mM Tris–HCl pH 7.7 with 1 mM DTT. Protein was dialyzed against the same buffer for 16–18 h with a buffer exchange after 12 h of dialysis at 4 °C. The dialyzed protein was then subjected to anion exchange chromatography (AEC) (HiTrap Q Hp 5 ml, GE healthcare) followed by size exclusion chromatography (SEC) (HiLoad 16/600 Superdex 200 pg. column). The monomeric fraction of αSyn eluted in 10 mM of sodium phosphate buffer (pH 7.4) was collected, and the protein concentration determined by UV-absorption at 280 nm with theoretical molar extinction coefficients calculated from the protein sequence using ProtParam80 (Expasy, Switzerland).

#### Small-scale protein expression and purification

The small-scale expression of αSyn KQ variants was carried out in 90 mL LB medium analogously to the large-scale expression. Pellets from the harvested cells were resuspended in 100 mM MES, 750 mM NaCl, 1 mM EDTA, 1 mM PMSF pH 7 and heated to 80 °C for 30 minutes. Next, acetic acid (c_final_ = 1% v/v) and streptomycin (c_final_ = 1% w/v) were added, and insoluble fraction removed by centrifugation (20 000×g, 20 min, 4 °C). Resulting cell-free extract was precipitated by addition of saturated (NH_4_)_2_SO_4_ (4 mL per 1 mL of supernatant). The solution was incubated on a rocking platform at 4 °C overnight and then centrifuged (20 000×g, 30 min, 4 °C). Resulting protein pellet was dissolved in 4 mL of 10 mM NaP buffer pH 7.4 and dialyzed twice to the same buffer. Proteins were purified by AEC using AcroPrep 96-well filter plates (Cytiva, USA) and Multi-well Plate Vacuum Manifold (Cytiva, USA). Each well was loaded with 350 μL of Nuvia HP-Q strong anion exchange resin (Bio-Rad, USA). Resins were washed by 10 mM NaP buffer pH 7.4 before samples were applied (1 mL /well, 4 wells/sample). Unbound proteins were washed with 5 x 0.6 mL of 10 mM NaP buffer and 5 x 0.6 mL of 10 mM NaP buffer with 100 mM NaCl. Single-point mutants, KQ clusters variants, and double KQ cluster variants were eluted by10 mM NaP buffer supplemented with 250, 300, or 350 mM NaCl, respectively. Proteins were concentrated and buffer-exchanged using Amicon ® Ultra Centrifugal Filters with 3 kDa Mw cut-off (Merck, USA), aliquoted, flash-frozen in liquid nitrogen, and stored at -80 °C. Their concentration, purity and size distribution were determined using UV-absorbance (ε_280_ = 5,960 cm^-1^M^-1^), SDS-PAGE analysis and flow-induced dispersion (FIDA) analysis.

### Fibril preparation

A 100 or 200 μM αSyn WT, KQ cluster variants, or αSyn-C1-125 monomers were buffer exchanged into the corresponding assembly conditions (Table 4, (71,88)). After the incubation, fibrils were pelleted by centrifugation (16,000 x g, 60 min, 25 °C) and supernatant carefully removed. The residual monomer concentration was quantified from the isolated soluble fraction using UV-absorbance, SDS-PAGE and FIDA. The fibrils were resuspended in the respective buffer to final concentration of 100 or 200 μM (in monomer equivalents), flash-frozen in liquid nitrogen and stored at -20 °C. Prior to the experiments (seeding assays, chemical depolymerization), fibrils were thawed and sonicated using an ultrasonic probe (Hielscher UP200St). Sonication was carried out in repeating 1s-pulses of 100% amplitude separated by 1 second pauses for 4 minutes (two minute total sonication time).

### Thioflavin T assays

All ThT kinetic measurements were carried out using the FLUOstar Omega plate reader (BMG, Germany) using 440/488 excitation and emission and bottom reading every 5 or 10 minutes. All reactions contained 50 μM ThT (final concentration) and were carried out in triplicates with reaction volumes of 15 μL per well unless stated otherwise. The conditions of specific ThT assays are provided in Table 5.

#### De novo aggregation assay

Protein samples were buffer exchanged into the assay conditions (Table 5), diluted to final concentrations of 5 – 200, 40, and 50 μM for monomer dependence, salt, and pH screening, respectively, and supplemented by 50 μM ThT. Resulting kinetic curves were fitted to the sigmoidal function described by Equation M1:

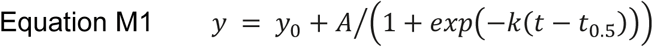

The y_0_ is the pre-transition baseline, A is the signal amplitude, k is the apparent growth rate, and t_0.5_ is the midpoint of the transition, i.e., half-time. (63)

The theoretical net charges of different variants at varying pH conditions were estimated using the Henderson-Hasselbach equation based on the protein sequence as

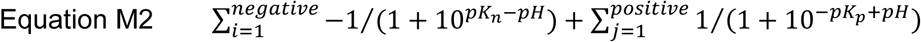

Where the pK_n_ and pK_p_ are dissociation constants of negatively and positively charged amino acid groups, respectively.

#### Seeded aggregation assay

A dilution series of the αSyn variants was prepared in 5-80 μM monomer range and sonicated seeds were added (final concentration 2.5 μM) just prior the measurement. A control reaction without ThT containing 40 or 80 μM of monomer was included for each series and used for quantification of the residual monomer concentration at the end of the reaction. A linear curve was fitted to the first 2.5 hours of the data and resulting slopes plotted against the initial monomer concentration ([M]_0_). The apparent elongation rate constants were extracted as the slopes in the linear data range of the plots ([M]_0_ ∼ 0-40 μM). To cancel out the contribution of the number of seeding-competent fibril ends ([S]), the effect of mutations was related to the elongation of WT measured on the same plate with the same batch of seeds according to Equation M3.

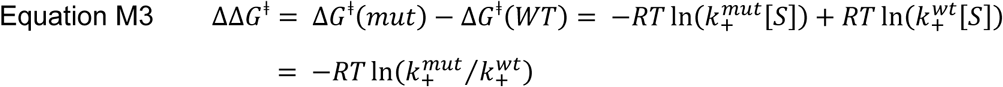

Where ΔG^ǂ^ is the Gibbs activation energy of elongation, R the universal gas constant, T the thermodynamic temperature, and *k*_+_ the microscopic elongation rate constant of wild type (wt) or mutant (mut).

In cases where saturation elongation was observed, initial rates were fitted to Equation M4.

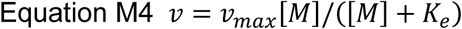

Where v and v_max_ is the observed and maximum rate, respectively, *K*_e_ is the elongation saturation constant, and [M] is the monomer concentration. (91)

#### Seed amplification assay at low pH

WT or mutant monomers (10 μM) were brought to the assay conditions by mixing from high protein concentration stock. Sonicated Fm or Ri fibrils were added to the samples to final concentrations of 0, 0.001, or 1 μM. The kinetic curves in the presence of 1 nM were fitted by Equation M1 to extract the aggregation half-times used for further analyses.

Brain samples were acquired from the Bispebjerg Brain Bank at Bispebjerg-Frederiksberg Hospital (University Hospital of Copenhagen, Denmark; Ethical approval: j.no.: H-15016232, data protection agency: j.no.: P-2020-937, Table 6). Brain tissue homogenates were prepared as follows: Approximately 50 mg of tissue samples were homogenized using a bead homogenizer (Precellys, Bertin Technologies) with 2 cycles of 45 seconds at 4,600 rpm in buffer containing 1x dPBS (Gibco), 1x HALTTM protease and phosphatase inhibitor cocktail (cat no.: 78444, Thermo ScientificTM) to a final concentration of 10% w/v. Aliquoted homogenates were stored at -80℃ for further use. For the plate assay, the brain homogenates were diluted 1000-fold further to the condition of seed amplification assay described in Table 5 with 10 μM monomer of different α-Syn variants.

### Weighted least-square regression of the data from aggregation assays

The data from unseeded experiments, elongation experiments with WT seeds at neutral pH, and mildly seeded experiments at low pH were compiled into three datasets. Variants were one-hot encoded at the single-mutation (elongation) or KQ-cluster (unseeded and low-pH) level, and each data point was parameterized by the number of mutations, pH, ionic strength, and net charge (Equation M2) derived from the experimental conditions. Experimental replicates were grouped by variant and fibril type to calculate mean ΔΔG values and corresponding standard errors of the mean (SEM), with each group assigned an inverse-variance weight (1/SEM²). For the unseeded and low-pH datasets, mean and standard deviation (SD) of log-transformed half-times from each triplicate measurement were used as a single data point and weights (1/SD), respectively. The noise level was estimated from the average correlation between replicate measurements, computed using Fisher’s z-transformed mean. Weighted least squares (WLS) regression and statistical analyses were performed in Python (v3.11) using statsmodels (v0.14).

### Quantification of residual monomer

#### UV absorbance

Samples without ThT were withdrawn from the plate and centrifuged to pellet down the fibrils (16,000 x g, 60 min, 25 °C). The concentration of monomer was determined by UV absorbance using NanoDrop (ThermoFisher, USA) and the extinction coefficient of αSyn (ε_280_ = 5,960 cm^-1^M^-1^) calculated from the sequence using Expasy webserver.

#### Flow-induced dispersion analysis (FIDA)

The oligomeric state analysis of the supernatant and monomer quantification were further determined using the FIDA1 instrument (FidaBio, Denmark). Samples were analyzed using the method provided in Table 7.

**Table 7.** FIDA method used for analysis of soluble αSyn fraction.

| Step | Component | Time (s) | Pressure (mbar) | Temperature (°C) |
| --- | --- | --- | --- | --- |
| Wash 1 | 1 M NaOH | 45 | 3500 | 25 |
| Wash 2 | Water | 45 | 3500 | 25 |
| Equilibration | Buffer | 40 | 3500 | 25 |
| Sample | Protein | 20 | 75 | 25 |
| Measurement | Buffer | 75 | 1500 | 25 |

The monomer concentration was quantified from the areas under the peak (obtained by fitting a Gaussian function to the Taylorgrams by the in-built software) using calibration curve of known monomer concentrations.

#### SDS-PAGE analysis

The samples were collected from the assay plate and centrifuged to pellet down the fibrils (16,000 x g, 60 min, 25 °C).The supernatant was mixed with the NuPAGE™ LDS Sample Buffer (ThermoFisher) in 1 to 1 ratio and applied to NuPAGE™ Bis-Tris Mini Protein Gels, 4– 12% (ThermoFisher). Calibration samples of SEC-isolated αSyn monomer of known concentrations were prepared in the same way. The electrophoresis was carried out at constant 200 V for 35 minutes, followed by staining using InstantStain Coomassie Stain (Kem-en-tec-nordic). Upon destaining in distilled water, the gels were imaged using ChemiDoc imaging system (BioRad), and intensity of bands corresponding to αSyn was analyzed using Image Lab software (BioRad). The concentration of residual monomer was calculated based on the calibration curve made using the monomer standards of known concentrations.

### Quartz crystal microbalance analysis of fibril growth

Elongation of WT or KQ fibrils were measured by their immobilization on a QCM sensor (Biolin Scientific, Gothenburg, Sweden) and measuring changes in mass upon subsequent incubation with WT or KQ monomer solution (100). Sonicated fibrils (65 μL, 100 μM monomer equivalent) were mixed with 10 μL of 1 mg.mL^-1^ of Traut’s reagent (2-Iminothiolane, ThermoFisher) and spotted on the QCM sensor following 1 hour incubation at room temperature. Next, the solution was pipetted out and the chip surface was blocked by addition of 1% mPEG and incubation for 30 min. The sensor with immobilized fibrils was then thoroughly washed by miliQ H_2_O and dried under gentle nitrogen stream. The measurements were performed with a QSense Pro QCM-D instrument (Biolin Scientific, Gothenburg, Sweden) by measuring the elongation rate as change in the resonant frequency over time. The sample chamber equilibrated to 37 °C was filled automatically by 3 cell volumes (60 μL) of WT monomer (50 μM) and the measurement proceeded until stable linear slope was achieved. Next, the sensor was cleaned using buffer (50 mM Tris-HCl 150 mM KCl pH 7.4) and mutant monomeric solution was injected until a stable slope was achieved. The elongated rates of WT and mutant variants were measured as the slopes of the third overtone frequency after the first and second injections and used to calculate ΔΔG^ǂ^ according to the Equation M3. For the KQ fibrils, the sequence of WT and KQ mutant monomer addition was reversed.

### AFM analysis of the fibrils

Fibrils were diluted to 2.5 µM monomer equivalent concentration and 20 µL of the solution was deposited onto freshly cleaved mica substrates. Following 2min of incubation, the substrates were cleaned extensively with miliQ water and dried under nitrogen gas flow. All fibrils were imaged in tapping mode in air using a DriveAFM (Nanosurf, Liestal, Switzerland) using PPP-NCLAuD cantilevers (Nanosensors, Neuchatel, Switzerland). The amyloid fibrils were characterized by their apparent twist (assuming 2_1_ helical symmetry as described for Fm polymorphs in (71)) and height extracted using automated python script (59,72).

### Transmission electron microscopy of KQ4 and KQ6 fibrils

Samples were prepared on a glow discharged, formvar/carbon-coated 400 mesh grid for 20 seconds. The grids were then washed with two drops of double-distilled water and stained twice with 2% uranyl acetate. Excess stain was blotted, and the grids were air-dried for 30 minutes before imaging. A 200 kV Tecnai T20 G2 electron microscope (FEI, USA) was used to analyze the fibrils. Images were captured with a TVIPS XF415 CMOS 4K camera using TVIPS EMplify v0.4.5 software.

### Thermodynamic stability of fibrils

The thermodynamic stability of WT and mutant fibrils was measured in 50 mM Tris-HCl 150 mM KCl buffer pH 7.4 using chemical depolymerization and FIDA analysis of residual monomer as described in (72). In short, sonicated fibrils (40 μM monomer equivalent) were incubated in series of buffers containing increasing concentrations of urea (0-5 M) for 3 days at 25 °C. Samples were analyzed using by FIDA using the method described above. The monomer concentration was extracted from the elution profiles after correction for the viscosity at different urea concentrations as previously described. (72) The chemical depolymerization curves were analyzed using NumPyro to sample posterior distributions of the isodesmic model (Equation M5) parameters using the No U-Turn Sampler (NUTS). (73,74,101)

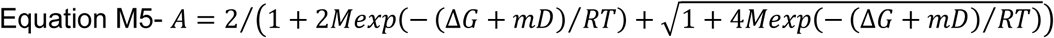

Where A is the area under the Gaussian peak from FIDA, M is the protein concentration in monomer equivalents, ΔG is the thermodynamic stability, m is the m-value, R the universal gas constant, and T is the temperature.

We fit a quadratic equation (Equation M6) to the correlation between denaturant m-value (m) and fibril diameter (d).

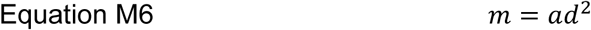

Where a is a proportionality constant corresponding to π/4T with T representing the average chain thickness (4.10^-10^ m for protein backbone). The quadratic dependence arises naturally from a “Clackson scroll” geometry, in which a rope or sheet is rolled into a cylindrical form. This configuration approximates the change in solvent-accessible surface area that occurs when a disordered protein chain assembles into a fibril. We assume that the *m*-value is directly proportional to this change in exposed surface area, consistent with the relationship observed for globular protein folding. (76)

### FoldX analysis of mutational changes on fibril stability

FoldX calculations were performed on a curated set of aSyn fibril structures selected from the Amyloid Atlas (v2024). (75) We curated the full set of aSyn structures by excluding (i) structures formed from conditions significantly different from physiological conditions used in this study, (ii) containing other compounds – such as lipids – or (iii) formed from mutant variants. Subsequently, the structures were aligned and manually investigated for similarity and structures with extensive overlap were removed to minimize bias from a single motif being represented multiple times. This curated set of 47 structures (Supplementary Table 9) represent the two major structure families recently identified in a meta-analysis of the full aSyn structure library, as well as the largest of the minor groups.(102)

FoldX calculations were performed using the FoldX4 suite as described in (59). (103) Briefly, before modelling any mutations in the structures, all PDB files were repaired using the REPAIRPDB command. Subsequent commands were only performed on the repaired structures. The ΔΔG upon mutagenesis was calculated using the BUILDMODEL command, ensuring that the mutation was introduced in all chains of the PDB file. The total ΔΔG is divided by the number of chains in the structure to evaluate a per chain ΔΔG. This was done for all possible single KQ mutations in the curated structure set.

### Clustering of mutational effects

Fibril stability, ΔΔG^ǂ^ values on KQ seeds, and cluster-specific coefficients from WLS regression of data from unseeded experiments, elongation experiments on WT at neutral pH, and mildly seeded experiments at low pH were log_2_-transformed. Biclustering of the resulting values was performed using seaborn.clustermap (Python 3.11) with hierarchical clustering applied to both assays and KQ variants, using the default settings of Euclidean distance and average linkage.

### Molecular dynamics simulations of αSyn monomers

All simulations were prepared and executed with the CALVADOS Python interface and through the publicly available Google Colab. (70) A single protein chain was simulated with both N-and C-terminal charges. The initial configuration was centered in a cubic box with a side length of 40 nm. Simulations were performed at 310.15 K, pH 7.5, and a range of ionic strengths (0, 5, 50, and 150 mM). Trajectories were saved every 100,000 integration steps (equivalent to 1 ns per saved frame). A total of 1000 frames were saved per replicate, corresponding to 100,000,000 integration steps and an aggregate production length of 1 μs. From each trajectory, ensemble observables including the radius of gyration (Rg), end-to-end distance (Ree), Flory scaling parameter (ν), and energy interaction map were computed automatically. Reported values represent averages across three independent replicates for each condition. Residue–residue contacts were calculated using a 9 Å cutoff applied to coarse-grained bead distances, excluding pairs with *i*–*j* ≤ 2 to remove bonded and next-nearest neighbour contributions. Contact probabilities and per-residue profiles were averaged across all three replicates. To connect simulations to experiment, Rg values were correlated with the aggregation half-times measured experimentally and averaged across salts for each ionic strength (for 150 mM, aggregation half-times at 100 and 200 mM NaCl, KCl, and NaI were averaged together).

## Authors contributions

A.K.B. supervised the work. A.K., S.F., R.K.N., and A.K.B. conceptualized the work. R.K.N., S.F., and A.K. designed the mutant primers, A.K. designed and carried out the experiments, analyzed the data, prepared the graphics, and wrote the manuscript. A.K. and S.F., carried out the seed amplification assay experiments, J.A.L and A.K. performed the QCM measurements, A.K., F.S., and C.F., expressed and purified the mutants, H.M.B. and A.S. carried out TEM analysis of the KQ fibrils, R.K.N. and C.F. wrote the python code for the analysis of urea depolymerization experiments. C.F. help with preparation of the graphics. J.F. and S.A. prepared the samples for seed amplification assay. All authors contributed to the preparation of the manuscript and agree with its content.

## Acknowledgements

A.K.B thanks the Novo Nordisk Foundation for funding (NNF17SA0028392 and NNF21OC0065495). This research was co-funded by the European Union (ERC CoG 101088163 EMMA to A.K.B.), Lundbeck foundation (grant number R366-2021-169 STADIC to A.K.B.). A.K. would like to acknowledge support through a Horizon MSCA individual postdoctoral fellowship (Grant number 101106115) for funding.

