## Supplementary Figures and Tables for "Probing the energy landscape of α-Synuclein amyloid fibril formation by systematic K-to-Q mutagenesis"

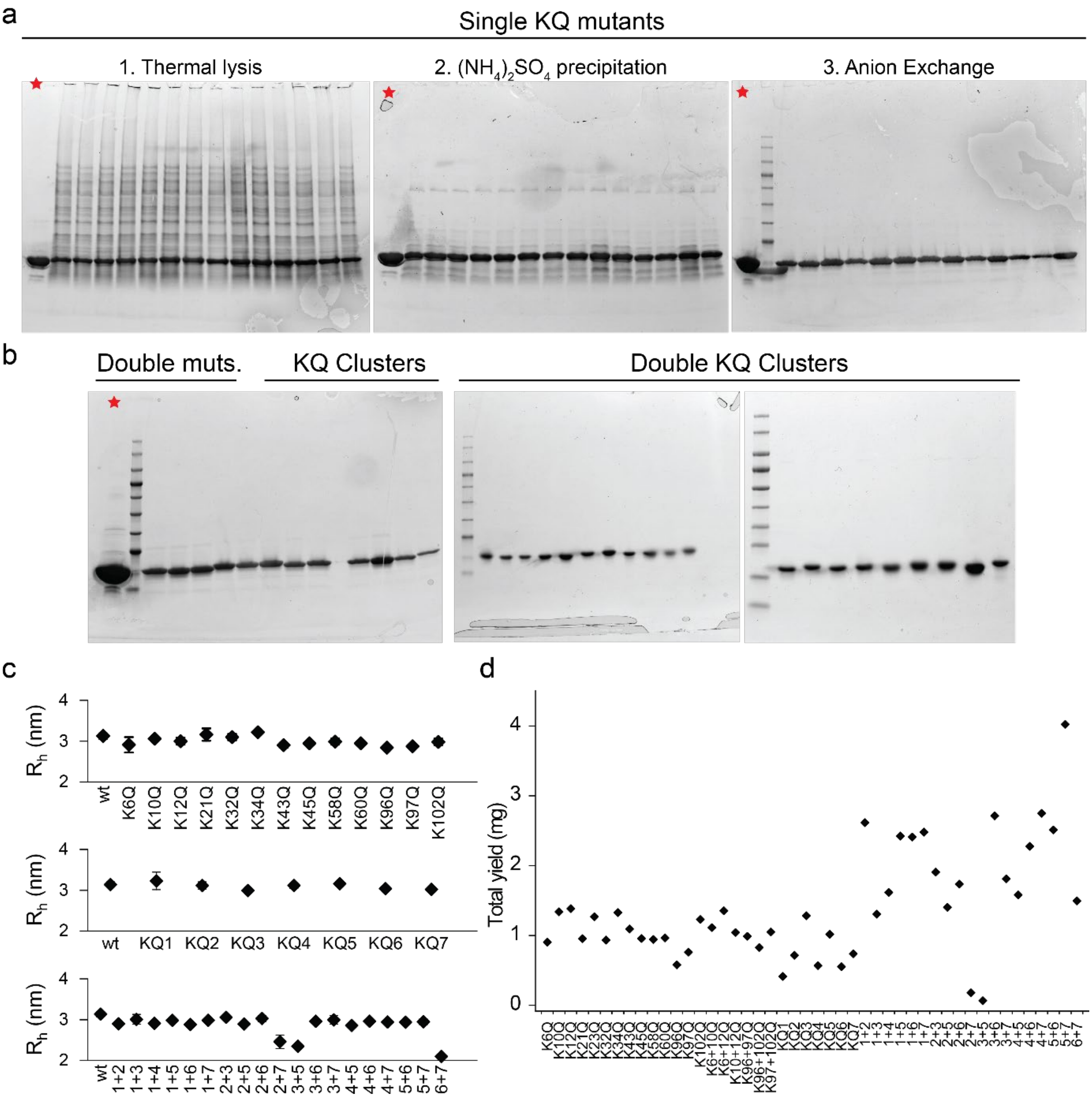


**Supplementary Figure 1. Small scale purification of mutant αSyn variants. a) SDS-PAGE analysis** of single-point KQ mutants after (1) thermal lysis and centrifugation of insoluble fraction, (2) precipitation by ammonium sulphate and subsequent centrifugation, and (3) anion exchange chromatography (AEC) in plate format. The control αSynWT monomer sample purified using standard protocol including size-exclusion is indicated by the star. **B) SDS-PAGE analysis** of double-point KQ mutants, KQ cluster, and double-KQ cluster variants after AEC. **C) Hydrodynamic radii** of the purified mutants determined using flow-induced dispersion analysis (FIDA). **D) Total yield** of all variants from 90 mL of LB cultures after AEC purification.

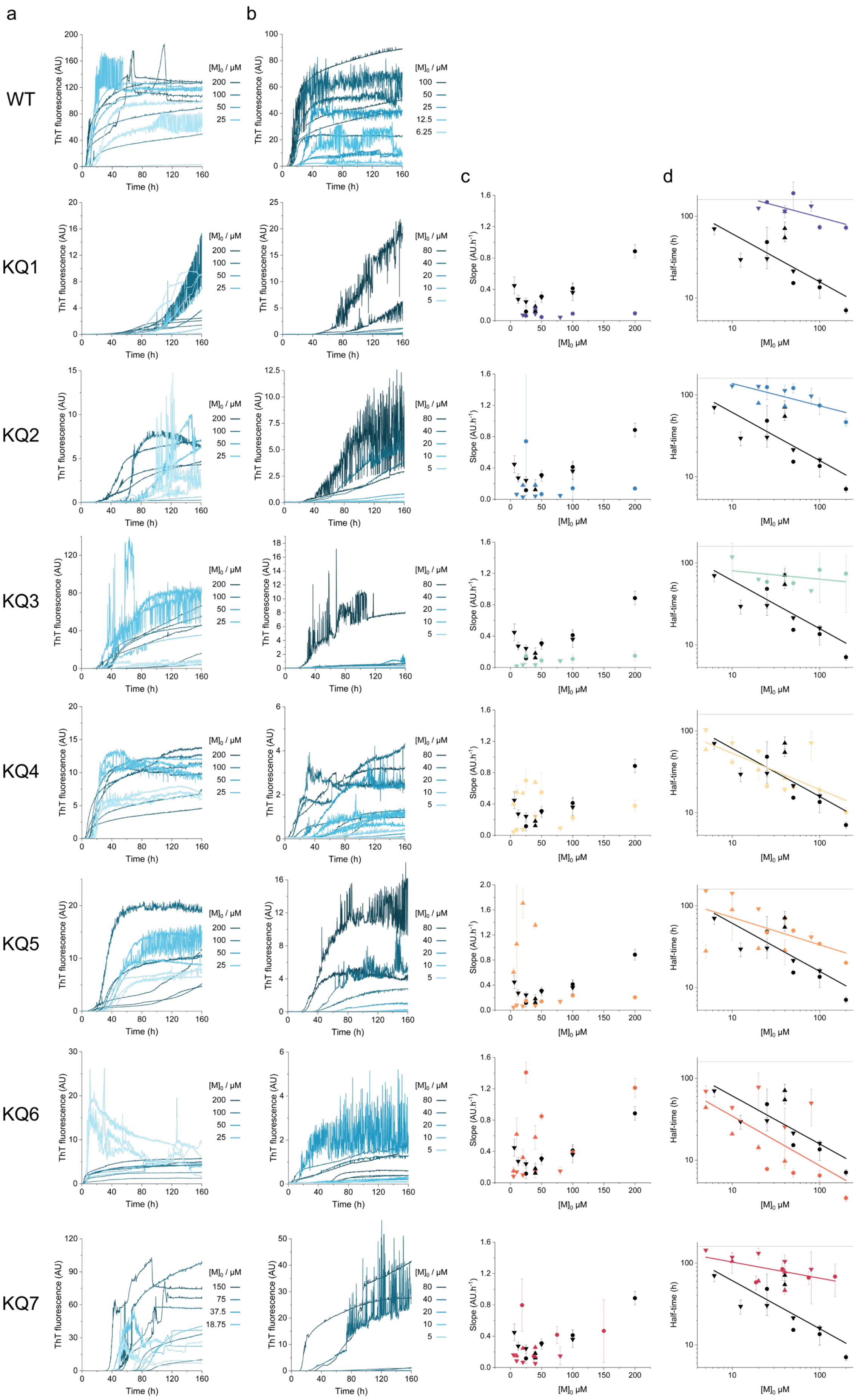
a
b
c
d
WT
KQ1
KQ2
KQ3
KQ4
KQ5
KQ6
KQ7
ThT fluorescence (AU)
Time (h)
[M]$_0$ / µM
200
100
50
25
12.5
6.25
80
40
20
10
5
150
75
37.5
18.75
Slope (AU.h$^{-1}$)
Half-time (h)
[M]$_0$ µM

**Supplementary Figure 2. De novo aggregation of WT and KQ cluster mutants at varying initial monomer concentrations.** Two independent experiments carried out in **(a) 10 mM NaP pH 7.5 with 150 mM NaCl** and **(b) 50 mM Tris-HCl pH 7.5 with 150 mM KCl.** The initial monomer concentrations used are provided in the legends. **(c) The apparent aggregation rates** (slopes) obtained from fitting the curves in a (circles) and b (triangles) as functions of initial monomer concentration. The points and error bars correspond to the mean and SD from a triplicate measurement, respectively. **(d) Scaling of aggregation half-times with initial monomer concentration** obtained from fitting the curves in a (circles) and b (triangles). The points and error bars correspond to the mean and SD from a triplicate measurement, respectively. The solid lines represent linear fit to derive the slope of the scaling. Dashed lines indicate time of the experiment end. The WT data in each plot of (c) and (d) correspond to the same dataset to facilitate easier comparison with the mutants.

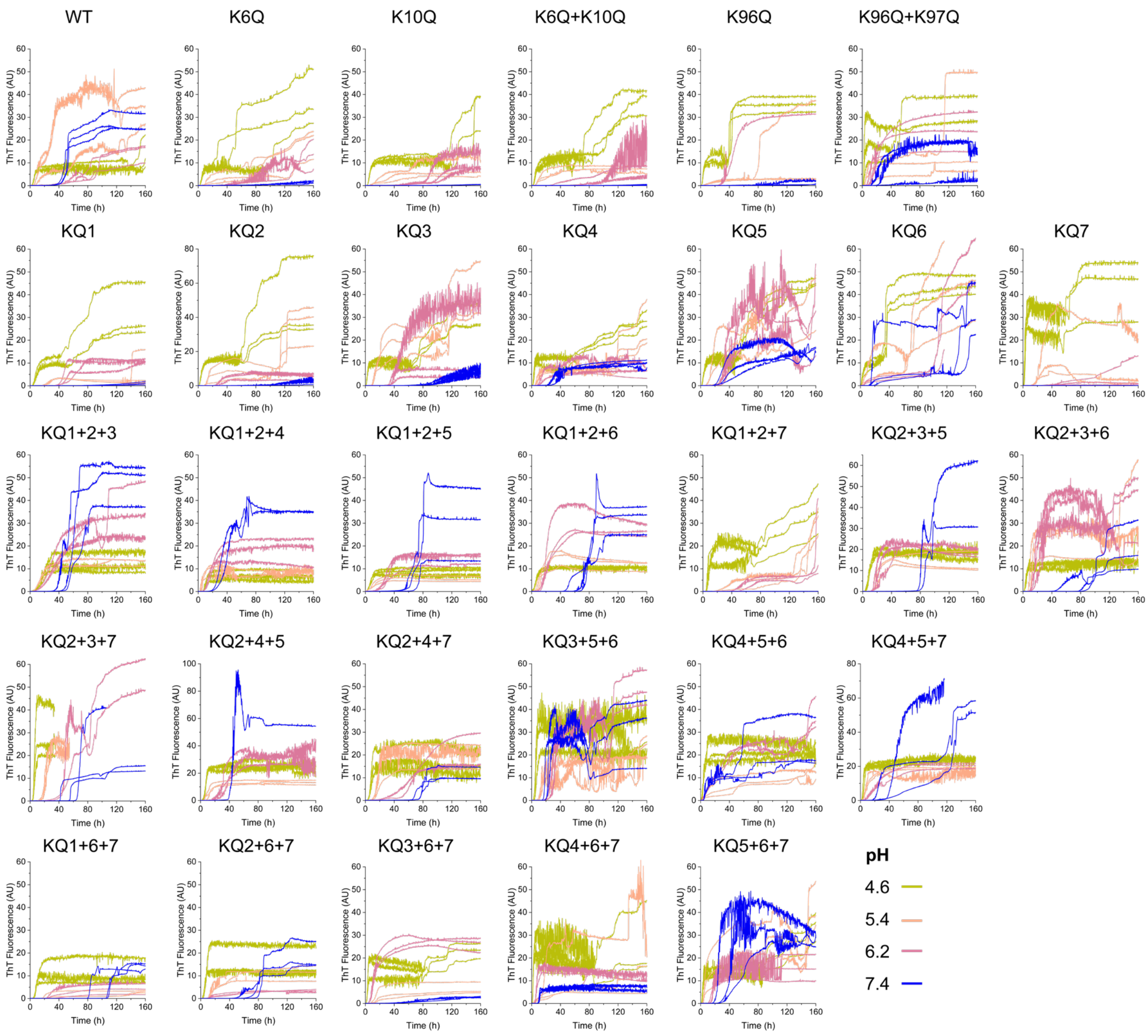


**Supplementary Figure 3. De novo aggregation of WT and KQ mutants in 50 mM citrate buffer at varying pH.** Two independent experiments were carried out with WT, single-point mutants, partial KQ cluster mutants (K6Q+K10Q, K96Q+K97Q, first row), KQ cluster mutants (second row), and triple cluster mutants (i.e., variants with three lysine clusters mutated to glutamines, e.g., KQ1+2+3 = 7-point mutant K6Q+K10Q+K12Q+K21Q+K23Q+K32Q+K34Q, third to fifth row). The pH of the buffers was 4.6 (green), 5.4 (orange), 6.2 (pink), and 7.4 (blue).

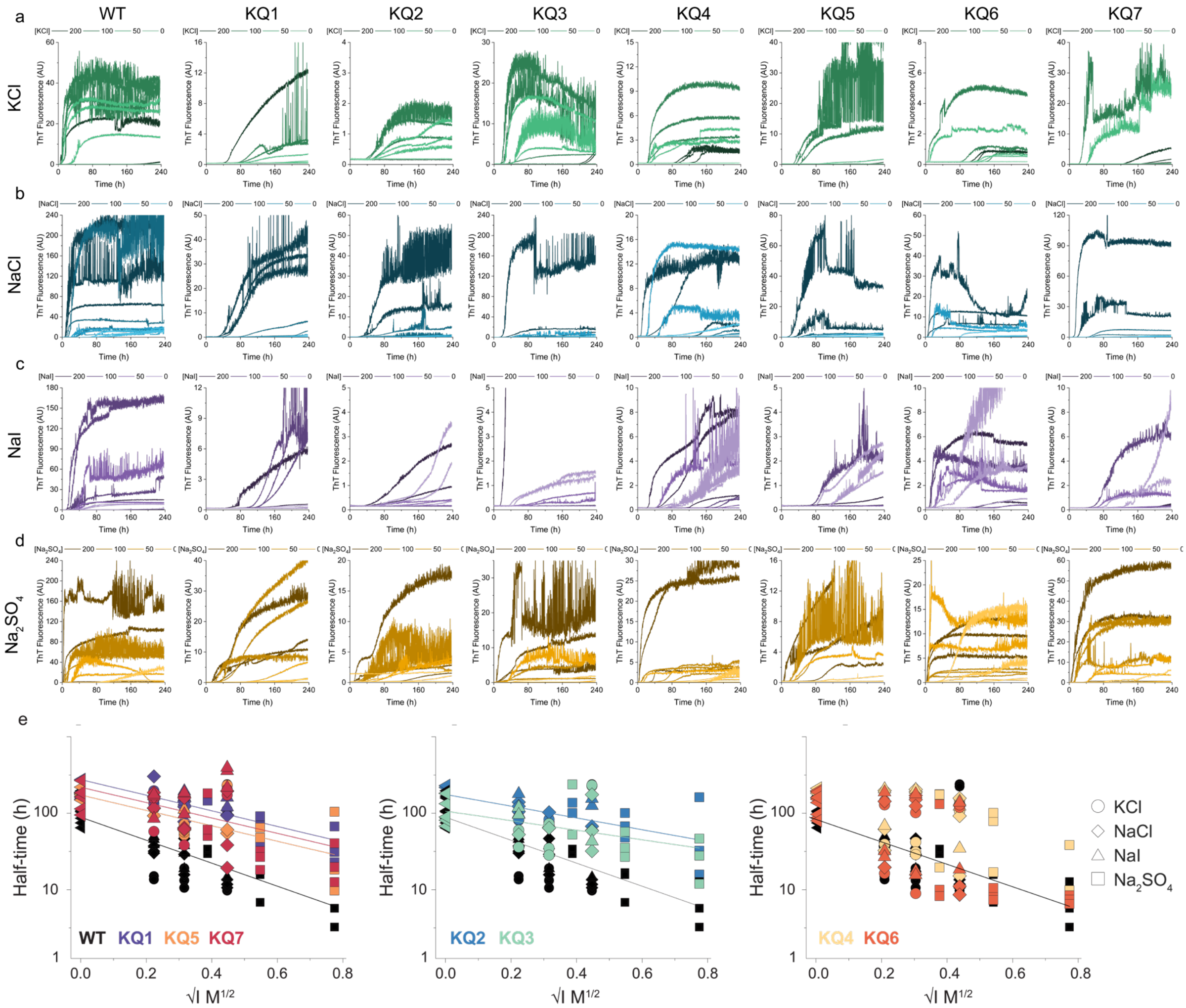


**Supplementary Figure 4. De novo ThT aggregation assay of WT and KQ cluster mutants in 10 mM sodium phosphate buffer with varying concentrations of (a) KCl, (b) NaCl, (c) NaI, and (d) $Na_2SO_4$.** The concentrations of salts are color-coded by different tones of the respective colours as shown in the legends above the plots**. (e) Debye plots of the log-transformed aggregation half-times** as a function of the square root of ionic strength (salt only). The points correspond to mean values from a triplicate measurement shown in the plots above (left triangle – no salt, circle – KCl, diamond – NaCl, triangle – NaI, square – $Na_2SO_4$). The lines represent linear fits to the data.

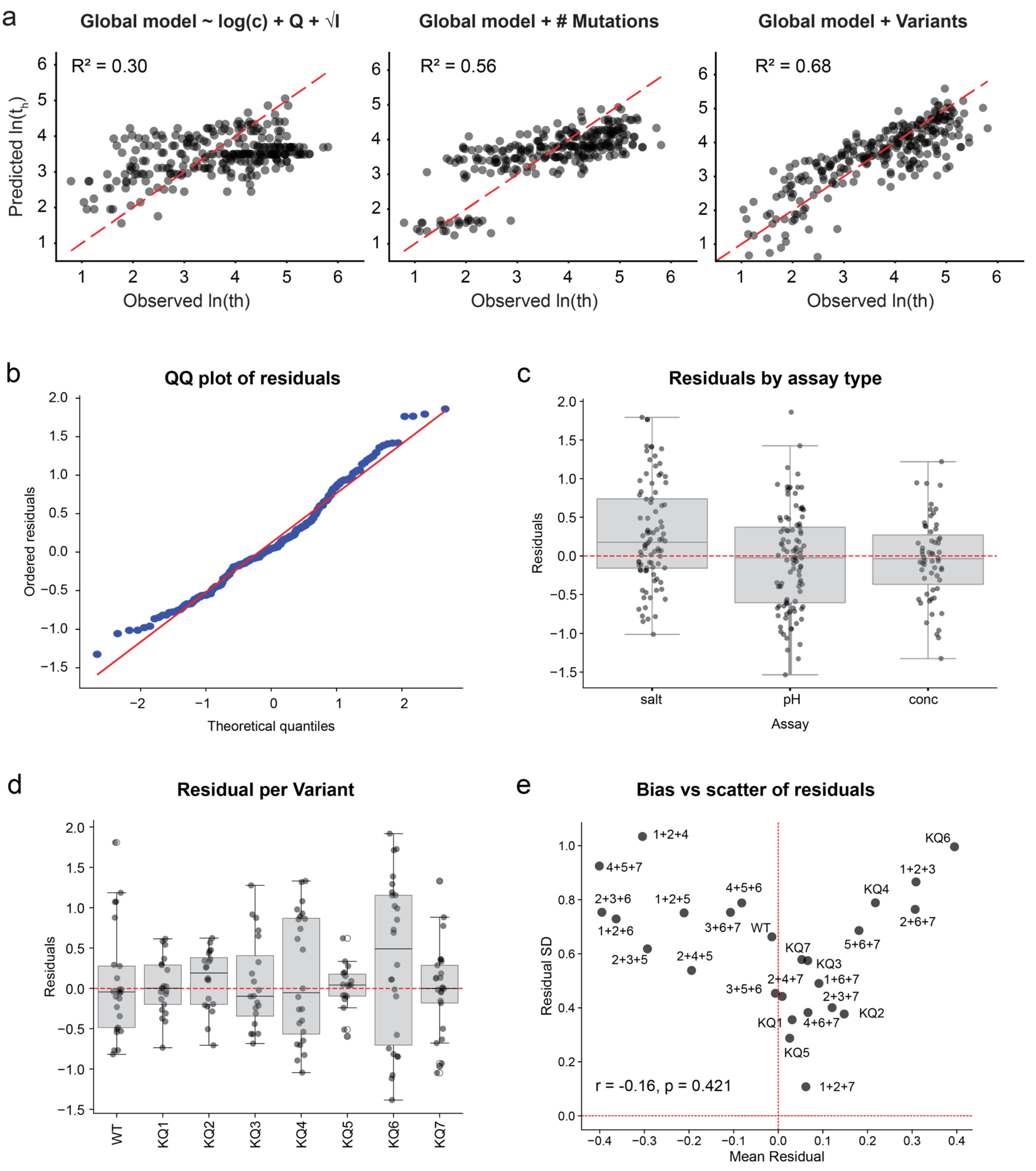


**Supplementary Figure 5. Global model of de novo αSyn aggregation. (a) Model performance with various parameters.** (left) Initial model involving only the global terms charge (Q), initial monomer concentration (c) and ionic strength (I). (right) Model including global terms and number of mutations as parameters. (right) Final model involving global terms and variant-specific slopes. **(b) Quantile–quantile (QQ) plot of model residuals.** The plot compares the distribution of standardized residuals from the final model against the theoretical quantiles of a normal distribution. Points falling along the red dashed line indicate approximate normality. **(c) Residuals by assay type.** The points correspond to residuals from the model (observed-predicted) group by the assay type. The box represents the interquartile range (IQR), the horizontal line shows the median, whiskers extend to 1.5×IQR, and the red dashed line marks the perfect model fit (zero residuals). **(d) Residuals by variant.** The points correspond to residuals from the model (observed-predicted) group by the assay type. The box represents the interquartile range (IQR), the horizontal line shows the median, whiskers extend to 1.5×IQR, and the red dashed line marks the perfect model fit (zero residuals). **(e) Correlation between mean and SD of the residuals.** The lack of correlation (Pearson's r and p-value shown in the graph) indicate no systematic deviations from the fit.

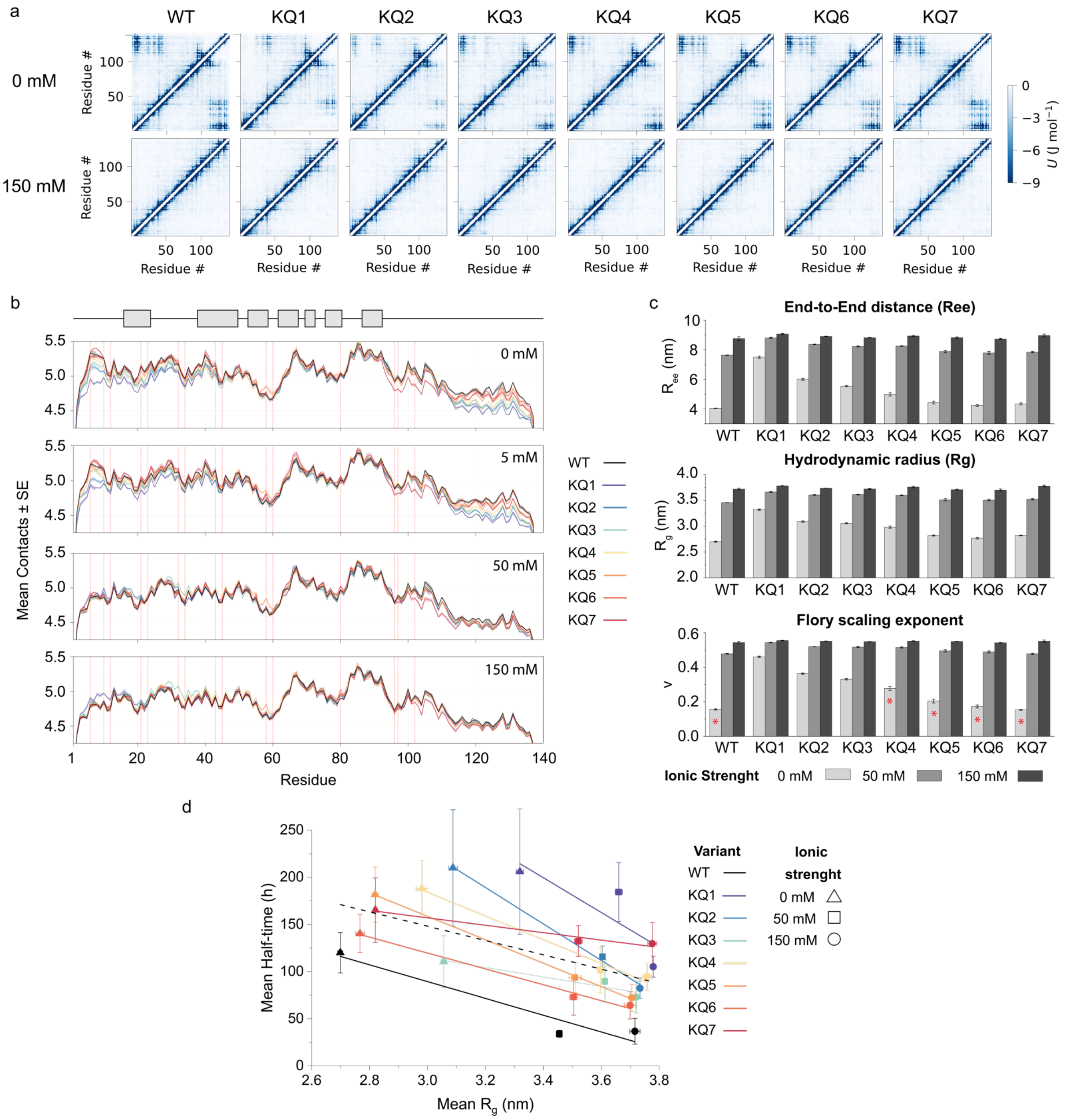


**Supplementary Figure 6. Molecular dynamics simulations of αSyn variants. a) Interaction energy maps** derived from simulations of single protein chains at 0 (top) or 150 mM (bottom) ionic strength. The interaction strength is color-coded according to the legend. **b) Sequence profiles of average intra-chain residue contacts** for WT and KQ cluster variants, excluding the neighbouring residues. The contact cut-off value was 9 Å. The plots correspond to the average from three simulation replicates and SE. The monomer secondary structure within fibril plane is provided on top for illustration. **c) Global scaling properties** derived from the MD simulations at varying ionic strengths. The values correspond to means ± SE from three simulation replicates. The red asterisk at the bottom graph denotes values resulting from poor fitting of the $R_{ij}=R_0|i-j|^3$ power law. **d) Correlations between experimental aggregation half-times and MD-derived compactness** (end-to-end chain distance). The solid and dashed lines correspond to linear fits to the data from individual mutants and their average, respectively.

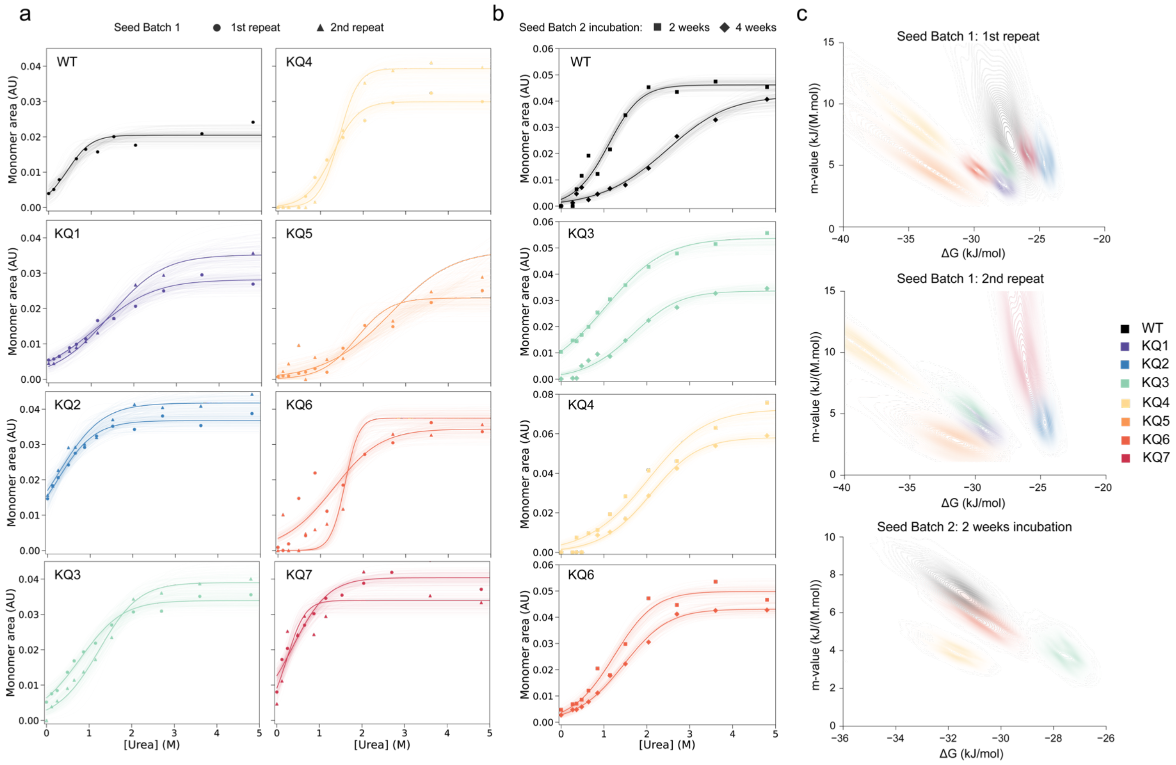


**Supplementary Figure 7. Chemical depolymerization of fibrils formed by WT and KQ cluster variants of αSyn. (a) and (b) Chemical depolymerization of different fibril batches.** Data points (area under the curve obtained from FIDA measurements corresponding to monomeric αSyn) and their fits to the isodesmic model using MCMC sampling of solutions (thin lines correspond to the 100 random solutions out of 2000 total, best fit is shown in bold). **(c) Correlations of the m-values and ΔG** obtained from the fitting. The possible solution space is represented by kernel density estimation color-coded according to the mutants.

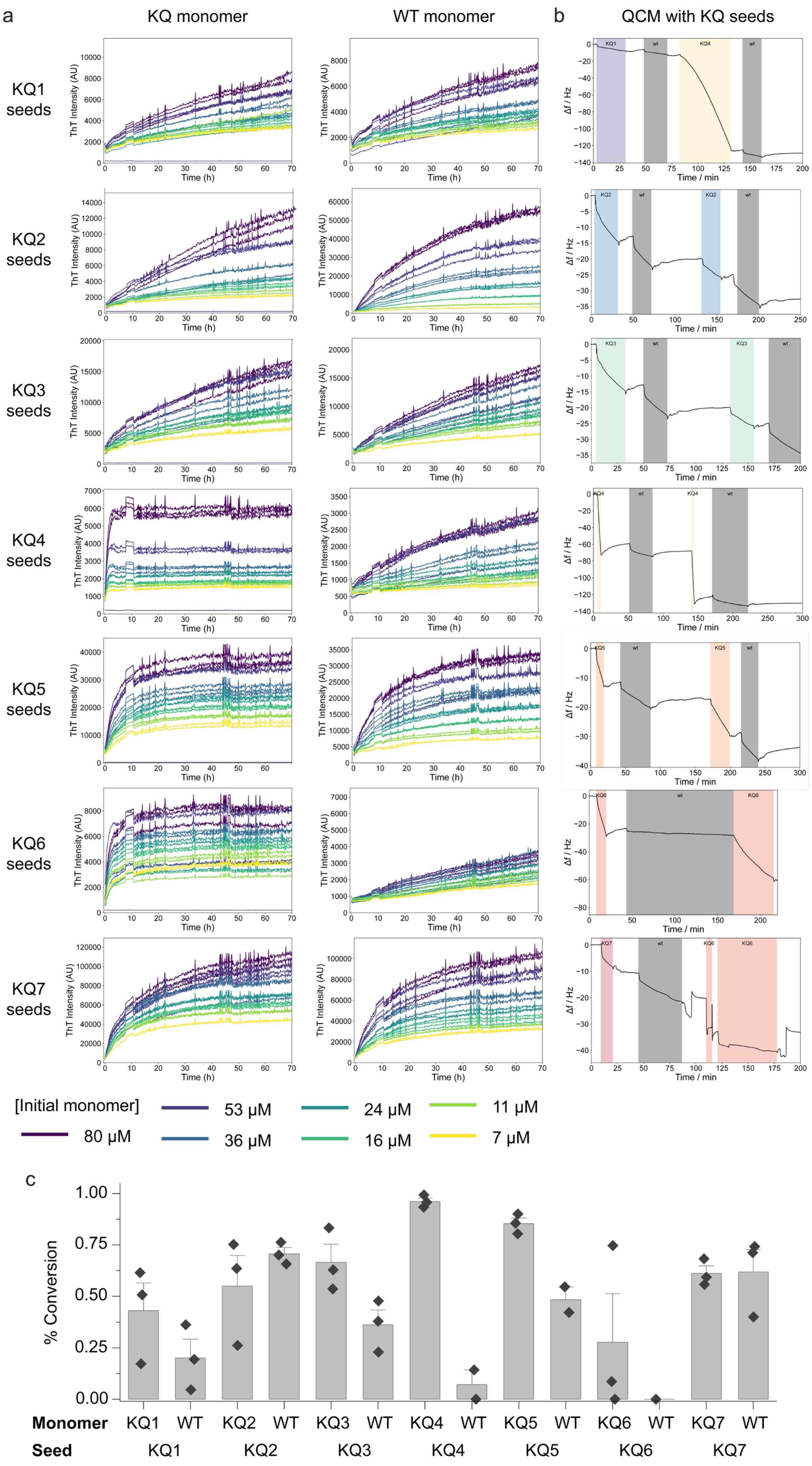
a
KQ monomer
WT monomer
b
QCM with KQ seeds
KQ1 seeds
KQ2 seeds
KQ3 seeds
KQ4 seeds
KQ5 seeds
KQ6 seeds
KQ7 seeds
ThT Intensity (AU)
Time (h)
Δf / Hz
Time / min
[Initial monomer]
80 μM
53 μM
36 μM
24 μM
16 μM
11 μM
7 μM
c
% Conversion
Monomer
KQ1 WT KQ2 WT KQ3 WT KQ4 WT KQ5 WT KQ6 WT KQ7 WT
Seed
KQ1 KQ2 KQ3 KQ4 KQ5 KQ6 KQ7

**Supplementary Figure 8. Aggregation kinetics using KQ seeds. (a) Aggregation kinetics monitored by ThT.** The mutant (left column) or WT (right column) monomers were seeded by 2.5 µM sonicated KQ fibrils. The color-scheme highlights different initial concentrations of monomers provided in the legend below. **(b) Aggregation kinetics monitored using quartz crystal microbalance (QCM).** The KQ seeds immobilized on a QCM sensor were first elongated by their respective monomer (homogeneous elongation, coloured boxes, followed by elongation by WT monomer (grey boxes), with thorough washing steps in between. The elongation rates of the mutant and WT were obtained by fitting the linear part of the change in the frequency of 3$^{rd}$ harmonic overtone (black line) from the first and second injection, respectively. **(c) Residual monomer analysis.** Soluble monomer at the end of aggregation reaction was quantified either by SDS-PAGE analysis or UV absorbance (ThT free samples, diamonds). The y-axis represents percentage of initial monomer concentration converted to fibrils. The grey bars and error bars represent mean ± SE from three independent measurements.

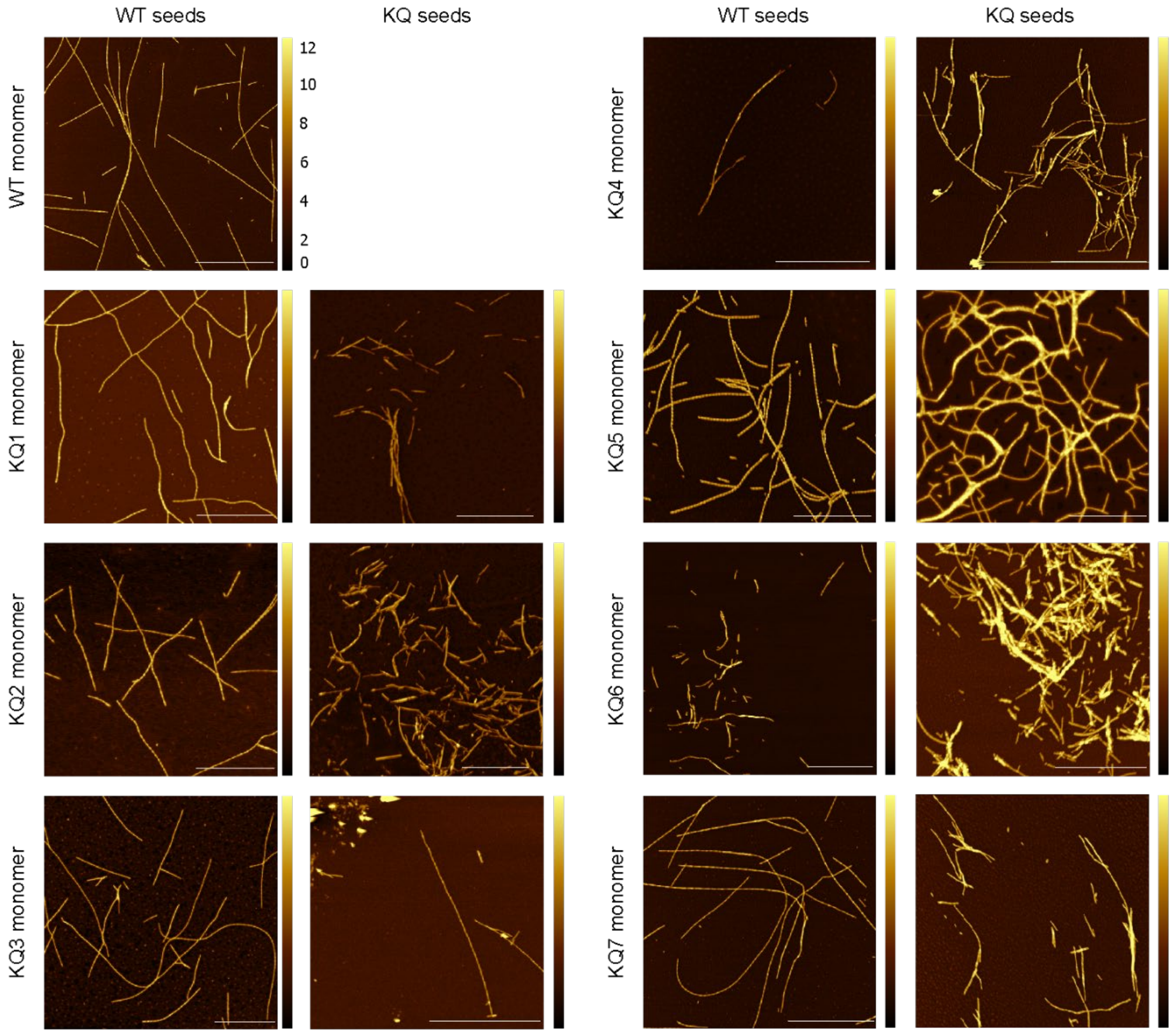


**Supplementary Figure 9. Fibril morphology of WT and mutant fibrils.** AFM images of the fibrils formed by aggregation of the αSyn monomers in the presence of pre-formed WT or KQ fibrils. The fibril heights in nm are indicated by the color-coding. The scale bars represent 2 μm.

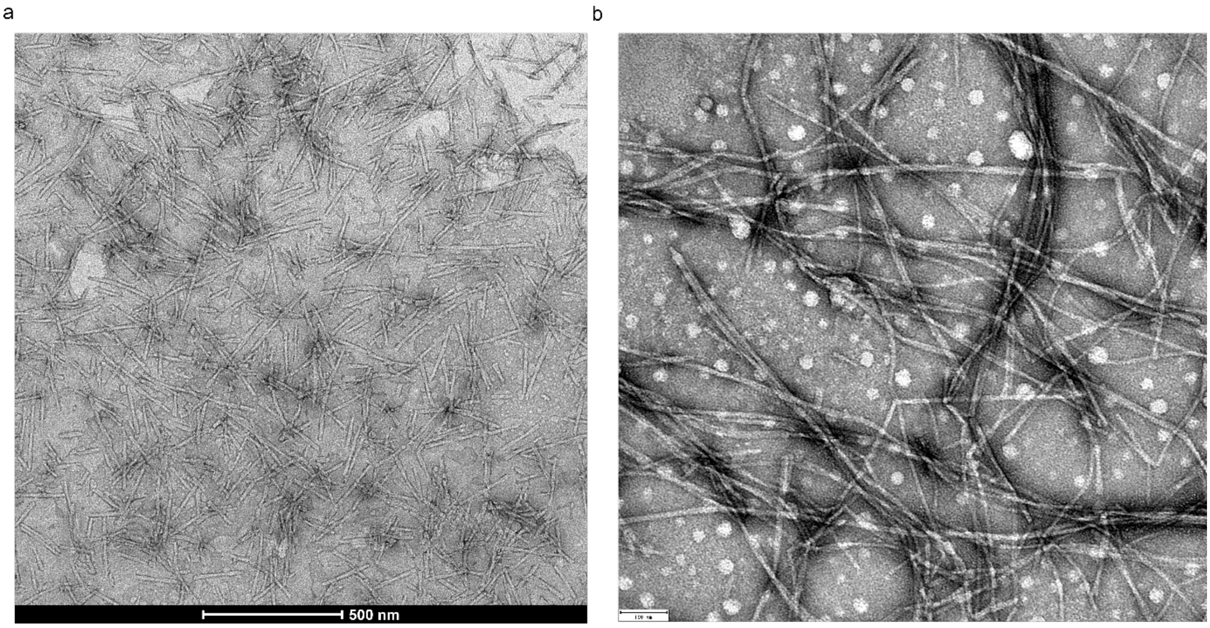


**Supplementary Figure 10. Representative TEM images of (a) KQ4 and (b) KQ6 fibrils.** The scale bars in a and b correspond to 500 and 100 nm, respectively.

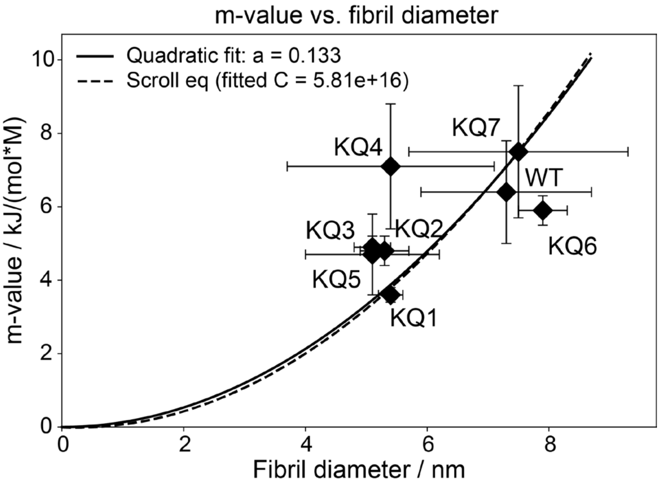


**Supplementary Figure 11. Correlation between AFM derived heights (fibril diameters) and urea m-values from depolymerization experiments.** The data correspond to the means ± SD of fibril heights obtained from the AFM analysis and means ± SD of m-values from chemical depolymerization of the fibrils. The solid and dashed lines represent fits to simple quadratic equation (Equation M6) and Clarkson scroll model approximation, respectively. C is the proportionality constant between the m-value and change in solvent exposed area based on the model (based on chain thickness T of 0.4 nm).

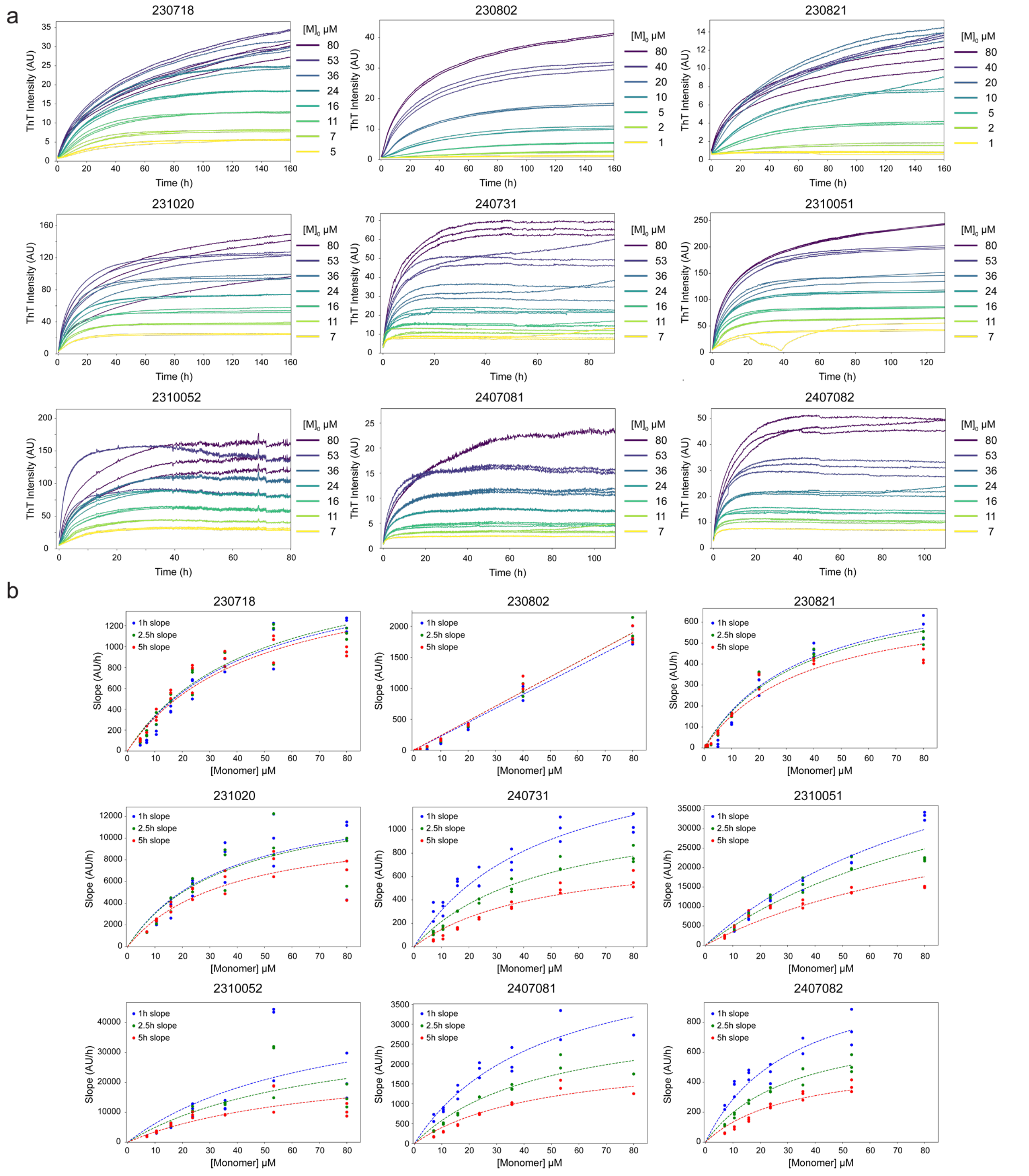


**Supplementary Figure 12. Reproducibility of the seeded aggregation assays. (a) Raw data from ThT aggregation experiments** using WT monomer and pre-formed WT Fm seeds. The initial monomer concentrations used are indicated in the legend. (**b) Initial rate analysis.** The initial rates were calculated from the data shown in (a) in the time range of 0 – 1 h (blue), 0 – 2.5 h (green), and 0 - 5 h (red). The fits of the initial rates as a function of initial monomer concentration to the equation describing saturating elongation are shown as dotted lines. The saturation concentrations ($K_e$) were fitted globally whilst the maximal rates were treated as free parameters within each dataset.

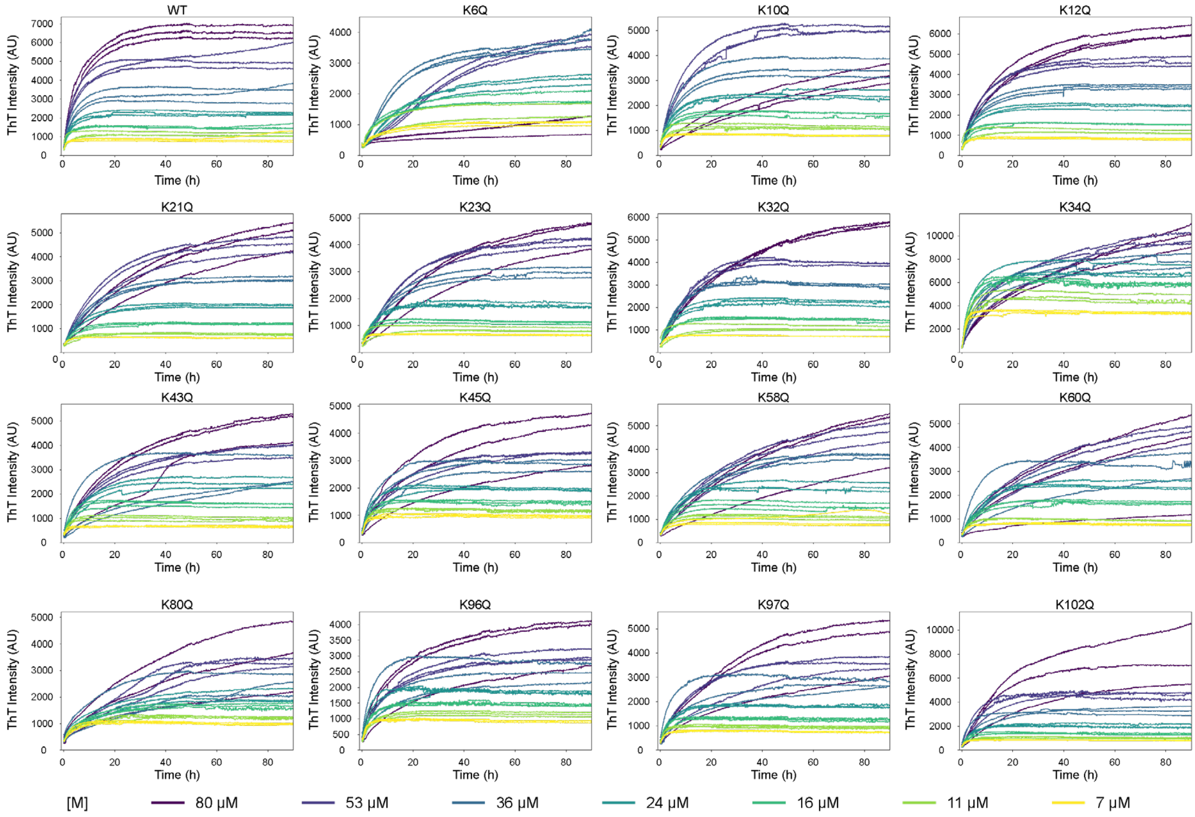


**Supplementary Figure 13. Seeded aggregation kinetics of single-point mutants on WT Fm seeds** followed using ThT kinetics. The initial rates as a function of initial concentration were used to derived ΔΔG‡ of elongation (one of the points in Figure 4c). The initial monomer concentrations are shown below the graphs.

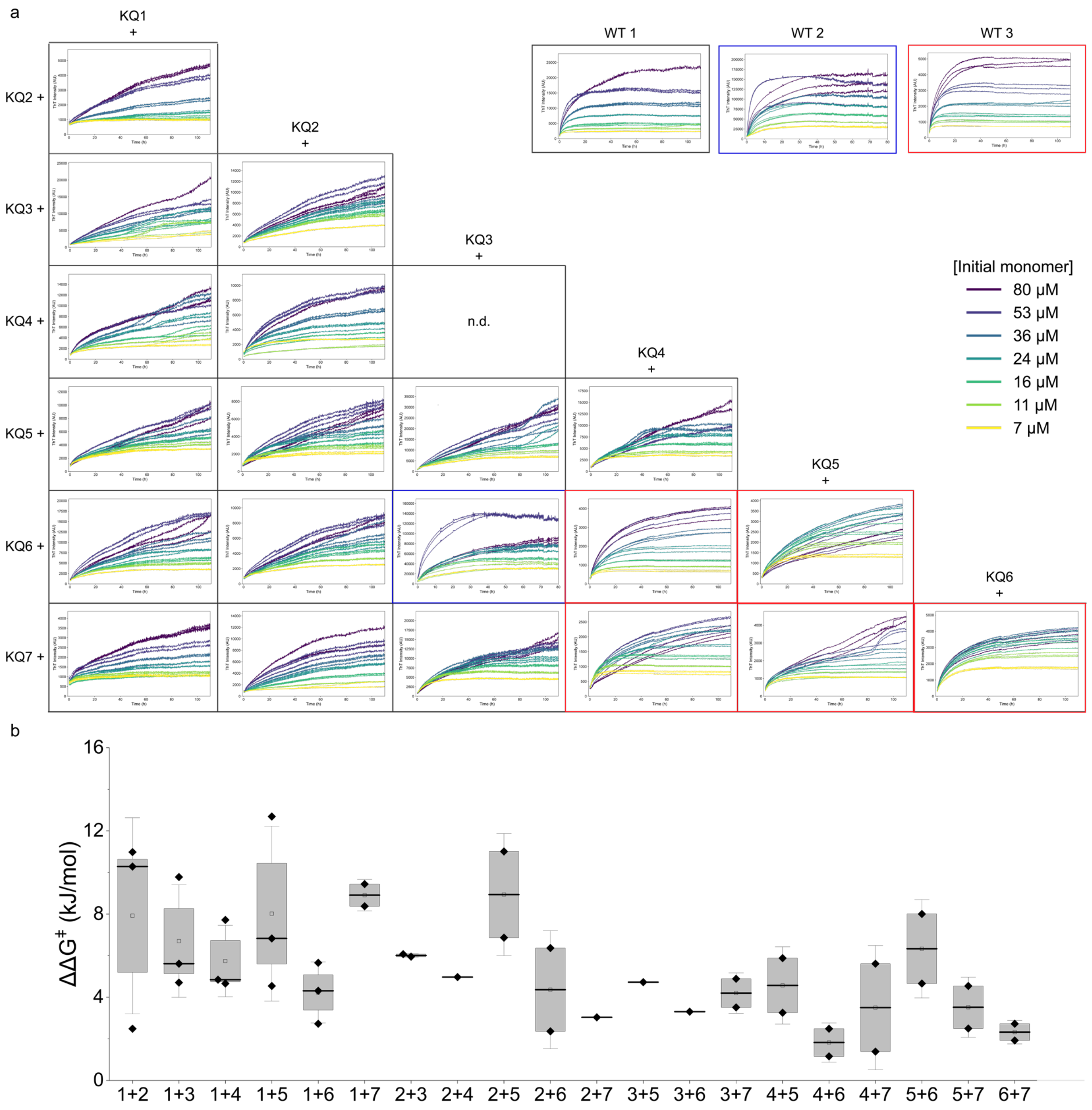


**Supplementary Figure 14. Seeded aggregation kinetics of double KQ clusters on WT Fm seeds. a) Raw ThT kinetics.** The data correspond to the variants carrying combination of the KQ cluster mutations indicated on top of each column and on the left side of each row (e.g., top left: KQ1+KQ2 = K6Q+K10Q+K12Q+K21Q+K23Q). The coloured graph borders indicate different experiments with corresponding WT datasets shown in the top right corner. **b) Change of the energy barrier of elongation (ΔΔG‡) for the double KQ cluster mutants** on the WT Fm seeds. The diamonds represent values calculated from individual experiments. The box and the whiskers represent 1 SE and 1 SD of the values, respectively. The mean and median are depicted by squares and lines, respectively.

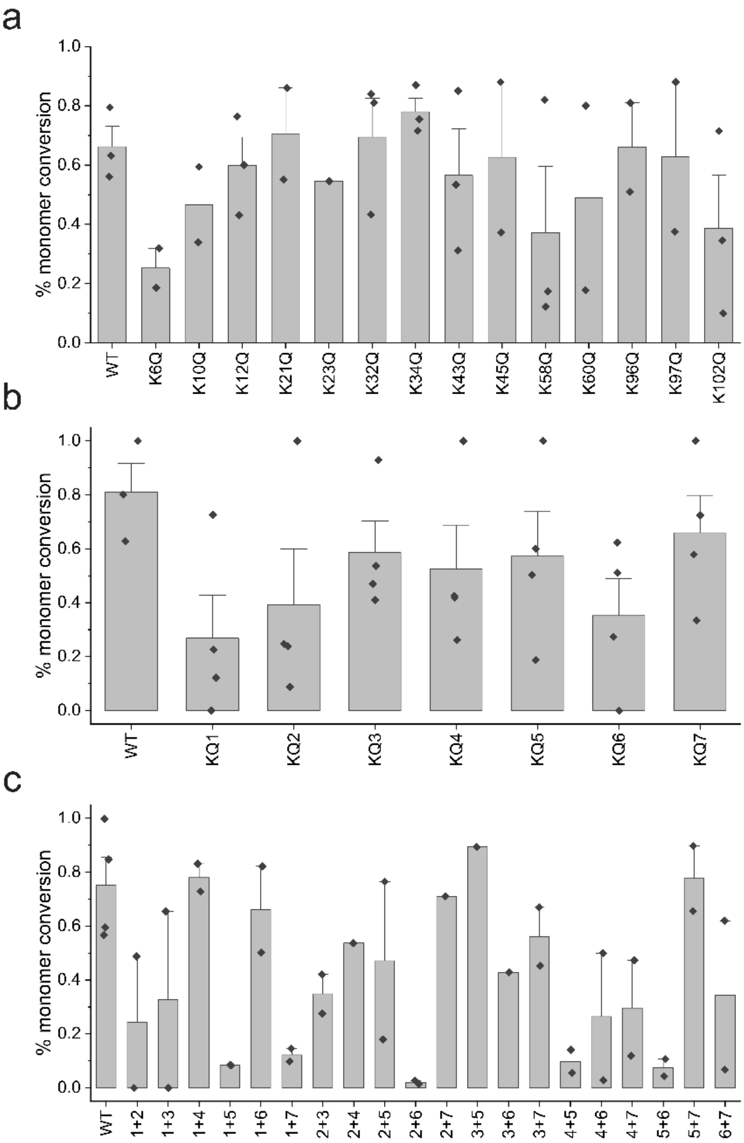


**Supplementary Figure 15. Analysis of residual soluble monomer concentration** at the end of the aggregation assays seeded by WT Fm polymorph. The points corresponding to the fraction of monomers converted to fibrils were obtained by SDS-PAGE, UV absorbance, and FIDA analyses or their combination from each elongation datasets with **A)** single-point mutants, **B)** KQ cluster variants, and **C)** double KQ cluster variants. The bar and error bar represent mean + SE, respectively.

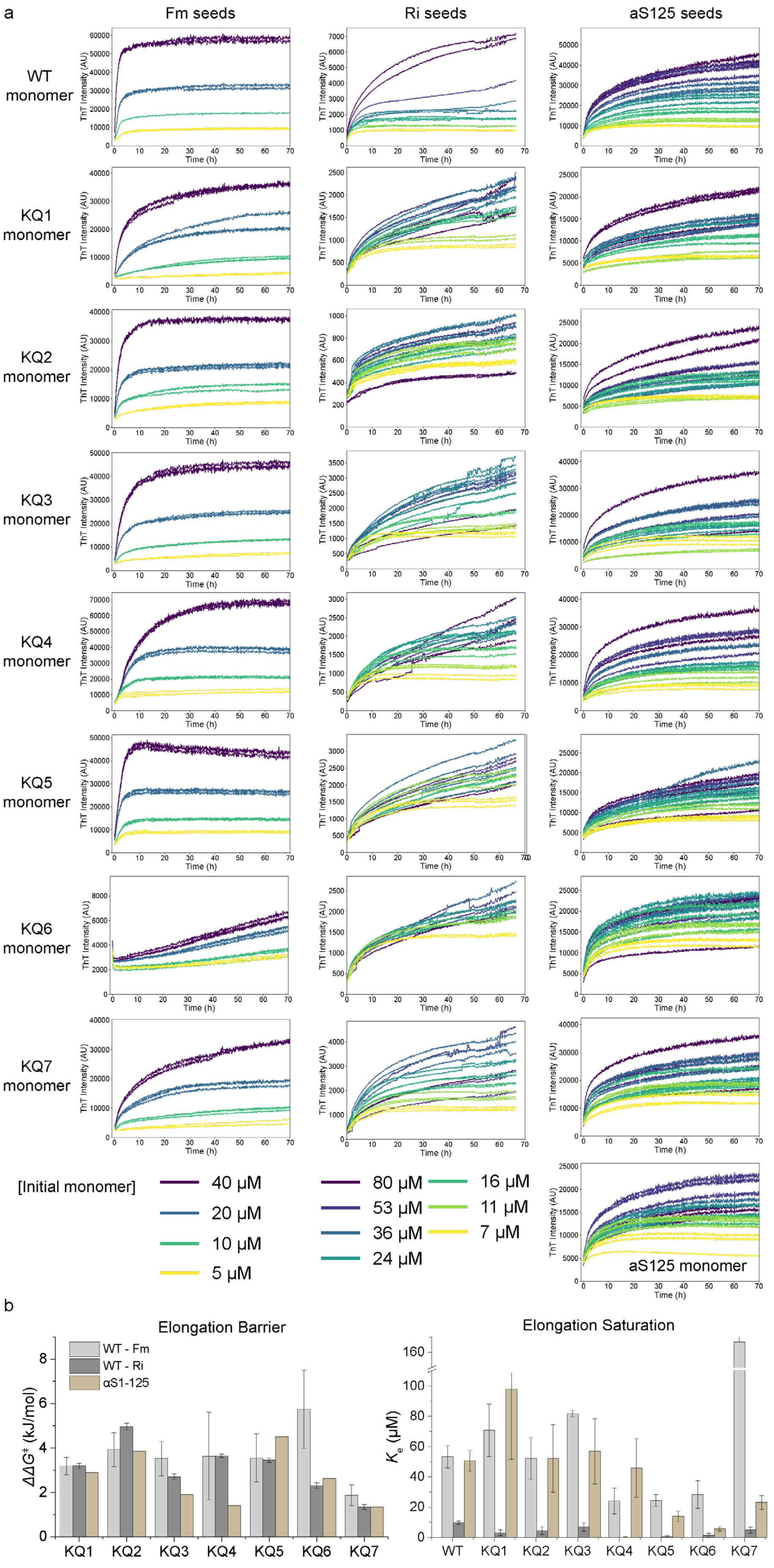
a
Fm seeds
Ri seeds
aS125 seeds
WT monomer
KQ1 monomer
KQ2 monomer
KQ3 monomer
KQ4 monomer
KQ5 monomer
KQ6 monomer
KQ7 monomer
ThT Intensity (AU)
Time (h)
[Initial monomer]
40 μM
20 μM
10 μM
5 μM
80 μM
53 μM
36 μM
24 μM
16 μM
11 μM
7 μM
aS125 monomer
b
Elongation Barrier
WT - Fm
WT - Ri
αS1-125
ΔΔG‡ (kJ/mol)
KQ1
KQ2
KQ3
KQ4
KQ5
KQ6
KQ7
Elongation Saturation
Ke (μM)
WT

**Supplementary Figure 16. (a) Example of aggregation kinetics of KQ clusters** seeded by WT Fm (left column), WT Ri polymorphs (middle column) and fibrils prepared from C-terminally truncated aS125 (right column). The initial monomer concentrations used in the experiments with Fm, and Ri or aS125 are shown below the graph. **(b) (left) Comparison of ΔΔG‡ and $K_e$ values of KQ variants on different polymorphs.** The Fm values represent mean ± SE of independent datasets corresponding to the values in Figure 3c. The values for Ri and aS125 are values and SE obtained from fitting of the initial rate analysis of datasets shown in a. The values are provided in Supplementary Table 3.

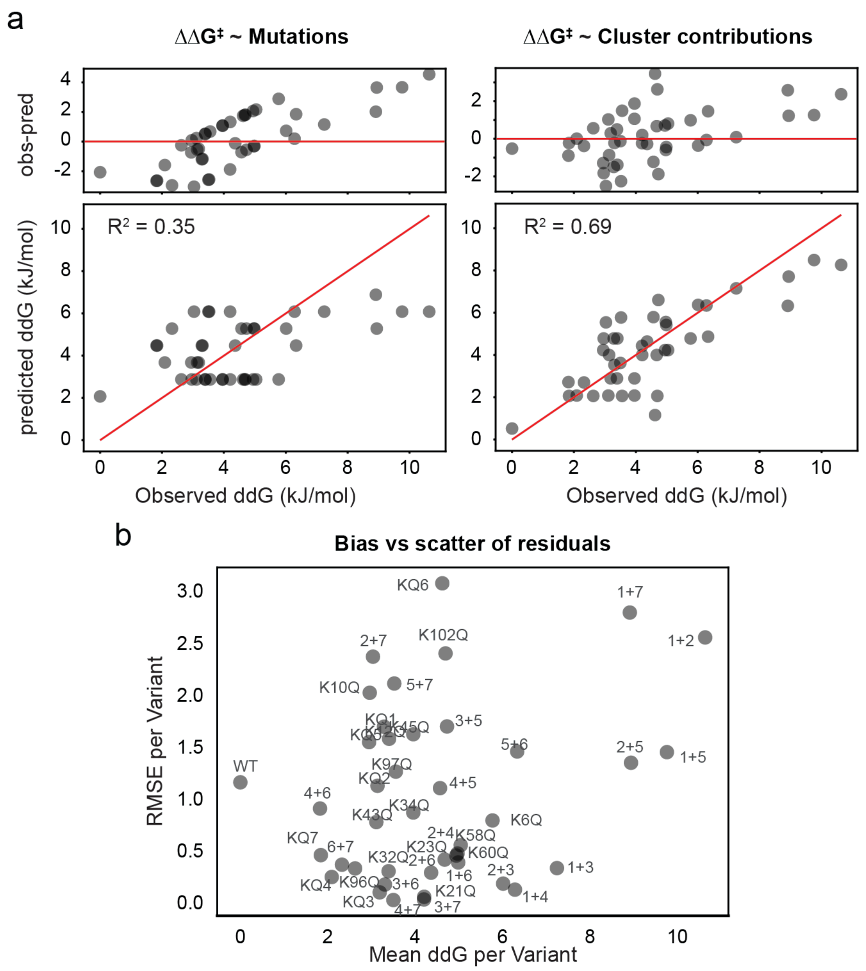


**Supplementary Figure 17. Modelling of relative growth rates (ΔΔG‡) by linear regression** as a function of number of mutations (**a, left**), or mean cluster contributions (**a, right**). Individual points correspond to the mean values from Figure 4c with 1:1 line depicted in red. Residuals (observed-predicted) are shown on top. **(b) Correlation between mean and SD of the residuals** from the model with mean cluster contributions.

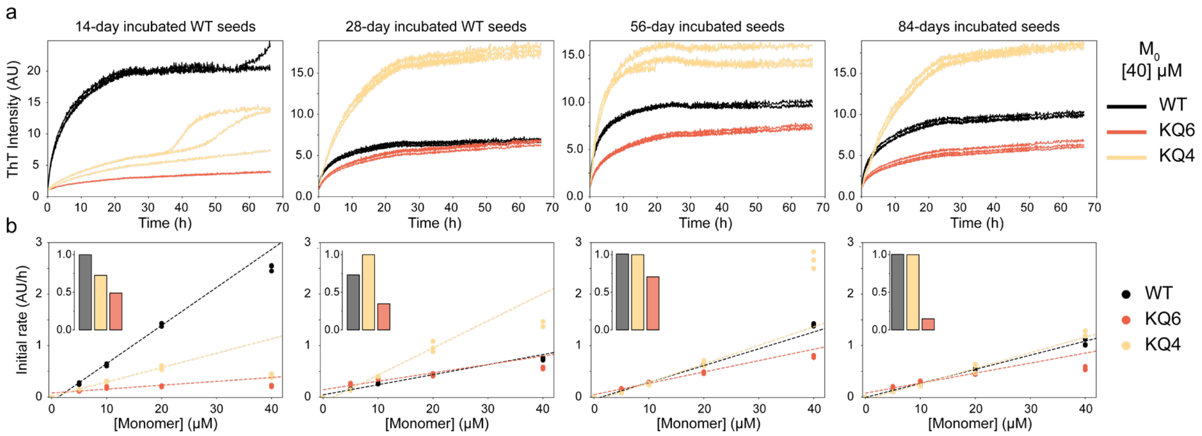


**Supplementary Figure 18. Time-dependent changes in Fm seeding efficiency.** For this experiment, WT monomer was incubated at the Fm conditions (Table 4) for varying time (14, 28, 56, and 84 days), after which the fibrils were centrifuged, resuspended in fresh buffer, sonicated and used as seeds in the same experiments. **a) ThT aggregation kinetics** of 40 μM WT (black), KQ4 (yellow), and KQ6 (red) monomers in the presence of 2.5 μM WT Fm seeds that were formed by (from left to right) 14, 28, 56, and 84 days of incubation at 37 °C. **b) The initial rate analysis.** The initial rates (0 - 2.5 h) plotted as a function of monomer concentration were fitted by a linear function (dotted lines) in the monomer range of 0-20 μM. Conversion of the monomers to aggregates was assessed from the SDS-PAGE analysis at the end of the kinetics shown in (a) and is indicated by the bar graphs as the inset of each plot (grey – WT, yellow – KQ4, red – KQ6).

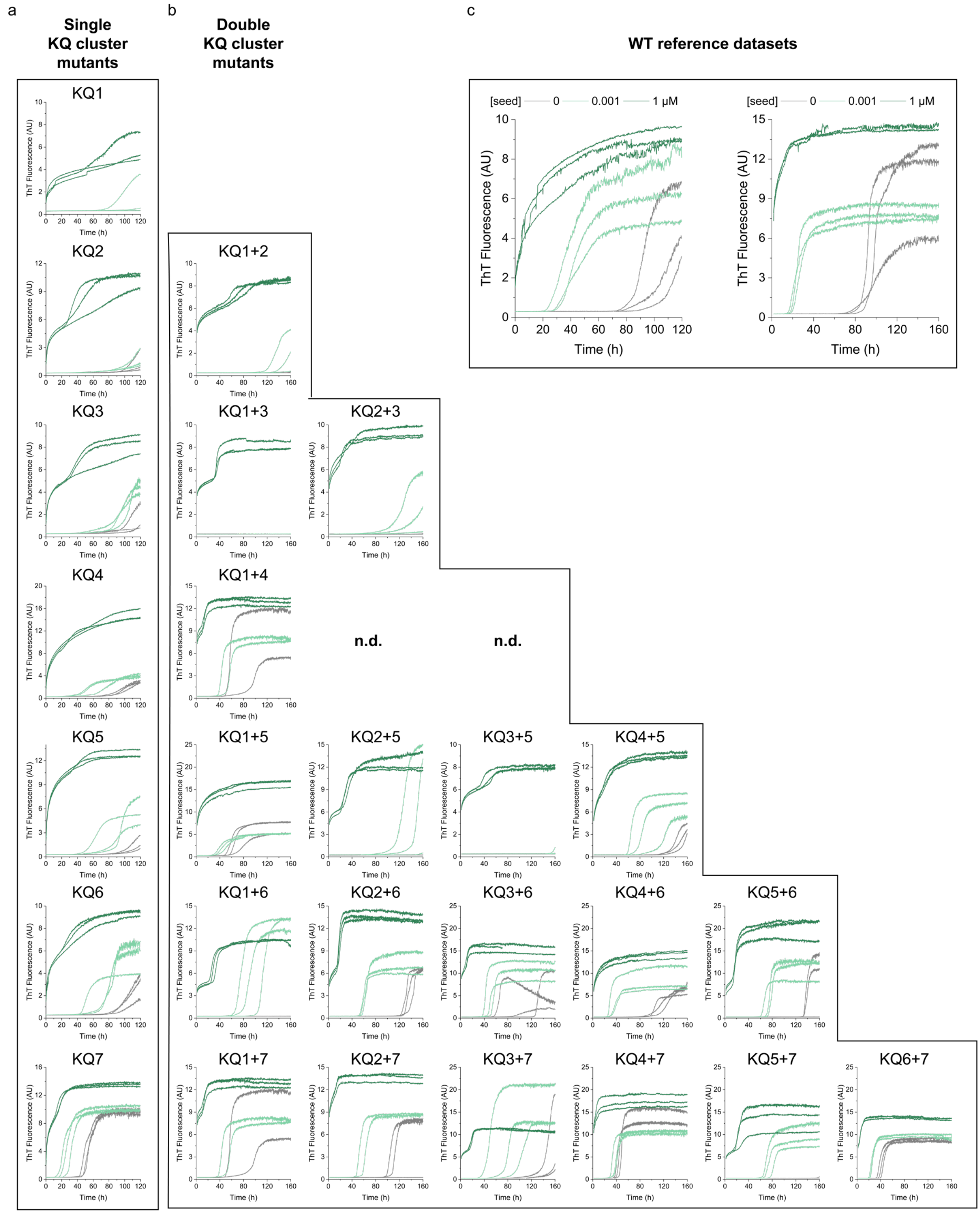


**Supplementary Figure 19. Amplification of Fm seed at low pH.** The data show raw data from two datasets with **(a)** single cluster and **(b)** double cluster variants. The corresponding WT references are shown in **c**. The concentration of seeds was 0 (grey), 1 nM (light green), and 1 µM (dark green).

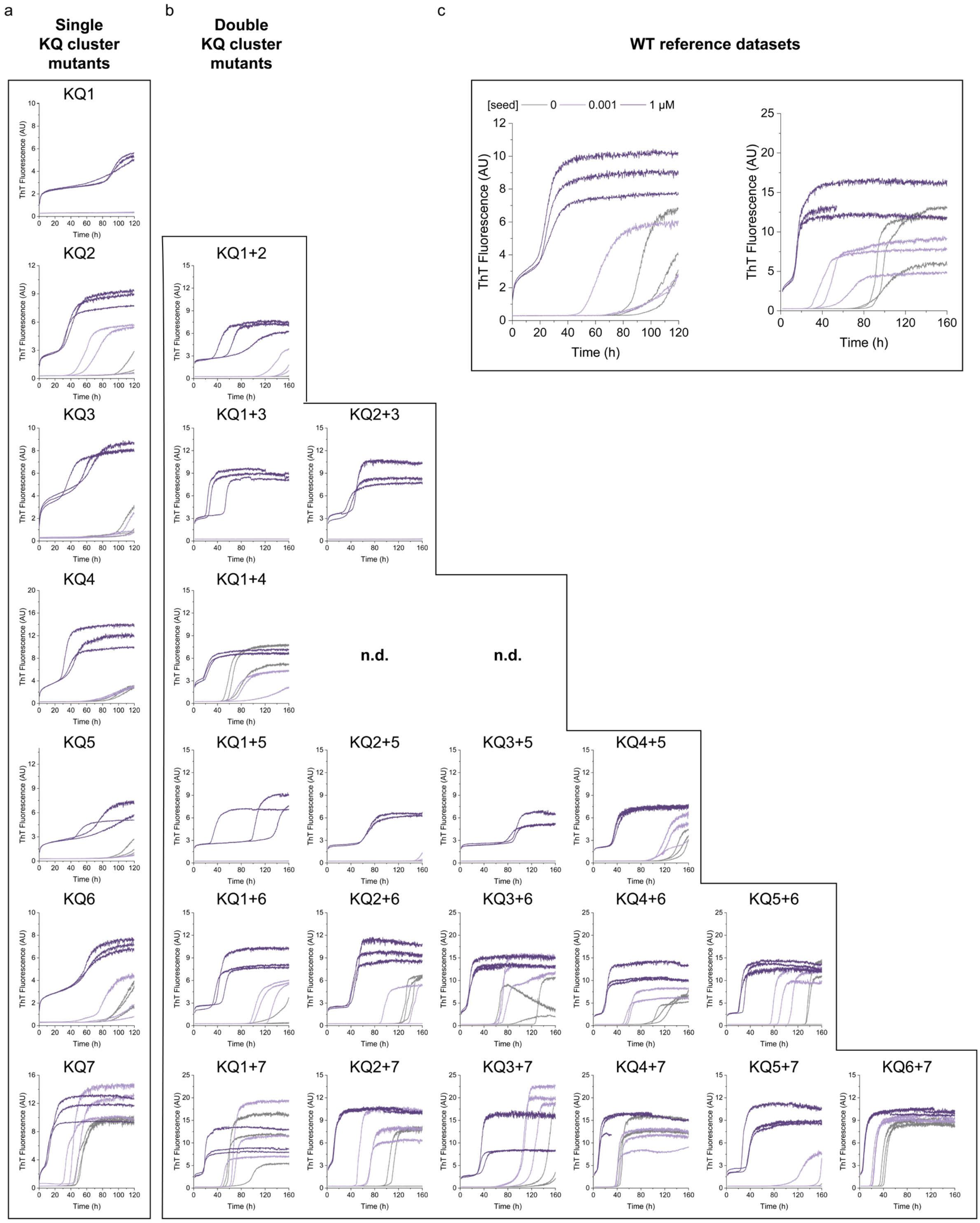


**Supplementary Figure 20. Amplification of Ri seed at low pH.** The data show raw data from two datasets with **(a)** single cluster and **(b)** double cluster variants. The corresponding WT references are shown in **c**. The concentration of seeds was 0 (grey), 1 nM (light purple), and 1 µM (dark purple).

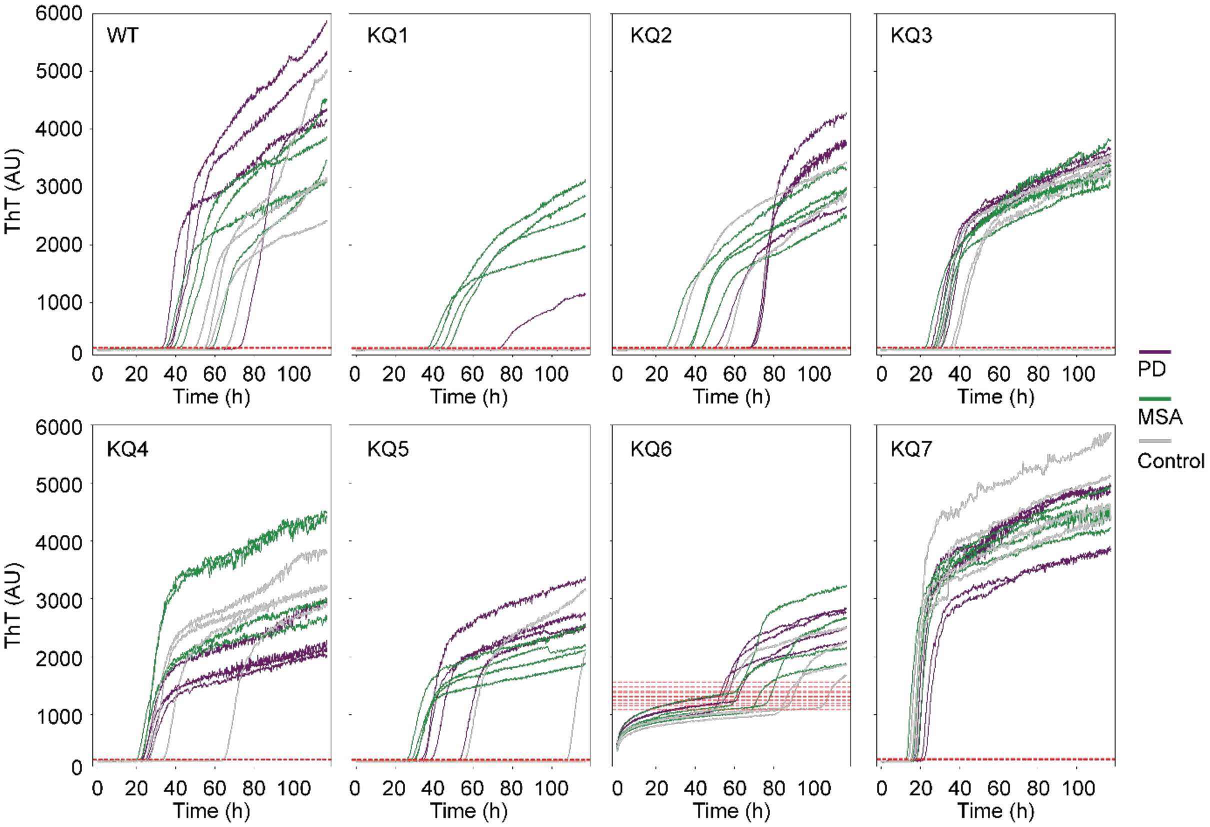


**Supplementary Figure 21. Raw data from seed amplification assay** with brain homogenates of patients with PD (purple), MSA (green), or healthy controls (grey). The threshold fluorescence used to derived time-till-threshold (TTT) are indicated in red dashed lines (**Supplementary Table 6**).

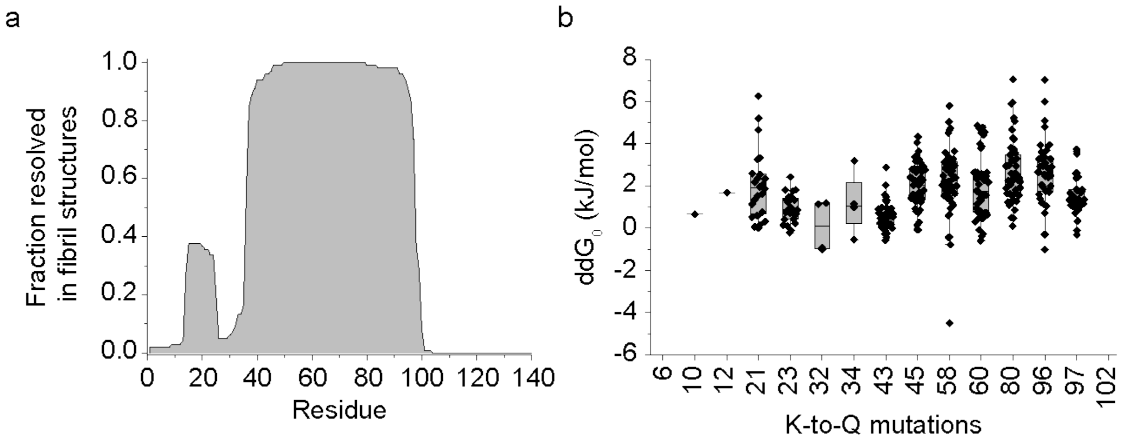


**Supplementary Figure 22. (a) Fraction of residues resolved in known αSyn fibril structures.** The data were taken from curated set of fibril structures from Amyloid Atlas.[1,2] **(b) *In silico* predictions of fibril stability changes** upon mutation of lysines to glutamine using FoldX.[3] The analysis was carried out on 47 representative fibril structures from Amyloid Atlas database (Supplementary table 9).

### Supplementary Tables

**Supplementary Table 1. Overview of the αSyn variants and assays used in this study.** The variant studied in a given assay is indicated by x. Conventional naming of single-point and double-point mutants is used throughout the manuscript (e.g., K6Q, K6Q+K10Q). Mutation K80Q is designated as variant KQ6 for consistency with the other cluster variants. # - number of mutations, [c] – de novo aggregation with varying protein concentration, I - – de novo aggregation with varying salt, pH - – de novo aggregation with varying pH, St – Fibril stability, A – AFM analysis of fibrils, E – elongation on WT fibrils, S – Seed amplification assay

| Variant name | # | K residues mutated to Q | Experimental assays | | | | | | |
|---|---|---|---|---|---|---|---|---|---|
| | | | [c] | I | pH | St | A | E | S |
| Single-point mutants | | | | | | | | | |
| | 1 | 6 | | | x | | | x | |
| | 1 | 10 | | | x | | | x | |
| | 1 | 12 | | | | | | x | |
| | 1 | 21 | | | | | | x | |
| | 1 | 23 | | | | | | x | |
| | 1 | 32 | | | | | | x | |
| | 1 | 34 | | | | | | x | |
| | 1 | 43 | | | | | | x | |
| | 1 | 45 | | | | | | x | |
| | 1 | 58 | | | | | | x | |
| | 1 | 60 | | | | | | x | |
| KQ6 | 1 | 80 | x | x | x | x | x | x | x |
| | 1 | 96 | | | x | | | x | |
| | 1 | 97 | | | | | | x | |
| | 1 | 102 | | | | | | x | |
| Double-point mutants (partial cluster variants) | | | | | | | | | |
| K6Q+ K10Q | 2 | 6+10 | | | x | | | | |
| K96Q+ K97Q | 2 | 96+97 | | | x | | | | |
| Single cluster mutant variants | | | | | | | | | |
| KQ1 | 3 | 6+10+12 | x | x | x | x | x | x | x |
| KQ2 | 2 | 21+23 | x | x | x | x | x | x | x |
| KQ3 | 2 | 32+34 | x | x | x | x | x | x | x |
| KQ4 | 2 | 43+45 | x | x | x | x | x | x | x |
| KQ5 | 2 | 58+60 | x | x | x | x | x | x | x |
| KQ7 | 3 | 96+97+102 | x | x | x | x | x | x | x |
| Double cluster mutant variants | | | | | | | | | |
| KQ1+2 | 5 | 6+10+12+21+23 | | | | | | x | x |
| KQ1+3 | 5 | 6+10+12+32+34 | | | | | | x | x |
| KQ1+4 | 5 | 6+10+12+43+45 | | | | | | x | x |
| KQ1+5 | 5 | 6+10+12+58+60 | | | | | | x | x |

| | | | | | |
|---|---|---|---|---|---|
| KQ1+6 | 4 | 6+10+12+80 | | x | x |
| KQ1+7 | 6 | 6+10+12+96+97+102 | | x | x |
| KQ2+3 | 4 | 21+23+32+34 | | x | x |
| KQ2+4 | 4 | 21+23+43+45 | | x | |
| KQ2+5 | 4 | 21+23+58+60 | | x | x |
| KQ2+6 | 3 | 21+23+80 | | | x |
| KQ2+7 | 5 | 21+23+96+97+102 | | x | x |
| KQ3+5 | 4 | 32+34+58+60 | | x | x |
| KQ3+6 | 3 | 32+34+80 | | x | x |
| KQ3+7 | 5 | 32+34+96+97+102 | | x | x |
| KQ4+5 | 4 | 43+45+58+60 | | x | x |
| KQ4+6 | 3 | 43+45+80 | | x | x |
| KQ4+7 | 5 | 43+45+96+97+102 | | x | x |
| KQ5+6 | 3 | 58+60+80 | | x | x |
| KQ5+7 | 5 | 58+60+96+97+102 | | x | x |
| KQ6+7 | 4 | 80+96+97+102 | | x | x |
| Triple cluster mutant variants | | | | | |
| KQ1+2+3 | 7 | 6+10+12+21+23+32+34 | x | | |
| KQ1+2+4 | 7 | 6+10+12+21+23+43+45 | x | | |
| KQ1+2+5 | 7 | 6+10+12+21+23+58+60 | x | | |
| KQ1+2+6 | 6 | 6+10+12+21+23+80 | x | | |
| KQ1+2+7 | 8 | 6+10+12+21+23+96+97+102 | x | | |
| KQ1+6+7 | 7 | 6+10+12+80+96+97+102 | x | | |
| KQ2+3+5 | 6 | 21+23+32+34+58+60 | x | | |
| KQ2+3+6 | 5 | 21+23+32+34+80 | x | | |
| KQ2+3+7 | 7 | 21+23+32+34+96+97+102 | x | | |
| KQ2+4+5 | 6 | 21+23+43+45+58+60 | x | | |
| KQ2+4+7 | 7 | 21+23+43+45+96+97+102 | x | | |
| KQ2+6+7 | 6 | 21+23+80+96+97+102 | x | | |
| KQ3+5+6 | 5 | 32+34+58+60+80 | x | | |
| KQ3+5+7 | 7 | 32+34+58+60+96+97+102 | x | | |
| KQ3+6+7 | 6 | 32+34+80+96+97+102 | x | | |
| KQ4+5+6 | 5 | 43+45+58+60+80 | x | | |
| KQ4+5+7 | 7 | 43+45+58+60+96+97+102 | x | | |
| KQ4+6+7 | 6 | 43+45+80+96+97+102 | x | | |
| KQ5+6+7 | 6 | 58+60+80+96+97+102 | x | | |

**Supplementary Table 2. Comparison of different parameterization of charge and ionic strength using weighted least square regression of log-transformed aggregation half-times** on full dataset. The values correspond to fitted parameters with error estimated obtained from the statsmodel module of python. AIC, BIC, df_resid, df_model correspond to Akaike information criterion, Bayesian information criterion, degrees of freedom of residuals, and degrees of freedom of model, respectively.

| **model** | $Q + \sqrt{I}$ | | | | $Q + Q\sqrt{I}$ | | | | $Q^2 + \sqrt{I}$ | | | | $Q^2 + Q^2\sqrt{I}$ | | | |
|---|---|---|---|---|---|---|---|---|---|---|---|---|---|---|---|---|
| **dataset** | full | | | | full | | | | full | | | | full | | | |
| **$R^2$** | 0.74 | | | | 0.73 | | | | 0.72 | | | | 0.71 | | | |
| **Adjusted $R^2$** | 0.71 | | | | 0.70 | | | | 0.68 | | | | 0.68 | | | |
| **AIC** | 592 | | | | 597 | | | | 613 | | | | 615 | | | |
| **BIC** | 694 | | | | 699 | | | | 715 | | | | 717 | | | |
| **df_resid** | 221 | | | | 221 | | | | 221 | | | | 221 | | | |
| **df_model** | 28 | | | | 28 | | | | 28 | | | | 28 | | | |
| **parameter** | **β ± SE** | | | **p** | **β ± SE** | | | **p** | **β ± SE** | | | **p** | **β ± SE** | | | **p** |
| **Intercept** | 2.85 | | 0.40 | 0.00 | 2.30 | | 0.39 | 0.00 | 4.65 | | 0.41 | 0.00 | 3.91 | ± | 0.37 | 0.00 |
| **logc** | -0.54 | ± | 0.08 | 0.00 | -0.54 | ± | 0.09 | 0.00 | -0.56 | ± | 0.09 | 0.00 | -0.56 | ± | 0.09 | 0.00 |
| **$\sqrt{I}$** | -1.71 | ± | 0.52 | 0.00 | | ± | | | -1.92 | ± | 0.55 | 0.00 | | ± | | |
| **Q** | -0.36 | ± | 0.02 | 0.00 | -0.39 | | 0.03 | 0.00 | | ± | | | | ± | | |
| **$Q\sqrt{I}$** | | ± | | | 0.11 | | 0.04 | 0.01 | | | | | | ± | | |
| **$Q^2$** | | | | | | | | | 0.02 | | 0.00 | 0.00 | 0.02 | ± | 0.00 | 0.00 |
| **$Q^2\sqrt{I}$** | | | | | | | | | | | | | -0.01 | ± | 0.00 | 0.00 |
| **KQ1** | 0.29 | ± | 0.20 | 0.14 | 0.35 | ± | 0.20 | 0.09 | 0.32 | ± | 0.21 | 0.13 | 0.36 | ± | 0.21 | 0.08 |
| **KQ2** | 0.28 | ± | 0.20 | 0.15 | 0.33 | ± | 0.20 | 0.10 | 0.28 | ± | 0.21 | 0.17 | 0.33 | ± | 0.21 | 0.11 |
| **KQ3** | 0.12 | ± | 0.22 | 0.58 | 0.16 | ± | 0.23 | 0.49 | 0.08 | ± | 0.23 | 0.72 | 0.12 | ± | 0.23 | 0.60 |
| **KQ4** | -0.36 | ± | 0.19 | 0.05 | -0.32 | ± | 0.19 | 0.09 | -0.28 | ± | 0.19 | 0.15 | -0.26 | ± | 0.19 | 0.19 |
| **KQ5** | -0.02 | ± | 0.18 | 0.91 | 0.02 | ± | 0.18 | 0.91 | 0.01 | ± | 0.19 | 0.97 | 0.04 | ± | 0.19 | 0.82 |
| **KQ6** | -0.63 | ± | 0.20 | 0.00 | -0.63 | ± | 0.20 | 0.00 | -0.51 | ± | 0.21 | 0.01 | -0.53 | ± | 0.21 | 0.01 |
| **KQ7** | 0.04 | ± | 0.23 | 0.87 | 0.09 | ± | 0.23 | 0.70 | 0.06 | ± | 0.24 | 0.80 | 0.10 | ± | 0.24 | 0.67 |
| **KQ1+2+3** | -1.98 | ± | 0.29 | 0.00 | -1.88 | ± | 0.29 | 0.00 | -2.26 | ± | 0.31 | 0.00 | -2.15 | ± | 0.30 | 0.00 |
| **KQ1+2+4** | -2.00 | ± | 0.38 | 0.00 | -1.89 | ± | 0.38 | 0.00 | -2.27 | ± | 0.40 | 0.00 | -2.11 | ± | 0.40 | 0.00 |
| **KQ1+2+5** | -1.70 | ± | 0.33 | 0.00 | -1.59 | ± | 0.33 | 0.00 | -2.05 | ± | 0.35 | 0.00 | -1.87 | ± | 0.34 | 0.00 |
| **KQ1+2+6** | -1.33 | ± | 0.31 | 0.00 | -1.24 | ± | 0.31 | 0.00 | -1.57 | ± | 0.33 | 0.00 | -1.42 | ± | 0.33 | 0.00 |
| **KQ1+2+7** | -1.91 | ± | 0.35 | 0.00 | -1.79 | ± | 0.35 | 0.00 | -2.27 | ± | 0.38 | 0.00 | -2.19 | ± | 0.37 | 0.00 |
| **KQ1+6+7** | -1.60 | ± | 0.34 | 0.00 | -1.50 | ± | 0.34 | 0.00 | -1.84 | ± | 0.36 | 0.00 | -1.73 | ± | 0.36 | 0.00 |
| **KQ2+3+5** | -1.00 | ± | 0.30 | 0.00 | -0.91 | ± | 0.30 | 0.00 | -1.19 | ± | 0.32 | 0.00 | -1.05 | ± | 0.32 | 0.00 |
| **KQ2+3+6** | -0.86 | ± | 0.28 | 0.00 | -0.79 | ± | 0.28 | 0.01 | -0.96 | ± | 0.30 | 0.00 | -0.84 | ± | 0.30 | 0.00 |
| **KQ2+3+7** | -1.69 | ± | 0.30 | 0.00 | -1.59 | ± | 0.30 | 0.00 | -1.87 | ± | 0.32 | 0.00 | -1.77 | ± | 0.31 | 0.00 |
| **KQ2+4+5** | -1.39 | ± | 0.29 | 0.00 | -1.30 | ± | 0.29 | 0.00 | -1.53 | ± | 0.30 | 0.00 | -1.40 | ± | 0.30 | 0.00 |
| **KQ2+4+7** | -1.55 | ± | 0.31 | 0.00 | -1.45 | ± | 0.31 | 0.00 | -1.82 | ± | 0.33 | 0.00 | -1.67 | ± | 0.33 | 0.00 |
| **KQ2+6+7** | -2.00 | ± | 0.32 | 0.00 | -1.91 | ± | 0.32 | 0.00 | -2.14 | ± | 0.33 | 0.00 | -2.05 | ± | 0.33 | 0.00 |
| **KQ3+5+6** | -1.45 | ± | 0.33 | 0.00 | -1.37 | ± | 0.34 | 0.00 | -1.51 | ± | 0.35 | 0.00 | -1.42 | ± | 0.35 | 0.00 |
| **KQ3+6+7** | -1.77 | ± | 0.28 | 0.00 | -1.68 | ± | 0.28 | 0.00 | -1.89 | ± | 0.29 | 0.00 | -1.77 | ± | 0.29 | 0.00 |
| **KQ4+5+6** | -1.96 | ± | 0.44 | 0.00 | -1.89 | ± | 0.45 | 0.00 | -2.04 | ± | 0.46 | 0.00 | -1.94 | ± | 0.46 | 0.00 |
| **KQ4+5+7** | -2.26 | ± | 0.58 | 0.00 | -2.15 | ± | 0.58 | 0.00 | -2.44 | ± | 0.60 | 0.00 | -2.30 | ± | 0.60 | 0.00 |
| **KQ4+6+7** | -2.78 | ± | 0.33 | 0.00 | -2.69 | ± | 0.33 | 0.00 | -2.98 | ± | 0.34 | 0.00 | -2.84 | ± | 0.34 | 0.00 |
| **KQ5+6+7** | -2.40 | ± | 0.33 | 0.00 | -2.31 | ± | 0.34 | 0.00 | -2.50 | ± | 0.35 | 0.00 | -2.41 | ± | 0.35 | 0.00 |

**Supplementary Table 3. Comparison of different parameterization of charge and ionic strength using weighted least square regression of log-transformed aggregation half-times** on dataset excluding the data from triple cluster variants. The values correspond to fitted parameters with error estimated obtained from the statsmodel module of python. AIC, BIC, df_resid, df_model correspond to Akaike information criterion, Bayesian information criterion, degrees of freedom of residuals, and degrees of freedom of model, respectively.

| **model** | $Q + \sqrt{I}$ | $Q + Q\sqrt{I}$ | $Q^2 + \sqrt{I}$ | $Q^2 + Q^2\sqrt{I}$ |
|---|---|---|---|---|
| **dataset** | single cluster only | single cluster only | single cluster only | single cluster only |
| **$R^2$** | 0.71 | 0.71 | 0.73 | 0.73 |
| **Adjusted $R^2$** | 0.69 | 0.69 | 0.72 | 0.71 |
| **AIC** | 427 | 429 | 411 | 413 |
| **BIC** | 462 | 464 | 446 | 448 |
| **df_resid** | 168 | 168 | 168 | 168 |
| **df_model** | 10 | 10 | 10 | 10 |

| **parameter** | **β** | **±** | **SD** | **p** | **β** | **±** | **SD** | **p** | **β** | **±** | **SD** | **p** | **β** | **±** | **SD** | **p** |
|---|---|---|---|---|---|---|---|---|---|---|---|---|---|---|---|---|
| **Intercept** | 2.79 | | 0.42 | 0.00 | 1.98 | | 0.40 | 0.00 | 4.10 | | 0.38 | 0.00 | 3.22 | ± | 0.35 | 0.00 |
| **logc** | -0.54 | ± | 0.08 | 0.00 | -0.54 | ± | 0.08 | 0.00 | -0.53 | ± | 0.08 | 0.00 | -0.53 | ± | 0.08 | 0.00 |
| **$\sqrt{I}$** | -2.55 | ± | 0.53 | 0.00 | | ± | | | -2.59 | ± | 0.51 | 0.00 | | ± | | |
| **Q** | -0.41 | ± | 0.03 | 0.00 | -0.47 | | 0.03 | 0.00 | | ± | | | | ± | | |
| **$Q\sqrt{I}$** | | ± | | | 0.23 | | 0.05 | 0.00 | | | | | | ± | | |
| **$Q^2$** | | | | | | | | | 0.03 | | 0.00 | 0.00 | 0.03 | ± | 0.00 | 0.00 |
| **$Q^2\sqrt{I}$** | | | | | | | | | | | | | -0.02 | ± | 0.00 | 0.00 |
| **KQ1** | 0.15 | ± | 0.20 | 0.45 | 0.22 | ± | 0.20 | 0.25 | -0.26 | ± | 0.20 | 0.20 | -0.12 | ± | 0.20 | 0.54 |
| **KQ2** | 0.19 | ± | 0.19 | 0.32 | 0.24 | ± | 0.19 | 0.20 | -0.02 | ± | 0.18 | 0.91 | 0.07 | ± | 0.18 | 0.71 |
| **KQ3** | 0.07 | ± | 0.21 | 0.73 | 0.12 | ± | 0.22 | 0.56 | -0.13 | ± | 0.21 | 0.53 | -0.04 | ± | 0.21 | 0.84 |
| **KQ4** | -0.48 | ± | 0.18 | 0.01 | -0.42 | ± | 0.18 | 0.02 | -0.70 | ± | 0.18 | 0.00 | -0.61 | ± | 0.18 | 0.00 |
| **KQ5** | -0.12 | ± | 0.18 | 0.50 | -0.07 | ± | 0.18 | 0.71 | -0.34 | ± | 0.17 | 0.05 | -0.25 | ± | 0.17 | 0.15 |
| **KQ6** | -0.67 | ± | 0.19 | 0.00 | -0.66 | ± | 0.19 | 0.00 | -0.73 | ± | 0.18 | 0.00 | -0.70 | ± | 0.18 | 0.00 |
| **KQ7** | -0.11 | ± | 0.23 | 0.64 | -0.03 | ± | 0.23 | 0.90 | -0.53 | ± | 0.22 | 0.02 | -0.39 | ± | 0.22 | 0.09 |

**Supplementary Table 4. Results of kinetic, thermodynamic, and morphological analyses of WT and KQ fibrils.** The values of relative changes of elongation barrier ($\Delta\Delta G^{\ddagger}$) between mutant and WT, fibril stability ($\Delta G$), and m-values correspond to means ± SEM (n=3-4). The values of AFM derived heights correspond to means ± SD (n = 11-118, **Figure 3**).

| | | Elongation | Stability | | AFM |
|---|---|---|---|---|---|
| **aSyn** | **K-Q mutated residues** | **$\Delta\Delta G^{\ddagger}$ (kJ/mol)** | **$\Delta G$ (kJ/mol)** | **m-value (kJ/M/mol)** | **Fibril height (nm)** |
| **WT** | - | - | -30.7 ± 1.6 | 6.4 ± 1.4 | 7.3 ± 0.8 |
| **KQ1** | 6+10+12 | -0.4 ± 0.9 | -28.3 ± 0.7 | 3.6 ± 0.2 | 5.4 ± 1.5 |
| **KQ2** | 21+23 | 1.9 ± 2.2 | -25.8 ± 1.1 | 4.8 ± 0.4 | 5.3 ± 1.4 |
| **KQ3** | 32+34 | -0.4 ± 0.4 | -29.5 ± 1.1 | 4.9 ± 0.3 | 5.1 ± 0.8 |
| **KQ4** | 43+45 | -8.5 ± 1.7 | -35.0 ± 1.8 | 7.1 ± 1.7 | 5.4 ± 1.3 |
| **KQ5** | 58+60 | -3.7 ± 1.5 | -32.7 ± 0.6 | 4.7 ± 1.1 | 5.1 ± 0.8 |
| **KQ6** | 80 | -8.2 ± 3.6 | -32.0 ± 0.8 | 5.9 ± 0.4 | 7.9 ± 1.5 |
| **KQ7** | 96+97+102 | 0.9 ± 3.4 | -27.4 ± 1.0 | 7.5 ± 1.8 | 7.5 ± 1.0 |

**Supplementary Table 5. Energy barriers (ΔΔG‡) and saturation constants (*K*e) of elongation on WT and truncated fibril polymorphs.** The parameters for Fm-WT polymorphs are means ± SEM from ThT and QCM measurements (n=2-9). Values for WT-Ri and aS1-125 correspond to values from a single ThT measurement ± error from the fitting.

| | | Seed polymorph | | | | | |
|---|---|---|---|---|---|---|---|
| | | WT- Fm | | WT - Ri | | aS1-125 | |
| aSyn | K-Q mutated residues | ΔΔG‡ (kJ/mol) | *K*e (μM) | ΔΔG‡ (kJ/mol) | *K*e (μM) | ΔΔG‡ (kJ/mol) | *K*e (μM) |
| KQ cluster variants | | | | | | | |
| **WT** | - | - | 53 ± 7 | - | 10 ± 1 | - | 51 ± 7 |
| **KQ1** | 6+10+12 | 3.2 ± 0.4 | 71 ± 17 | 3.2 ± 0.1 | 3 ± 2 | 2.9 ± 0.3 | 98 ± 47 |
| **KQ2** | 21+23 | 3.9 ± 0.8 | 52 ± 14 | 5.0 ± 0.2 | 4 ± 3 | 3.9 ± 0.2 | 52 ± 22 |
| **KQ3** | 32+34 | 3.5 ± 0.8 | 82 ± 2 | 2.7 ± 0.1 | 7 ± 3 | 1.9 ± 0.2 | 57 ± 22 |
| **KQ4** | 43+45 | 3.6 ± 2.0 | 24 ± 8 | 3.6 ± 0.1 | 0.1 ± 0.5 | 1.4 ± 0.2 | 46 ± 19 |
| **KQ5** | 58+60 | 3.6 ± 1.1 | 24 ± 4 | 3.4 ± 0.1 | 0.4 ± 0.9 | 4.5 ± 0.1 | 14 ± 3 |
| **KQ6** | 80 | 5.7 ± 1.8 | 28 ± 9 | 2.3 ± 0.1 | 1 ± 2 | 2.6 ± 0.1 | 6 ± 1 |
| **KQ7** | 96+97+102 | 1.9 ± 0.5 | 167 ± 38 | 1.3 ± 0.1 | 5 ± 2 | 1.3 ± 0.1 | 23 ± 5 |

**Supplementary Table 6. Changes of energy barriers of elongation (ΔΔG‡) of single-point mutant and double cluster variants on WT – Fm seed.** The values correspond to means ± SEM from n number of measurements.

| Single-point mutant variants | | | |
|---|---|---|---|
| **aSyn** | **K-Q mutated** | **ΔΔG‡ (kJ/mol)** | |
| K6Q | 6 | 5.8 ± 0.3 | n=2 |
| K10Q | 10 | 3.0 ± 1.1 | n=2 |
| K12Q | 12 | 3.4 ± 0.5 | n=3 |
| K21Q | 21 | 4.2 ± 0.7 | n=2 |
| K23Q | 23 | 4.7 | n=1 |
| K32Q | 32 | 3.4 ± 0.3 | n=3 |
| K34Q | 34 | 4.0 ± 0.8 | n=2 |
| K43Q | 43 | 3.1 ± 0.7 | n=3 |
| K45Q | 45 | 4.0 ± 0.5 | n=2 |
| K58Q | 58 | 5.0 ± 0.6 | n=3 |
| K60Q | 60 | 4.9 ± 0.6 | n=2 |
| K96Q | 96 | 5.7 ± 1.8 | n=2 |
| K97Q | 97 | 2.6 ± 1.2 | n=2 |
| K102Q | 102 | 3.6 ± 0.7 | n=3 |

| Double cluster mutant variants | | | |
|---|---|---|---|
| **aSyn** | **K-Q mutated** | **ΔΔG‡ (kJ/mol)** | |
| 1+2 | 6+10+12+21+23 | 10.6 ± 0.3 | n=2 |
| 1+3 | 6+10+12+32+34 | 7.2 ± 2.5 | n=2 |
| 1+4 | 6+10+12+43+45 | 6.3 ± 1.4 | n=2 |
| 1+5 | 6+10+12+58+60 | 9.8 ± 2.9 | n=2 |
| 1+6 | 6+10+12+80 | 5.0 ± 0.7 | n=2 |
| 1+7 | 6+10+12+96+97+102 | 8.9 ± 0.5 | n=2 |
| 2+3 | 21+23+34+35 | 6.0 ± 0.1 | n=2 |
| 2+4 | 21+23+43+45 | 5.0 | n=1 |
| 2+5 | 21+23+58+60 | 8.9 ± 2.1 | n=2 |
| 2+6 | 21+23+80 | 4.4 ± 2.0 | n=2 |
| 2+7 | 21+23+96+97+102 | 3.0 | n=1 |
| 3+5 | 34+35+58+60 | 4.7 | n=1 |
| 3+6 | 34+35+80 | 3.3 | n=1 |
| 3+7 | 34+35+96+97+102 | 4.2 ± 0.7 | n=2 |
| 4+5 | 43+45+58+60 | 4.6 ± 1.3 | n=2 |
| 4+6 | 43+45+80 | 1.8 ± 0.7 | n=2 |
| 4+7 | 43+45+96+97+102 | 3.5 ± 2.1 | n=2 |
| 5+6 | 58+60+80 | 6.3 ± 1.7 | n=2 |
| 5+7 | 58+60+96+97+102 | 3.5 ± 1.0 | n=2 |
| 6+7 | 80+96+97+102 | 2.3 ± 0.4 | n=2 |

**Supplementary Table 7. Half-time analysis of ThT kinetics at low pH in the presence of 1 nM WT-Fm or WT-Ri polymorph.** The half-times were obtained by fitting equation M1 to the raw kinetic data. Values correspond to the mean ± SD from a triplicate measurement. The relative half times were obtained as $t_{0.5}$ (mut)/$t_{0.5}$(WT) where $t_{0.5}$ is the aggregation half-time. n.d. – not determined.

| aSyn | WT - Fm | | WT - Ri | |
|---|---|---|---|---|
| | Half-time (h) | Relative half-time | Half-time (h) | Relative half-time |
| | | Dataset 1 | | |
| WT | 42.3 ± 2.7 | 1 | 99.7 ± 0.0 | 1 |
| KQ1 | 119.9 ± 20.5 | 2.8 | n.d. | |
| KQ2 | n.d. | | 65.7 ± 7.4 | 0.7 |
| KQ3 | 100.8 ± 5.2 | 2.4 | 107.3 ± 20.4 | 1.1 |
| KQ4 | 61.6 ± 12.3 | 1.5 | 99.3 ± 5.0 | 1 |
| KQ5 | 83.4 ± 13.9 | 2 | n.d. | |
| KQ6 | 72.8 ± 13.6 | 1.7 | 123.8 ± 31.0 | 1.2 |
| KQ7 | 28.5 ± 4.3 | 0.7 | 44.6 ± 6.5 | 0.4 |
| | | Dataset 2 | | |
| WT | 26.3 ± 0.0 | 1 | 52.8 ± 0.0 | 1 |
| KQ1+2 | 135.8 ± 0.0 | 5.2 | 158.3 ± 14.7 | 3 |
| KQ1+3 | n.d. | | n.d. | |
| KQ1+4 | 49.4 ± 4.7 | 1.9 | 80.2 ± 4.7 | 1.5 |
| KQ1+5 | 94.5 ± 15.4 | 3.6 | n.d. | |
| KQ1+6 | 79.7 ± 5.4 | 3 | 118.1 ± 12.1 | 2.2 |
| KQ1+7 | 66.1 ± 24.3 | 2.5 | 63.9 ± 8.4 | 1.2 |
| KQ2+3 | 145.9 ± 19.0 | 5.5 | n.d. | |
| KQ2+5 | 139.8 ± 12.3 | 5.3 | n.d. | |
| KQ2+6 | 62.6 ± 2.3 | 2.4 | 122.1 ± 24.1 | 2.3 |
| KQ2+7 | 62.0 ± 6.0 | 2.4 | 84.4 ± 14.9 | 1.6 |
| KQ3+5 | n.d. | | n.d. | |
| KQ3+6 | 50.4 ± 5.8 | 1.9 | 74.3 ± 4.7 | 1.4 |
| KQ3+7 | 79.0 ± 22.3 | 3 | 113.4 ± 9.0 | 2.1 |
| KQ4+5 | 91.3 ± 22.6 | 3.5 | 123.9 ± 2.0 | 2.3 |
| KQ4+6 | 37.0 ± 5.1 | 1.4 | 61.7 ± 2.6 | 1.2 |
| KQ4+7 | 39.1 ± 3.6 | 1.5 | 43.5 ± 2.8 | 0.8 |
| KQ5+6 | 74.5 ± 5.4 | 2.8 | 94.9 ± 12.6 | 1.8 |
| KQ5+7 | 77.8 ± 4.5 | 3 | 127.0 ± 0.0 | 2.4 |
| KQ6+7 | 26.9 ± 0.9 | 1 | 26.1 ± 1.5 | 0.5 |

**Supplementary Table 8. Analysis of seed amplification assay (SAA).** Times to threshold (TTT) in the presence homogenates from brains of patients with multiple system atrophy (MSA), Parkinson's disease (PD), or control (-) are provided as means ± SD (n=4). The mean values (n=4) of baseline and threshold ThT fluorescence use to derive the TTT are provided.

| aSyn | Seed | Time to Threshold (h) | Mean baseline (AU) | Mean Threshold (AU) |
|---|---|---|---|---|
| WT | - | 57 ± 6 | 184 | 223 |
| | MSA | 44 ± 10 | 191 | 223 |
| | PD | 45 ± 19 | 189 | 223 |
| KQ1 | - | 117 ± 0 | 187 | 221 |
| | MSA | 42 ± 5 | 186 | 216 |
| | PD | 106 ± 21 | 186 | 221 |
| KQ2 | - | 80 ±44 | 188 | 223 |
| | MSA | 36 ± 8 | 192 | 228 |
| | PD | 64 ± 9 | 188 | 221 |
| KQ3 | - | 55 ± 42 | 189 | 222 |
| | MSA | 27 ± 3 | 190 | 224 |
| | PD | 28 ± 2 | 190 | 228 |
| KQ4 | - | 38 ± 19 | 191 | 227 |
| | MSA | 22 ± 1 | 190 | 224 |
| | PD | 25 ± 2 | 191 | 223 |
| KQ5 | - | 100 ± 29 | 189 | 219 |
| | MSA | 29 ± 2 | 188 | 223 |
| | PD | 40 ± 9 | 188 | 226 |
| KQ6 | - | 84 ± 22 | 625 | 1218 |
| | MSA | 69 ± 8 | 675 | 1338 |
| | PD | 58 ± 5 | 721 | 1361 |
| KQ7 | - | 16 ± 1 | 187 | 222 |
| | MSA | 16 ± 2 | 185 | 220 |
| | PD | 20 ± 2 | 191 | 227 |

**Supplementary Table 9. In-silico analysis of fibril stability changes upon K-to-Q mutation using FoldX.**

| | K-to-Q ddG (kJ/mol) | | | | | | | | | | | | | | |
|---|---|---|---|---|---|---|---|---|---|---|---|---|---|---|---|
| PDB code | **6** | **10** | **12** | **21** | **23** | **32** | **34** | **43** | **45** | **58** | **60** | **80** | **96** | **97** | **102** |
| **2n0a** | | | | | | 1.2 | -0.5 | 0.0 | -0.1 | 5.8 | 1.3 | 2.3 | 1.8 | 1.8 | |
| **6cu7** | | | | | | | | 0.4 | 2.2 | 0.7 | 0.8 | 5.2 | 2.5 | 1.4 | |
| **6cu8** | | | | | | | | -0.6 | 0.7 | 2.1 | 0.8 | 1.3 | | | |
| **6rt0** | | | | 2.5 | 1.0 | | | 1.0 | 2.8 | 1.4 | 1.4 | 1.5 | 1.1 | 0.9 | |
| **6rtb** | | 0.7 | 1.7 | 0.2 | 1.4 | | | 0.3 | 2.0 | | | 1.6 | | | |
| **6sst** | | | | 5.2 | 0.7 | | | 0.1 | 2.7 | -0.4 | 3.9 | 1.3 | 1.1 | | |
| **6ssx** | | | | 0.6 | 0.7 | | | 0.0 | 2.8 | 2.2 | 4.5 | 1.8 | 2.5 | | |
| **6xyo** | | | | 0.8 | 0.9 | 1.1 | 3.2 | 0.2 | 2.3 | 5.0 | 4.0 | 6.0 | 3.1 | 1.8 | |
| **6xyp** | | | | 2.3 | 0.4 | -1.0 | 1.2 | 0.3 | 0.1 | 3.0 | 4.8 | 4.3 | 3.4 | 2.6 | |
| **6xyq** | | | | 2.1 | -0.1 | -0.9 | 1.0 | 1.5 | 0.3 | 3.4 | 4.8 | 3.2 | 3.7 | 1.2 | |
| **7nca** | | | | | | | | 0.2 | 0.8 | 2.6 | 1.5 | 0.5 | 2.0 | 1.3 | |
| **7ncg** | | | | | | | | 0.4 | 3.2 | 4.7 | 2.6 | 1.6 | 3.2 | 1.3 | |
| **7nch** | | | | 2.2 | 0.1 | | | 0.8 | 1.4 | 0.6 | 2.5 | 0.1 | | | |
| **7nci** | | | | 2.0 | 0.2 | | | 0.6 | 1.4 | 1.7 | 2.5 | 0.9 | | | |
| **7ncj** | | | | 2.3 | -0.2 | | | 0.7 | 2.4 | 1.9 | 2.1 | 0.5 | 3.9 | 1.4 | |
| **7nck** | | | | | | | | 0.1 | 1.7 | 4.5 | 1.7 | 7.1 | 3.6 | 2.4 | |
| **7ozg** | | | | 4.7 | 0.8 | | | 0.6 | 1.3 | -0.4 | 2.3 | 1.3 | -1.0 | | |
| **7ozh** | | | | | | | | 1.4 | 2.8 | 2.1 | 1.5 | 1.1 | | | |
| **7v47** | | | | | | | | 0.0 | 3.5 | 2.7 | 1.1 | 2.3 | 2.9 | | |
| **7v48** | | | | | | | | 1.3 | 1.5 | 4.7 | 0.5 | 2.6 | 3.0 | 2.4 | |
| **7v49** | | | | | | | | 1.6 | 0.9 | -4.5 | 1.6 | 4.7 | | | |
| **7xjx** | | | | | | | | 0.6 | 3.7 | 3.2 | 1.8 | 5.1 | 1.8 | 3.5 | |
| **7xo0** | | | | | | | | 1.3 | 1.3 | 2.8 | -0.4 | 3.2 | 1.2 | 1.1 | |
| **7xo1** | | | | | | | | 1.0 | 2.7 | 2.4 | -0.6 | 2.6 | 3.9 | 1.0 | |
| **7xo2** | | | | | | | | 0.8 | 1.2 | 2.8 | 4.4 | 3.0 | 3.9 | 1.0 | |
| **7xo3** | | | | | | | | 1.6 | 2.0 | 3.0 | 0.5 | 2.1 | 2.4 | 3.7 | |
| **7yk2** | | | | | | | | -0.1 | 3.2 | 3.3 | 2.6 | 3.7 | 2.0 | 3.6 | |
| **7yk8** | | | | 0.0 | 0.7 | | | 1.0 | 4.3 | 3.1 | 2.3 | 4.2 | 0.7 | 1.0 | |
| **7yng** | | | | | | | | -0.4 | 1.4 | 3.2 | 1.6 | 3.8 | 1.7 | 0.3 | |
| **7ynr** | | | | | | | | 0.6 | 2.3 | 3.3 | 2.4 | 3.6 | 1.4 | 1.7 | |
| **7yns** | | | | | | | | 2.9 | 1.4 | 2.2 | 0.6 | 3.5 | 2.6 | -0.1 | |
| **7ynt** | | | | | | | | 0.5 | 2.8 | 3.6 | 2.5 | 2.6 | 2.9 | -0.3 | |
| **8cyr** | | | | 0.1 | 1.5 | | | 0.2 | 2.9 | 1.4 | 0.8 | 1.6 | 7.1 | 1.6 | |
| **8cys** | | | | 0.3 | 2.4 | | | 0.5 | 3.2 | 1.2 | 0.7 | 3.0 | 2.5 | 1.7 | |
| **8cyt** | | | | 3.3 | 1.5 | | | -0.6 | 2.4 | 1.9 | 4.9 | 2.3 | 6.0 | 1.1 | |
| **8cyv** | | | | 0.5 | 1.2 | | | -0.3 | 1.6 | 2.4 | 2.6 | 2.3 | 2.7 | 0.7 | |
| **8cyw** | | | | 6.3 | 0.4 | | | 0.9 | 1.7 | 2.5 | 3.8 | 1.2 | -0.3 | | |
| **8cyx** | | | | 1.5 | 0.9 | | | 0.2 | 1.6 | -0.8 | 0.9 | 1.6 | -0.3 | | |
| **8cyy** | | | | 1.6 | 1.8 | | | 0.0 | 3.4 | 3.6 | 2.6 | 2.4 | 5.1 | 1.8 | |
| **8cz0** | | | | 1.3 | 1.3 | | | 0.7 | 4.0 | 0.9 | 2.6 | 3.3 | 4.8 | 1.7 | |
| **8cz1** | | | | 2.6 | 1.3 | | | 0.7 | 1.8 | 1.1 | 0.7 | 3.0 | 2.9 | 1.2 | |
| **8cz2** | | | | 1.9 | 0.2 | | | 0.7 | 3.1 | 1.8 | 0.3 | 1.7 | 2.6 | 1.4 | |
| **8cz3** | | | | 3.3 | 1.8 | | | 0.5 | -0.1 | 3.1 | 4.6 | 1.9 | 3.3 | 0.9 | |
| **8cz6** | | | | 1.1 | 1.8 | | | -0.3 | 2.8 | 1.6 | -0.3 | 2.2 | 2.0 | 1.4 | |
| **8h03** | | | | | | | | 0.9 | 0.9 | 2.0 | 0.6 | 1.9 | 1.6 | | |
| **8h04** | | | | | | | | 0.7 | 2.4 | 2.0 | -0.1 | 2.9 | 3.3 | 2.6 | |
| **8h05** | | | | | | | | 2.0 | 1.4 | 1.5 | 2.2 | 5.9 | | | |

| | | | | | | | | | | | | | | | |
|---|---|---|---|---|---|---|---|---|---|---|---|---|---|---|---|
| **n - resolved** | 0 | 1 | 1 | 24 | 24 | 4 | 4 | 47 | 47 | 46 | 46 | 47 | 40 | 33 | 0 |
| **% - resolved** | 0.0 | 0.0 | 0.0 | 0.5 | 0.5 | 0.1 | 0.1 | 1.0 | 1.0 | 1.0 | 1.0 | 1.0 | 0.9 | 0.7 | 0.0 |
| **mean** | | 0.7 | 1.7 | 2.0 | 0.9 | 0.1 | 1.2 | 0.6 | 2.0 | 2.2 | 2.0 | 2.7 | 2.6 | 1.5 | |
| **SD** | 0.0 | 0.0 | 0.0 | 1.6 | 0.7 | 1.1 | 1.3 | 0.7 | 1.1 | 1.7 | 1.5 | 1.5 | 1.6 | 0.9 | 0.0 |
